# EternaBrain: Automated RNA design through move sets and strategies from an Internet-scale RNA videogame

**DOI:** 10.1101/326736

**Authors:** Rohan V. Koodli, Benjamin Keep, Katherine R. Coppess, Fernando Portela, Eterna participants, Rhiju Das

## Abstract

Emerging RNA-based approaches to disease detection and gene therapy require RNA sequences that fold into specific base-pairing patterns, but computational algorithms generally remain inadequate for these secondary structure design tasks. The Eterna project has crowdsourced RNA design to human video game players in the form of puzzles that reach extraordinary difficulty. Here, we demonstrate that Eterna participants’ moves and strategies can be leveraged to improve automated computational RNA design. We present an eternamoves-large repository consisting of 1.8 million of player moves on 12 of the most-played Eterna puzzles as well as an eternamoves-select repository of 30,477 moves from the top 72 players on a select set of more advanced puzzles. On eternamoves-select, we present a multilayer convolutional neural network (CNN) EternaBrain that achieves test accuracies of 51% and 34% in base prediction and location prediction, respectively, suggesting that top players’ moves are partially stereotyped. Pipelining this CNN’s move predictions with single-action-playout (SAP) of six strategies compiled by human players solves 61 out of 100 independent puzzles in the Eterna100 benchmark. EternaBrain-SAP outperforms previously published RNA design algorithms and achieves similar or better performance than a newer generation of deep learning methods, while being largely orthogonal to these other methods. Our study provides useful lessons for future efforts to achieve human-competitive performance with automated RNA design algorithms.

## Introduction

Due to its versatility and important roles throughout biology, there is strong interest in designing RNA-guided machines for disease detection, virus defense, and gene therapy, e.g., for gene silencing and CRISPR/Cas9 gene editing (1, 2). Much of RNA’s functionality is dependent on well-defined structures, and so these and future RNA technologies require computational methods to effectively design sequences that fold into a target structure or set of structures suited to a desired task. The simplest problem involves designing RNA sequences that energetically favor one specific secondary structure – a target pattern of Watson-Crick base pairs – over alternative secondary structures. Even this most basic problem is computationally difficult (3). While exact solutions can be determined through exhaustive calculation (4), such computational enumeration generally takes an impractically long time for solve complex target structures.

Numerous groups have developed RNA secondary structure design algorithms, including MODENA (5), RNAinverse (6), INFO-RNA (7), RNA-SSD (8), and NUPACK (9). In the original studies presenting these methods, tests typically involved simple structures that do not capture the symmetries, duplex lengths, and sizes needed for biotechnology applications (10). The incompleteness of these prior methods and tests became clear with the release of the *Eterna* game (11) in 2011, which crowdsources RNA design in the form of puzzles through an internet-scale videogame (Figure 1A-C). Eterna participants learn the basics of RNA design through examples that are initially tested *in silico* through a computational model of secondary structure folding. Advanced participants can submit their RNA designs in lab challenges and receive wet-lab feedback on how their molecules fold during *in vitro* experiments performed on a weekly time scale. These efforts expose the ‘reality gap’ in RNA design – the mismatch between current computational folding models and experiment. In preparation for these experimental challenges, participants also challenge fellow participants through *in silico* puzzles that require learning or developing sophisticated puzzle-solving strategies; these separate ‘games within the game’ are useful for guiding the development of computational RNA design methods (10). Since Eterna’s inception, the community has grown to over 250,000 registered participants as of 2019, with over 17,000 player-created puzzles. This community has been successful in designing RNAs that consistently outperform RNA design algorithms in both *in silico* and *in vitro* tests (10).

**Figure 1.**
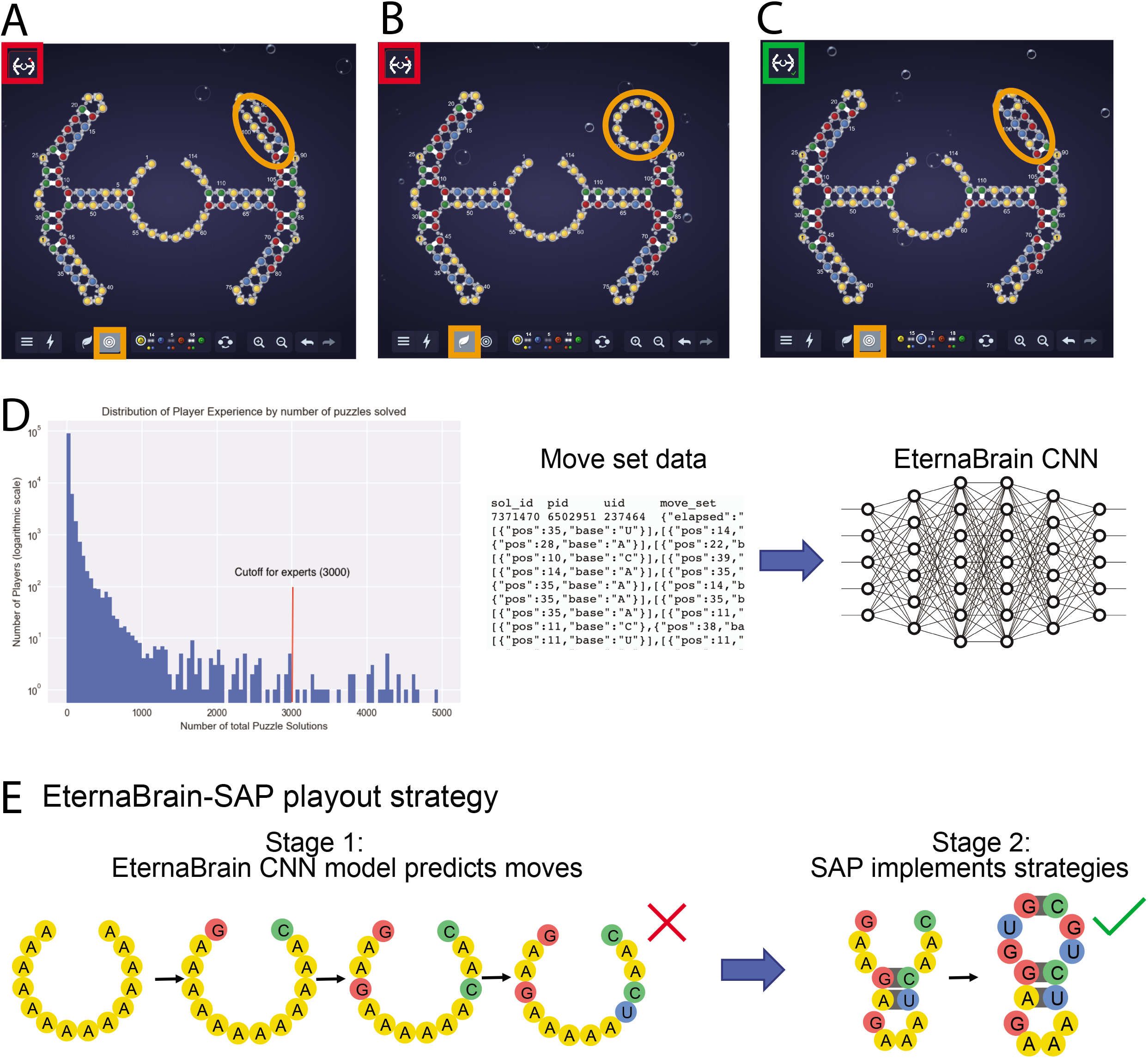
Eterna and EternaBrain. (A-C) Puzzle-solving interface presented to human players of Eterna including the state of the puzzle (whether it is solved or not) in the top left corner (red/green outline), the puzzle itself (in the middle), and the toolbar (bottom) with which the players can mutate the RNA sequence to make it fold into the desired state; yellow, blue, red, and green symbols represent A, U, G, and C nucleotides. (A) The desired target structure for the RNA molecule, as indicated by the bullseye in the bottom left (orange highlight). (B) Nature mode, as indicated by the leaf in the bottom left (orange highlight), gives the predicted minimum free energy structure for the current sequence. Since the bases in the top right should be paired with each other (orange circle), this puzzle is not yet folding correctly; this status is shown by the red indicator in the top left corner. (C) The solved puzzle. The nature-mode structure matches the target structure, and the indicator in the top left corner turns green, meaning the puzzle has been solved. (D) (left) Numerous players of contributed Eterna solutions. For preparing the *eternamoves-select* data set, we selected any user who had solved more than 3000 distinct puzzles, which left us with 72 players. (right) In EternaBrain, we tested whether information on players’ moves could be used to train a convolutional neural network. (E) For solving new puzzles, the final EternaBrain-SAP framework first uses the EternaBrain convolutional neural net model to predict sequence changes (‘moves’) for new RNA puzzles. In a second stage, the Single Action Playout (SAP), six additional hand-coded strategies are applied to complete the solution.

Since Eterna’s inception, participants have discovered classes of RNA secondary structures for which prior algorithms cannot find sequence solutions even *in silico*, i.e., when folded with computational energy models that can be rapidly evaluated (10). However, under the same computational energy models, solutions to these design problems can be discovered by experienced human participants. Thus, there remains a gap between algorithms and humans even for the purely computational problem of *in silico* design. Fortunately, Eterna has produced rich resources that might allow for this gap to be closed. First, in 2016, several participants curated a benchmark of 100 problems of increasing difficulty, termed the ‘Eterna100’ (10). By offering a wider spectrum of difficulty than prior benchmark sets based on inferred structures of random or biological RNAs (5–9), these secondary structures allow stringent tests of advances in computational design methods while still guaranteeing that solutions can be found. Second, player-created tutorial puzzles have ‘canonized’ new strategies for *in silico* RNA design. In 2016, Eterna developers installed these puzzles as the standard progression of problems for new participants. At the same time, Eterna participants have agreed to share the sequences of moves that lead to successful solutions to scientific research, resulting in a data repository including nearly 2 million player moves (Figure 1D).

The availability of such a large collection of move sets as well as canonical player strategies suggests new approaches to solving the RNA secondary structure design problem. While previous RNA design algorithms have used hierarchical decomposition of target structures, genetic algorithms, and probabilistic sampling of sequences (5,7,8), a recent generation of classification, translation, and game-playing algorithms make powerful use of statistical pattern recognition through multi-layer artificial neural networks (12). One striking example is Google DeepMind’s AlphaGo (13), which outcompeted expert human participants in the complex game of *Go*. This approach was inspired by the discovery that expert participant moves in the game Go were sufficiently stereotyped that they could be predicted with better than 50% accuracy by a convolutional neural network (CNN), despite the large space of possible moves in this game (up to 19 x 19 = 361 board positions for pieces, giving a baseline random guess accuracy of less than 0.5%) (13). AlphaGo and its successor AlphaGo Zero have further improved their performance using reinforcement learning (14). While reinforcement learning has shown promise for RNA design, it is not yet human competitive (15, 16). This observation suggests that there remain lessons to be learned from human moves and strategies. Nevertheless, it remains unclear whether single base changes during human RNA design are sufficiently stereotyped as to inform neural network or other machine learning methods – strategies that involve back-and-forth flipping of base changes, sets of multiple sequence moves, or sequence explorations outside the game browser may be important for the success of the best Eterna participants.

To test whether new strategies for automated RNA design might be gleaned from successful single base moves, we present a data set of 1.8 million moves on Eterna’s most played puzzles, called *eternamoves-large*, appropriately cleaned and labeled for machine learning applications. We conduct tests of CNNs to predict these moves given the game state, and report generally poor results. However, we do find that a set of 30,477 moves made by the most experienced participants on more difficult puzzles (*eternamoves-select*) are sufficiently stereotyped to allow training of an automated neural network EternaBrain move predictor with accuracy well above random baseline. We then challenge the resulting predictor to go beyond simply predicting individual moves and to instead perform a series of moves to solve novel RNA secondary structure design problems from scratch (as evaluated by *in silico* folding models). We find generally modest performance from this CNN-only approach, with results poorer than previously published algorithms. We then collate several of the participants’ hand-crafted strategies and pipeline a single-action-playout (SAP) of these strategies to the EternaBrain neural network approach. We show that the resulting EternaBrain-SAP algorithm (Figure 1E) achieves excellent performance on the Eterna100 benchmark by completing 61 of 100 puzzles, exceeding the performance of methods published prior to our work. Furthermore, EternaBrain-SAP performs similarly to newer methods developed concomitantly by us and other groups using complementary algorithmic approaches. In the discussion, we compare EternaBrain-SAP’s performance on the Eterna100 with these more recently-reported methods, including SIMARD (17), SentRNA (18), NEMO (19), antaRNA (20), MCTS-RNA (21), and the reinforcement learning methods of Eastman et al. (15) and LEARNA (16), drawing lessons for future efforts in automated RNA design. Additionally, we highlight the likelihood of future progress as other methods (4, 22) and newer computational energy functions (23) are developed and tested on the same benchmark as well as on biological RNA structures.

## Results

### Initial Training on 1.8 Million Moves

We tested several different neural network architectures and training sets for EternaBrain. We chose to use convolutional neural network (CNN) architectures, because of their success in other areas of game playing and machine learning (13,14) and also because of their expected ability to capture patterns visually recognized by humans during actual Eterna gameplay. A CNN (24) is a specific type of neural network that hierarchically builds up complex features from simpler features that neighbor each other. CNNs consist of convolutional and pooling layers which retrieve subsections of the input features and make their own larger-scale features. A CNN is a natural choice for tackling this problem because it mimics how human participants approach Eterna puzzles; that is, by learning features of the data from spatial information such as those seen in the puzzle interface and in solution browsers. Our work involves supervised learning: the CNN’s features and relationships were randomly initialized and then iteratively trained on large input datasets (training sets) and assessed for their predictive power when applied to previously unseen datasets (test sets).

We first trained the model on a large data set of 1.8 million moves spanning 12 representative puzzles from the Eterna progression (Supplemental Figure S1), which we call the *eternamoves-large* data set. The information in player move set data includes nucleotide sequence, RNA secondary structure (in ‘dot-bracket’ notation widely used in the RNA literature (25)), and predicted minimum Gibbs free energy of folding. These data on the game state are passed as input features to the CNN (Table 1). The location of the player’s chosen nucleotide change and the nucleotide identity itself were passed as labels to the CNN – they are the output to be predicted (Table 1). We chose to train one ‘base predictor’ CNN (BP-CNN) to predict the identity of the base change (e.g. A, U, C, or G) made by a player provided the location of the change; as well as a separate ‘location predictor’ CNN (LP-CNN) to predict the location of the change. For the BP-CNN, we did not require that the neural network change the base nor that it maintain Watson-Crick pairing at nucleotides that were supposed to be paired in the target structure. Doing so provided a means to assess if the neural network could independently derive these fundamental move requirements. For cross-validation, we randomly split the data into training and test sets (see Methods). Training took three days using 4 NVidia Titan X GPUs.

**Table 1.**
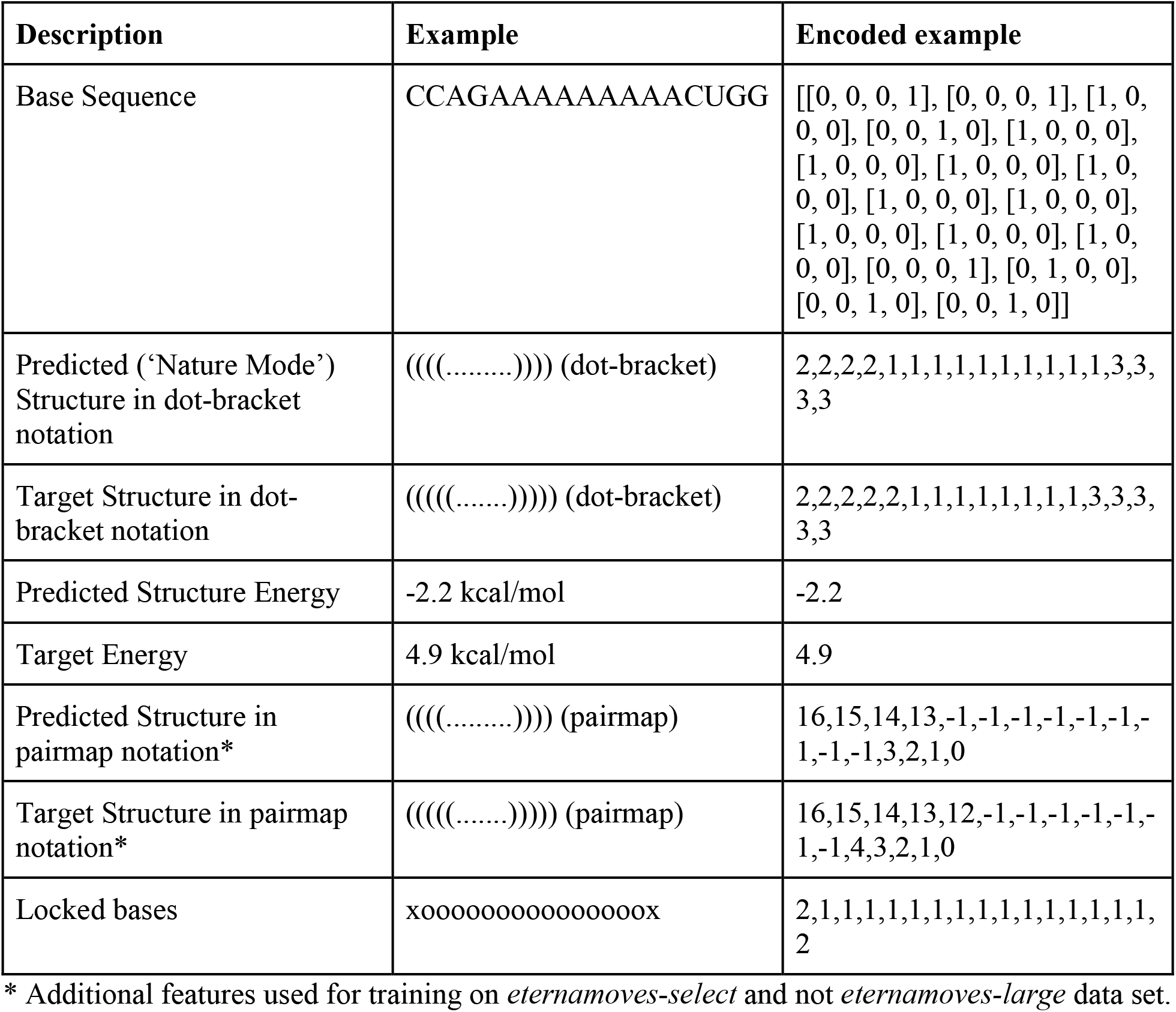
Input features used for training and testing EternaBrain convolutional neural network.

Random guessing of moves would give accuracies of 0.33 for the BP-CNN (if the base is forced to change from its starting identity to one of the three other bases) and 0.019 for the LP-CNN (based on the average inverse length of the puzzles). Our best training attempts achieved move accuracies for the BP-CNN and LP-CNN of 0.50 and 0.10, respectively on the training dataset-both higher than random guessing. However, when applied to the test dataset, the BP-CNN and LP-CNN achieved move accuracies of 0.34 and 0.021, respectively, which are not significantly better than random guessing. Furthermore, we discovered poor results in subsequent tests of this model applied to full puzzle playouts. In these playouts, we had the CNN predict both the identity and location of the move starting from its initial sequence. Then, we updated the puzzle state according to ViennaRNA energy calculations and repeated this process. The model was unable to solve the easiest puzzle on the Eterna100 - *Simple Hairpin* - even with modifications to allow stochastic choice of moves after ranking by the CNN (see Methods).

The low test accuracies and problems in an actual puzzle playout suggested that the CNNs might be overfitting their parameters to the training data and not generalizing well to Eterna puzzles that were outside the dataset. To counteract this overfitting, we had the networks randomly exclude randomly chosen nodes in each of their layers (dropout) and thus reduce the chance of overfitting. However, increasing this dropout rate gave only marginal improvements in test accuracy, suggesting that there was high variance in the training data (26). This could result if, in these 12 early Eterna puzzles, players were using a highly heterogeneous set of strategies or clicking randomly without a clear strategy. Other parameter changes to the network (e.g.. use of a single CNN to predict base and location; change in the type of neural network; number of layers; activation functions; see Supplemental Table S1) also did not significantly improve test accuracy. These results suggested that moves made by players across this entire set are not sufficiently similar to allow their automation, at least with the convolutional neural network architectures tested.

### Training on Selected Subsets of Players and Puzzles

We hypothesized that training on a select number of advanced Eterna participants would minimize the variance in the dataset. To select these ‘expert’ Eterna participants, we identified players who had solved at least 3000 Eterna puzzles. The resulting dataset contained moves from 72 players (Figure 1D; Supplemental Table S2). In addition, instead of using the 12 introductory simple puzzles used for *eternamoves-large* data set described above, we chose to collate moves from 78 puzzles compiled during a revision of the game’s main puzzle progression (Supplemental Figure S2). These puzzles were selected because they introduced key strategies used by Eterna players to solve many of the RNA design challenges on Eterna. The resulting dataset, dubbed *eternamoves-select*, was comprised of 30,447 moves. In addition to narrowing the training data to expert players and specialized puzzles, we added a more explicit representation of RNA secondary structure by introducing pairmaps. Rather than merely providing dot-bracket notation to the CNN, a structure pairmap uses indices of a list to explicitly show which bases are paired to other bases. The format of the pairmap is shown in Table 1.

Combining *eternamoves-select* and introducing pairmaps increased the predictive power of our CNNs. The final move accuracies for cross-validation were 0.51 for the BP-CNN and 0.34 for the LP-CNN (Supplemental Figures S3A and S3B; Table 2) when applied to the test dataset of *eternamoves-select*. These move accuracies were higher than the expected random baselines for these puzzles of 0.33 and 0.031, respectively. We expected that the model’s predictive powers would negatively correlate with the puzzle length, and this trend was indeed apparent. For example, the LP-CNN accuracy dropped from 0.43 to 0.28 when tested on puzzles with lengths of up to 50 nucleotides (nts), compared to puzzles with lengths from 101 to 150 nts; Table 3). The BP-CNN accuracy reduced from 0.57 to 0.47 with the same increase in puzzle length (Table 3). We also expected that the model would have higher prediction accuracy at nucleotides that involved paired regions of the target structure. In those regions, the model can correct ‘mismatches’ – nucleotides that should be paired but are not A-U, G-C, or G-U – and such moves would provide obvious location and base choices. Surprisingly, our model exhibited slightly better performance in both base and location prediction in regions of the puzzle that were unpaired rather than paired in the target structure (0.62 vs. 0.49, base; 0.38 vs. 0.33, location; Table 3).

**Table 2.** EternaBrain CNN accuracies on *eternamoves-select* with different splits of training and test sets.

| **Model** | **Training Accuracy** | **Test Accuracy** |
| --- | --- | --- |
| EternaBrain - base | 0.71 | 0.51 |
| EternaBrain - location | 0.31 | 0.34 |
| Half experts - base | 0.66 | 0.38 |
| Half experts - location | 0.30 | 0.11 |
| Half puzzles - base | 0.70 | 0.27 |
| Half puzzles - location | 0.33 | 0.02 |
| One expert - base | 0.79 | 0.25 |
| One expert - location | 0.5 | 0.01 |

**Table 3.**
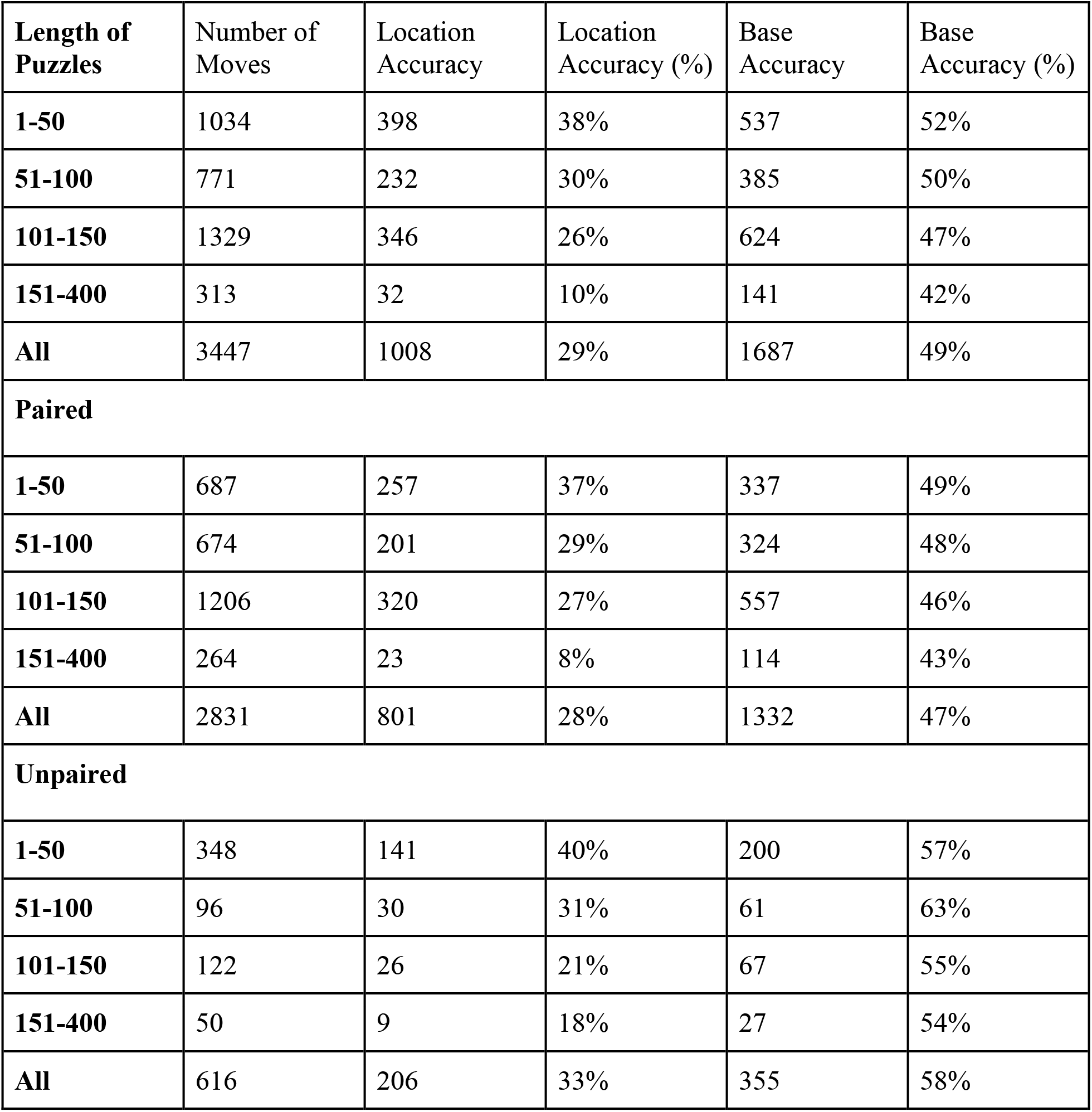
EternaBrain CNN accuracies on *eternamoves-select*, grouped by length of puzzle and paired/unpaired status of nucleotide at which move was applied.

The improved CNN accuracies using *eternamoves-select* compared to *eternamoves-large* suggested that moves from experienced players and on specially designed problems were sufficiently stereotyped so as to allow for CNN prediction. To further explore whether these improvements could be generalized across different puzzles or between different expert players, we split *eternamoves-select* into training and test sets in different ways (see Table 2). Rather than allowing moves in the training and test sets to be drawn from the same puzzles (as above), we trained our CNNs on 15,265 moves from 39 of the 78 puzzles in our training set. The predictive accuracies on the 15,182 moves from the remaining 39 puzzles were 0.26 and 0.023 (BP-CNN and LP-CNN, respectively). These numbers were lower than for our initial CNN tests on *eternamoves-select*, in which the training and test set drew moves from the same puzzles. Indeed, the BP-CNN accuracy dropped below the random baseline value of 0.33 when the puzzles in the training and test set were separated. These observations indicate that the CNN predictions depend on the overlap between puzzles seen in training and testing. We next split the *eternamoves-select* dataset so that the moves in the training set derived from half of the expert players. The test accuracies of our models applied to the dataset of moves from the remaining half of expert players were 0.38 and 0.11. These accuracies remained above random baselines (0.33 and 0.031, respectively) and suggested that, while expert players vary in their puzzle solving styles, there are some commonalities that can be learned by the CNNs. Last, we trained our models on just 587 moves of a single expert player and tested its performance on the remaining 29,877 moves in the *eternamoves-select* dataset. The resulting test accuracies were 0.27 (BP-CNN) and 0.011 (LP-CNN) - significantly worse than above. However, this poor result may be due to the dramatic decrease in training data for the CNN compared to prior comparisons. Moving forward, we decided to utilize the CNN trained on our original random split of *eternamoves-select* across all puzzles and expert players, but taking note that its move prediction accuracy would likely depend on similarity of any new challenge puzzles to the puzzles in the training set. We named this predictor the EternaBrain CNN.

### Playouts on the Eterna100 benchmark

After the cross-validation tests above, we set out to solve complete puzzles with the EternaBrain CNN model trained on *eternamoves-select* and the stochastic playout scheme described above. Encouragingly, the model was successfully able to solve several puzzles completely, including the *Simple Hairpin* puzzle, which the CNN trained on *eternamoves-large* failed to do, as described earlier. However, EternaBrain CNN failed to solve puzzles greater than 40 nucleotides in length. Visual inspection of the CNN’s series of moves revealed minor but obvious mistakes, such as mismatched nucleotides at pairs of positions that should have been Watson-Crick paired in the target structure. In order to prevent some of these mistakes, we followed the CNN with a second stage that we called single action playout (SAP; Figure 1E). SAP uses six canonical strategies that are standard among Eterna players and are taught to new players during the game’s main puzzle progression (Figure 2). SAP traversed the puzzle to find areas that were not folding correctly, implemented the relevant strategies, and accepted the sequences if they made these specific areas of the puzzle fold correctly; Figure 2 and Methods give more detailed descriptions of the SAP’s standard Watson-Crick pairing rule, the G-C end pair rule, the G-internal loop and hairpin-loop boosts, the U-G-U-G super boost, and the flipping pairs strategy.

**Figure 2.**
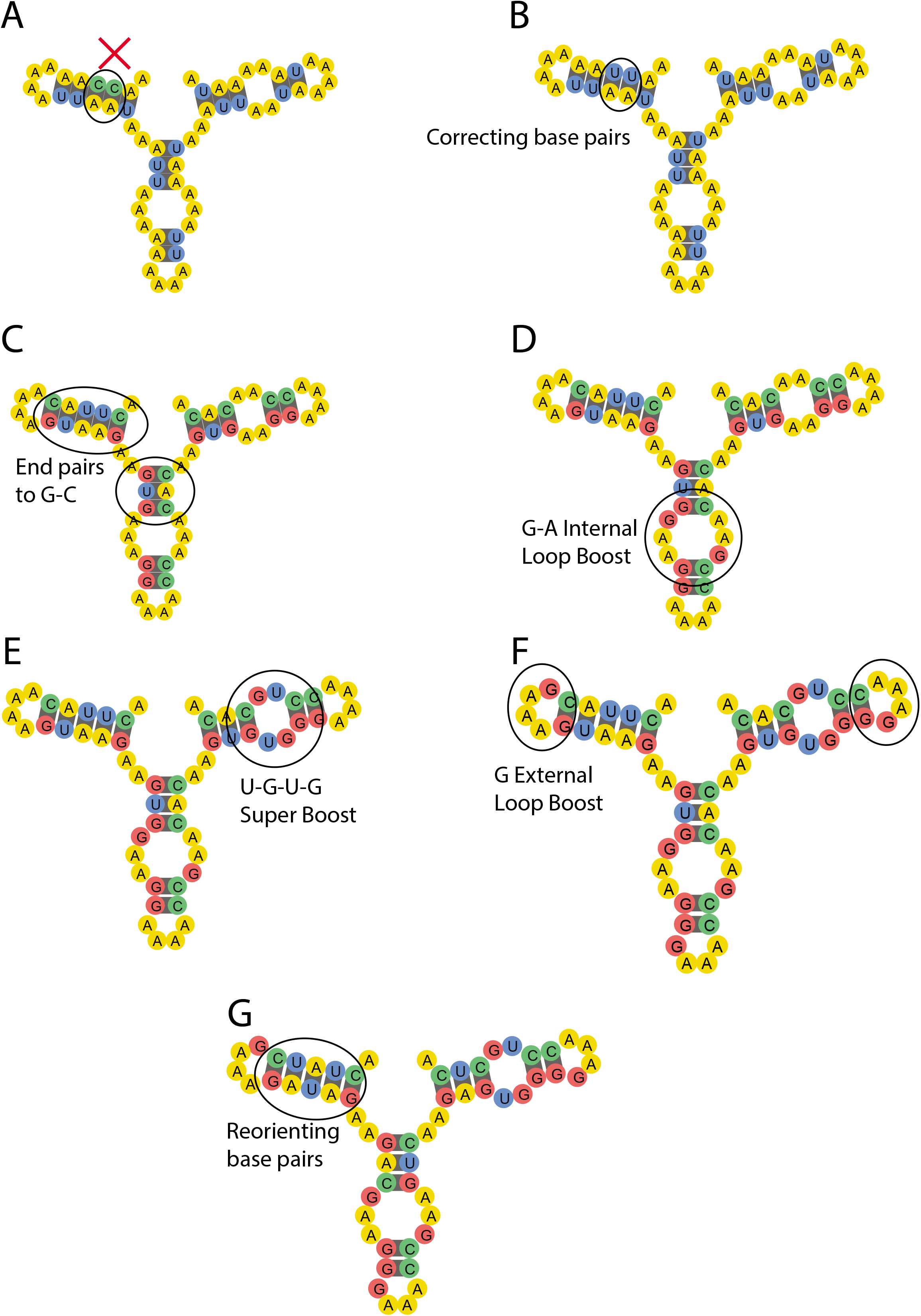
The 6 strategies included in the SAP. (A) The original state of the puzzle before SAP. This represents a puzzle initiated with an arbitrary sequence of nucleotides; panel displays the target structure, where mismatched nucleotides (C-A) are highlighted. (B) The first step of the SAP is to correct mismatched pairs. Here, the cytosine nucleotides are switched to uracil to pair with adenine. (C) Changing end pairs to G-C. Changing base pairs that are at the edges of stems and flank loops to G-C pairs lowers the free energy of the molecule. (D) G-internal loop boost. The first nucleotide in an internal loop on either side is switched to a guanine. (E) U-G-U-G super boost. In an internal loop with 2 unpaired bases on either side, the 2 bases are changed to uracil and guanine, in that order, on either side. (F) G-hairpin boost. The first nucleotide in each strand of a hairpin loop is changed to a guanine. (G) Reorienting base pairs. Target base pairs that are not predicted to be folded correctly are ‘flipped’ to lower the energy of the structure. Here, alternating the A-U pairs lowers the energy of the stack. The 5’ end of each puzzle is at the top left, with the puzzle drawn counter-clockwise from that point.

We called this enhanced pipeline the EternaBrain-SAP method and tested it against the Eterna100 benchmark. These 100 puzzles span a wide range of different structural motifs but are known to be solvable in the Turner1999 energy model, which is widely used in RNA structure prediction software packages including the original RNAinverse and NUPACK (6, 9). EternaBrain-SAP was able to solve 61 of the 100 puzzles, surpassing six of the algorithms previously benchmarked with the Eterna100 puzzles (Figure 3A). We note, however, that newer design methods using complementary algorithmic strategies provide similar or better performance compare to EternaBrain-SAP, and these will be discussed below.

**Figure 3.**
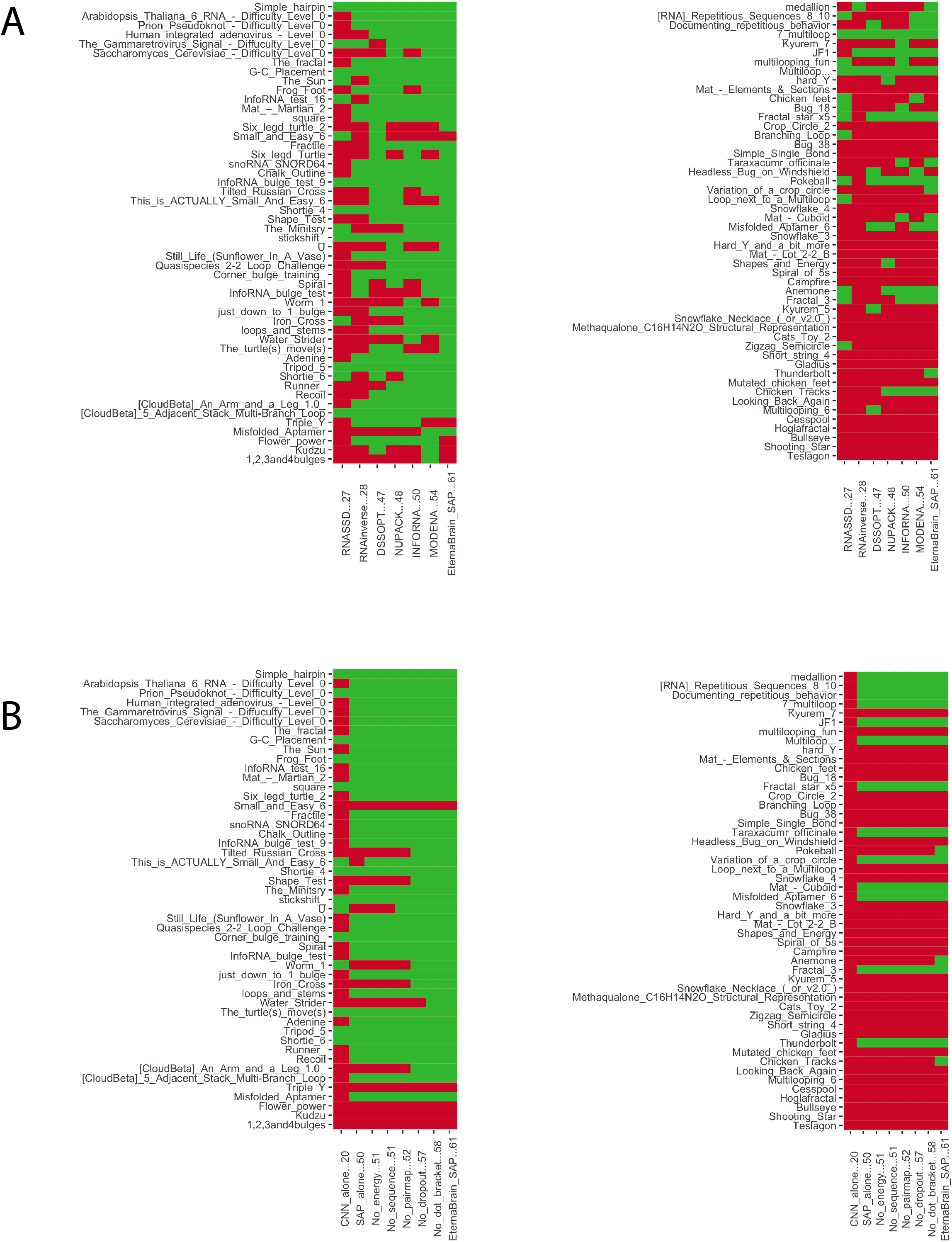
EternaBrain Performance. (A) Performance of EternaBrain and 6 previously published algorithms on Eterna100 benchmark. EternaBrain solves 61/100, followed by MODENA (54/100), INFO-RNA (50/100), NUPACK (48/100), DSS-Opt (47/100), RNAinverse (28/100), and RNA-SSD (27/100). (B) Performance of Alternative Model Constructions. The CNN alone could solve only 20/100, and the SAP alone could solve 50/100. Removing various input features passed into the CNN resulted in drops in performance, confirming the importance of these features.

Importantly, both the CNN and the SAP stages were necessary for achieving this performance. Using EternaBrain’s CNN-based moves alone solves only 20 puzzles on the Eterna100. Using the SAP alone (i.e., hard-coded canonical player strategies) solves 50 puzzles on the Eterna100, 11 short of the 61 puzzles solved by the CNN-SAP combination. However, it is important to note that the puzzles become significantly more difficult, as evidenced by a steep decrease in the number of players and prior algorithms who have solved them with increasing number (Figure 3A and (10)); and so this represents a significant improvement over SAP alone. We confirmed that several choices that we made for the CNN architecture and game state representations (Table 1) were important for the success of EternaBrain-SAP. Removing the sequence, the pairmap, or the dot-bracket representation as input features or not using dropout also decreased performance in Eterna100 playouts (from 61 to 51, 52, 56, and 57 puzzles solved, respectively). A puzzle-by-puzzle breakdown of the performance of the CNN alone, the SAP alone, and CNN-SAP with these reduced-input-feature training sets is given in Figure 3B. We also note that while EternaBrain’s CNN-based moves have a stochastic component (see Methods), its success on puzzles is consistent across runs using different random seeds (Table 4).

**Table 4.** EternaBrain-SAP performance on Eterna100 upon five additional playouts on the 61 puzzles it solved in its first run.

| **Puzzle** | **Number**<br>**of Times**<br>**Solved**<br>**out of 5** | **Puzzle** | **Number**<br>**of Times**<br>**Solved**<br>**out of 5** |
| --- | --- | --- | --- |
| Simple_hairpin | 5 | medallion | 4 |
| Arabidopsis_Thaliana_6_RNA_-<br>_Difficulty_Level_0 | 5 | [RNA]_Repetitious_Sequences_8<br>_10 | 5 |
| Prion_Pseudoknot_-<br>_Difficulty_Level_0 | 5 | Documenting_repetitious_behavi<br>or | 5 |
| Human_integrated_adenovirus_-<br>_Level_0 | 4 | 7_multiloop | 5 |
| The_Gammaretrovirus_Signal_-<br>_Diffuculty_Level_0 | 5 | Kyurem_7 | 0 |
| Saccharomyces_Cerevisiae_-<br>_Difficulty_Level_0 | 5 | JF1 | 5 |
| The_fractal | 5 | multilooping_fun | 0 |
| G-C_Placement | 5 | Multiloop... | 3 |
| The_Sun | 5 | hard_Y | 0 |
| Frog_Foot | 5 | Mat_-_Elements_&_Sections | 0 |
| InfoRNA_test_16 | 5 | Chicken_feet | 0 |
| Mat_-_Martian_2 | 5 | Bug_18 | 0 |
| square | 5 | Fractal_star_x5 | 5 |
| Six_legd_turtle_2 | 5 | Crop_Circle_2 | 0 |
| Small_and_Easy_6 | 0 | Branching_Loop | 0 |
| Fractile | 5 | Bug_38 | 0 |
| Six_legd_turtle_2 | 5 | Simple_Single_Bond | 0 |
| snoRNA_SNORD64 | 5 | Taraxacumr_officinale | 5 |
| Chalk_Outline | 4 | Headless_Bug_on_Windshield | 0 |
| InfoRNA_bulge_test_9 | 4 | Pokeball | 1 |
| Tilted_Russian_Cross | 5 | Variation_of_a_crop_circle | 3 |
| This_is_ACTUALLY_Small_And_Easy_6 | 5 | Loop_next_to_a_Multiloop | 0 |
| Shortie_4 | 5 | Snowflake_4 | 0 |
| Shape_Test | 3 | Mat_-_Cuboid | 3 |
| The_Ministry | 5 | Misfolded_Aptamer_6 | 2 |
| stickshift_ | 5 | Snowflake_3 | 0 |
| U | 4 | Hard_Y_and_a_bit_more | 0 |
| Still_Life_(Sunflower_In_A_Vase) | 5 | Mat_-_Lot_2-2_B | 0 |
| Quasispecies_2-2_Loop_Challenge | 4 | Shapes_and_Energy | 0 |
| Corner_bulge_training_ | 5 | Spiral_of_5s | 0 |
| Spiral | 5 | Campfire | 0 |
| InfoRNA_bulge_test | 5 | Anemone | 3 |
| Worm_1 | 4 | Fractal_3 | 5 |
| just_down_to_1_bulge | 3 | Kyurem_5 | 0 |
| Iron_Cross | 5 | Snowflake_Necklace_(or_v2.0_) | 0 |
| loops_and_stems | 5 | Methaqualone_C16H14N2O_Structural_Representation | 0 |
| Water_Strider | 5 | Cats_Toy_2 | 0 |
| The_turtle(s)_move(s) | 5 | Zigzag_Semicircle | 0 |
| Adenine | 5 | Short_string_4 | 0 |
| Tripod_5 | 4 | Gladius | 0 |
| Shortie_6 | 5 | Thunderbolt | 4 |
| Runner_ | 5 | Mutated_chicken_feet | 0 |
| Recoil | 5 | Chicken_Tracks | 2 |
| [CloudBeta]_An_Arm_and_a_Leg_1.0_ | 5 | Looking_Back_Again | 0 |
| [CloudBeta]_5_Adjacent_Stack_Multi-Branch_Loop | 5 | Multilooping_6 | 0 |
| Triple_Y | 0 | Cesspool | 0 |
| Misfolded_Aptamer | 4 | Hoglafractal | 0 |
| Flower_power | 0 | Bullseye | 0 |
| Kudzu | 0 | Shooting_Star | 0 |
| 1,2,3and4bulges | 0 | Teslagon | 0 |

### Performance on specific features of difficult puzzles

The Eterna100 benchmark contains puzzles of increasing difficulty that showcase structural features that are difficult for RNA inverse folding algorithms to design. Comparison of EternaBrain-SAP’s performance across these different puzzle types clarifies its current abilities and limitations.

#### Simple Motifs - Stacks, Loops, Hairpins

The EternaBrain-SAP algorithm was particularly successful at solving puzzles that contained several stacks (i.e., RNA stems), loops (i.e., internal loops or two-way junctions), and hairpins (i.e., external loops). One example of a difficult puzzle that EternaBrain’s CNN was able to solve is *U*. This puzzle contains several short stacks and short loops abutted next to each other (Figure 4A), and was not solvable by the SAP alone or by five of the six prior design algorithms (Figure 3A). This example suggests that EternaBrain had successfully learned from its training data how to strengthen stacks and to stabilize loops. Specifically, EternaBrain’s CNN was able to learn strategies like the G-C end pair strategy (which strengthens a base pair stack) and the G-hairpin boost (which stabilizes an internal loop); see Figures 2C and 2F. While these strategies are well known and, indeed, are encoded in the SAP (Figure 2), the EternaBrain CNN learned further patterns that were not encoded in the SAP, as is demonstrated by the inability of SAP alone to solve the puzzle *U* (Figure 3B).

**Figure 4.**
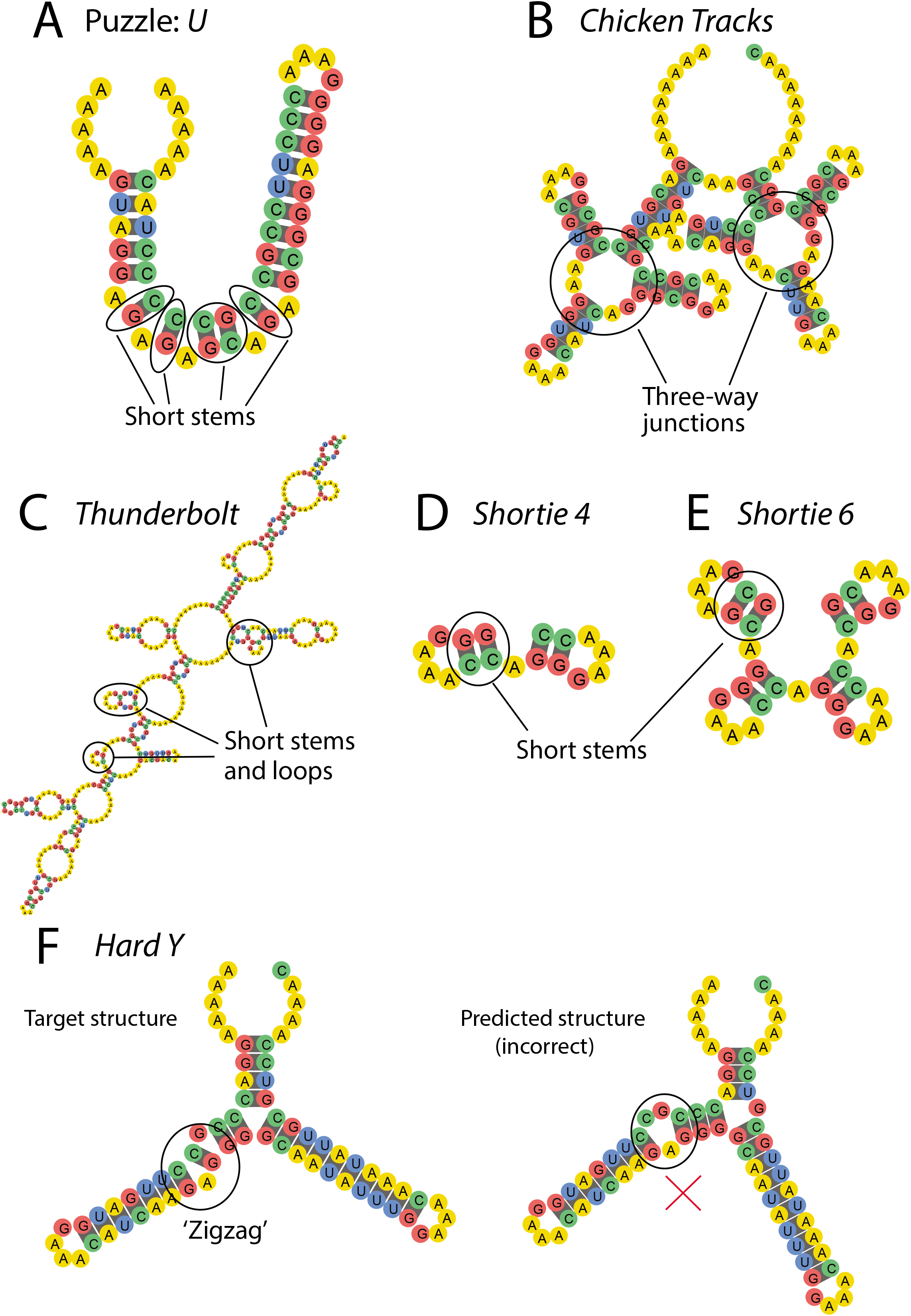
Example EternaBrain-SAP solutions to Eterna100 puzzles. (A) *U* solution highlights the fact that the EternaBrain CNN alone can solve puzzles with short stems. (B) *Chicken Tracks* solution: EternaBrain-SAP can solve puzzles with three stems intersecting in one internal loop. (C) *Thunderbolt* solution demonstrates that EternaBrain-SAP can solve large puzzles (400 nucleotides long) and solve loops and stems in combination. (D) *Shortie 4* solution shows EternaBrain-SAP can solve puzzles with multiple short stems (2 nucleotides long). (E) *Shortie 6* is quite similar to *Shortie 4*, but with the same motif (short stems) repeated. The other algorithms mentioned could not solve *Shortie 6* because of the repeated motifs. (F) *Hard Y* - target structure (left) vs nature-mode (right) structure. EternaBrain-SAP could not solve *Hard Y* because it required use of a little-used strategy to solve a motif called a zigzag. Since the strategy is not often used by players, the EternaBrain CNN did not learn the strategy and the strategy was not included in the SAP. In each panel, the 5’ end of each puzzle is at the top left, with the puzzle drawn counter-clockwise from that point.

#### Specific Orientation of Base Pairs

If the puzzle contained motifs whose solutions critically depended on unique sequences, the CNN was inadequate and SAP was important for EternaBrain-SAP’s success. An example of such a puzzle was *Chicken Tracks* which required a very specific orientation of base pairs (Figure 4B). As a result, the SAP was used heavily in solving *Chicken Tracks*. By locating the areas that were not folding correctly and then using the canonical player strategy of reorienting base pairs, the SAP was able to find the optimal orientation of the base pairs to correctly solve *Chicken Tracks*.

#### Repetitive Structures

Some of EternaBrain-SAP’s successes can be attributed to its ability to solve repetitive structures better than previous automated RNA design algorithms. Example puzzles that include many repetitive structures include *Thunderbolt, Shortie 4*, and *Shortie 6* (Figures 4C-E). Previous algorithms were able to solve puzzles with several stacks, such as *Shortie 4* (Figure 4D). However, when the number of stacks increased, e.g., in *Shortie 6* (Figure 4E), other algorithms struggled. EternaBrain-SAP, however, could solve both *Shortie 4* and *Shortie 6. Thunderbolt* involves repeating elements within a large structure (Figure 4C) and also was not solvable by prior algorithms (Figure 3A). Given that SAP was able to solve each of these puzzles alone (Figure 3B), the moveset-trained CNN of EternaBrain appears to offer little benefit over SAP alone in stabilizing repetitive structures.

EternaBrain-SAP’s failure to solve other puzzles in the benchmark seems, in part, to be a result of the incompleteness of the training data for the EternaBrain CNN and of the player strategies in the SAP algorithm. For example, *Hard Y* (Figure 4F), requires uncommon strategies, including a different type of boost (a stabilizer mutation at the beginning of a loop) to stabilize a special ‘zigzag’ structural motif which did not appear in the training set. The SAP was unable to solve this puzzle since reorienting bases without the required boost did not stabilize this zigzag. Training the CNN on larger movesets and incorporating more sophisticated player strategies in SAP might resolve these issues and help EternaBrain-SAP complete more complex puzzles.

## Discussion

EternaBrain-SAP attempts to solve the RNA secondary structure design problem by learning from a large compilation of human player moves and incorporating a special set of canonical player strategies. We decided to use a convolutional neural network (CNN) since it can be trained to extract information from the nearest neighbors of elements in the feature space, mimicking how Eterna players look at the local neighborhoods of structures and nucleotides to decide their next move. To reduce variance in the training set, we found it important to use movesets only from expert players. By reducing the full *eternamoves-large* moveset (1.8 million moves) to the *eternamoves-select* move set (~30,000 moves), we were able to achieve test accuracies after CNN training that were better than random guesses. Notably, for the location predictor (LP-CNN), the test accuracy of 0.34 substantially exceeded the baseline prediction accuracy for random guessing (0.019).

Despite decreased variance from training on the *eterna-select* moveset, we found that the resulting EternaBrain CNN had difficulty with solving longer puzzles. Indeed, alternative splitting of test and training sets suggested that the CNNs’ prediction accuracy largely depends on similarity of puzzles in the training set with the puzzles of any new challenges. The inability to extrapolate “out of sample” has been noted to be a limitation of artificial neural networks (27). To overcome this limitation, we added a single-action playout (SAP) algorithm based on compiled Eterna player strategies to aid the CNN model. We tested this hybrid EternaBrain-SAP algorithm on the Eterna100, and found that it could solve 61 of 100 puzzles. On one hand, this performance is better than other algorithms that had been tested on the Eterna100 at the time of development (54 of 100 was the previous maximum). On the other hand, the EternaBrain-SAP performance is similar to or worse than newer algorithms (SIMARD, sentRNA, the reinforcement learning algorithm of Eastman et al., NEMO) that have been developed concomitantly or after EternaBrain and whose performance on the Eterna100 has been reported in newer papers or preprints (15–19). Furthermore, none of these methods match the level of the top ten Eterna human players, who can solve all 100 puzzles of the Eterna100 benchmark.

It is instructive to compare EternaBrain-SAP to the newer generation of RNA design algorithms. Like EternaBrain-SAP, the new methods SentRNA, the Eastman et al. method, and LEARNA use artificial neural networks to distill potentially useful information from gameplay and solve 80, 60, and 65 out of 100 Eterna100 puzzles, respectively (15, 16, 18). SentRNA seeks to find solutions to RNA secondary design problems in ‘one shot’ rather than through EternaBrain’s iterative moves (18). Furthermore, SentRNA differs from EternaBrain as it is trained on Eterna player’s solutions (<u>the *eternasolves* data set</u>) rather than the individual moves that lead to solutions, and it makes use of a three-layer fully-connected neural network rather than EternaBrain’s deep convolutional neural network. Despite these differences, both the SentRNA and our EternaBrain-SAP study find that neural network approaches alone give poor performance in test puzzles (in both cases solving fewer than half of the Eterna100 puzzles). The success of both studies required pipelining starting solutions from neural network approaches with hand-coded strategies that Eterna players collectively learned and ‘canonized’ in tutorial puzzles for new players. The Eastman et al. and LEARNA algorithms (15)(16) are reinforcement learning methods which have not leveraged prior human solutions or strategies at all; nevertheless, both use additional hand-coded rules to enforce Watson-Crick pairing of nucleotides during design and, in the case of LEARNA, an additional refinement step. Furthermore, at least the Eastman et al. method does not learn ‘standard’ human strategies like the G-boosts (Figure 2), which might explain weaker performance compared to EternaBrain-SAP and SentRNA.

Given these initial results from neural network methods, we propose future updates that may allow automated design to reach the level of expert players and to allow for a general strategy that will work on more complex RNA design problems including multi-state switch design and 3D structure design. First, EternaBrain-SAP and SentRNA achieve success on different problems in the Eterna100. This observation suggests that the two methods could be integrated, with each one providing starter solutions for the other, or the convolutional neural network architecture from EternaBrain used as an alternative neural network for SentRNA. Second, the hand-coded player strategies used in both EternaBrain and SentRNA often involve multiple moves. These strategies could possibly be captured by the neural networks underlying all four available deep learning approaches if they are trained to make moves based not just on its current game state but also including immediately previous moves as input. For example, such training would allow EternaBrain-SAP to ensure Watson-Crick compatibility across nucleotides that are paired in the target structure, an ‘obvious’ feature that the CNN is not always recognizing and that had to be hard-coded into the Eastman et al. method and LEARNA. Finally, it is important to point out that more advanced deep learning architectures may outperform these approaches and achieve human-level performance. Recent results in protein structure prediction (13th Community Wide Experiment on the Critical Assessment of Techniques for Protein Structure Prediction, http://predictioncenter.org/casp13/index.cgi) demonstrate how rapidly such gains are happening. These methods have benefitted strongly from large data sets drawn from the large public archives of protein sequence and structural data; for the RNA design problem, the *eternamoves* and *eternasolves* data sets provide potentially analogous data.

Complementary gains in automated RNA design methods may come not from fine-tuning the neural network architectures or training sets, but through adoption or integration of more completely distinct strategies. Methods like SIMARD (17, 28), antaRNA (20), and MCTS-RNA (21) achieve performances similar to or slightly better than EternaBrain (54 to 67 out of 100) while using quite different but complementary simulated annealing, ant colony optimization, and Monte Carlo tree search strategies, respectively. Perhaps most strikingly, one of us (FP) has recently reported that a nested Monte Carlo (NEMO) method can solve 98/100 puzzles in the Eterna100 benchmark, only missing the last two problems, called *Shooting Star* and *Teslagon* (19). While NEMO fills in candidate solutions from 5’ to 3’, like SentRNA, other aspects of the method are completely distinct, and NEMO does not use a neural network approach. Second, we note that human Eterna players often solve complex RNA design puzzles by manually preparing sub-puzzles and using detailed mathematical reasoning to infer solutions. Neither of these steps is recorded in the Eterna move sets or has, to our knowledge, been captured in design algorithms to date. Finally, other methods like RNAifold (4) and the recently presented RNAstructure *Design_preselected* routine (22) have yet to be tested on the Eterna100 benchmark; they may show promise on puzzles that are not solvable by other methods.

In addition to their prospects for achieving human-competitive performance on *in silico* single structure design, we speculate that CNN-based move prediction and deep learning frameworks will be useful for design of functional RNAs that actually work *in vitro* or *in vivo*. The Eterna project is currently soliciting designs for ligand-responsive multi-state riboswitches and for redesigning large biological RNAs like the ribosome for *in vitro* tests. Current computational design methods are not able to automatically provide solutions to these problems. When these computational methods are ready, datasets involving hundreds of thousands of RNA molecules are accumulating in the Eterna project (27) and should provide rich resources for training and prospective tests.

## Acknowledgements

We thank J. Nicol for expert technical assistance; J. Shi, M. Wu, P. Eastman, and B. Ramsundar for scientific discussions; and M. Gotrik for comments on the manuscript. We acknowledge funding from a Stanford Graduate Fellowship (to B.K.), the U.S. National Institutes of Health (R01 GM100953 and R35 GM122579 to R.D.), and a Discovery Innovation Award (Stanford University School of Medicine to R.D.). We acknowledge the following Eterna participants for designing the puzzles used to develop EternaBrain: Kieros, jandersonlee, drake178, Brourd, hoglahoo, Janelle, eternacloud, Hyphema, RedSpah, nihilnove, steven123505, player4596, portalbob340, mat747, Jieux, redsoxwy, Eli Fisker, RedSimple, Malcolm, firedrake969, pdub93, ElNando888, Dennis9600, Nidoking, cake. We further acknowledge the following Eterna participants for taking part in solving the puzzles used to develop EternaBrain, with contributions ordered from most moves to fewest: Eli Fisker, dl2007, Pi, DeNa, mat747, HarryS, Malcolm, Jieux, eternacac, JR, benrh, cynwulf28, Omei, Poll na gColm, GFRANK2, JSci, Brourd, spvincent, whbob, tommyd, skyblue, dizzywings, lroppy, AndrewKae, atanas.atanasov, rxmullin, worseize, ElNando888, hoglahoo, holoaaron, 137.036, jandersonlee, Astromon, Brevitz, bjorns, novice, BHunter, chloep17, Tesla’sDisciple, kriss888, Sgtbird08, lcurtisadams, carmenmarbella, pmlkjn, a1of2twin2016, quantropy, JONATHANROHR, RasmusVH, RAH1, Fusion7, Warren09, Gres, ulfang, joy45, Zanna, D’Wydd, garrin, yceev, Arnthi_470, astralcrescent, erwinmulders, MacSousa, garydfisher, AndrewReedy13, sn69, Starna, Gracier, wawan151, ZOONIVERSE12, wateronthemoon, Radioactive7, IceBolt, ererexiue, LarryB, camdenmurray, manzet, armin, ppgbubbles, FurElise, biggestlegoheroicafanever, AnnYang, Snowpup, chase2003, eagercheesecake, Sven4, heynorma, DaIto, jsydrah, dudbomber, incal11, kwharton, fallenwarior, Citizen Science Guy, Sargon, c-quence, LiquidOvar, dblueskye, ZachFR, Shayera, mastorasa, Grelko, scientificdash, Marculius, SockTaters, elasmo, TheDomBom13, jIyHHbIu, davidwhittaker, ecedenia, Ethaniel, kkdixon, ciprian, JoshS5, doofenshmirtz, justintwayland, jbrandim, MeowNow360, delcastle, Pytho, AustonTheriault, Eized, BirchSci25, cjaltrichter, Nirrame, VeraMarsova, CarlosFaustino, billygoat, everyday847, Matias Nicolas, wookieetank, Citizen Bane, ShelleyVAdams, gian666, skamil1204, PaulaSphere, SourCreamKing, ChrisRWitt, tone, annamary488, ljchapa, king1000, Pilot69, sellena, Loreleykaa, tamaki, tmjlowe, danielmcmurray, DarkAndroid, bbueno5000, SalarianScientist, Wulveniel, rchrdjhnsn, Overide, mtor1, henege, Masterbajurf, Leepretorius, janetmason, Tati69, alebec, lindelenilda, dillon101001, kkowales1, Mongoose, rocketdog42, apeter14, Nova631693, Ekymose, chanceyko, RunDMC, Armordines, pamisza, a1puterboy, mzeinstra, dragoncendre, Radek24, Psykovsky, MageOfTime, kiernan2, marcuac, jessesmall, jonah.pandey, booti386, LAdarkTrooper, qq47, D1l3mm4, science123, Zugai, Felixpi, Aerendil, PereKastor, 200611736, mjmtl, akhyatt, mircea, Suboptimalautomation, Menosfree, smwalsh1228, StormTroop, Sylva, trashy, Jeshyr, SolubleCarpet, EcceruElme, RNA123, tklight13, Darkasleif, GermanP, Lewis656, nico1630, bjackblack, TheStaker07, kylelyk, Sean O’Neil, jnicol, Gigante, mariapaglinawan, bryanb2, YenRaven, TAG, chalouky, onilink_, Tessaract2, MasterStormer, deltaq84, Suprem, Renthel, awesomehuman, cosmia, cmmtchll, Xnessax, angusd, Ambiseus, nuqotw, Manabender, oli.veri, Lyssaodr, alacarus, chop&toss, XFYLESGIRL20, Trentis1, manthonyd1618, wkdus108, Taflin, Mayanne, lastresortist, xotame, Hyphema, MThrasher0, Baron, Moir, dbm, JulesLeNomad, Soken, Meechl, norikusan, AxelHawke, Jimedney, redrocz, HawkEyesMiHawk, Marzena11, emjay, zsherman, sing146, acebassmaster, Njdilaan, hai-meh, Calanor, kersevich, AshlyW42, Aleks59, freznow, anik, all, taslack, Ipiu, SkyDancer, alecpikachu, SrdjanM, rlhruza, saili6, potato43, pabloguillen, Nsyp, 2016_talias, RayA, Egin, AmyW, sneaky90210, Monsoon, EPogue, Antireta, jahfe, whispywhip, rubixcube517, pyrazole, trili, Willcalliw, sebC, akio123, grass, Cawa, RechleckiJ, aceiestatheist, randomfull9, paÁoqueiro, madman200, PFKThorin, HARDWAREGUY, Ibanezmtl, chuckcoleman, harpokrates, Vincorporated, AHamel, Eceri, KT006, GColdwell, adamsunny, Niemand, marisa31, mrbell, TedStudley, jyoshimi, Approved, IchigoNandato, PenguinLea, stevetclark, bbk123, sharmoni, melliniumfalcon1029, JennTM, Natura, bookbird, Duha Eldow, Teutonius, mdurbin, kartonrealistah, rafa_prog, 12walch, bkopel, TakumaInoue, caro9923, Theos, LordHaralVII, legotourist, ThatYouGoOn, 1900029, theta123, raek57, MysteryBacon, Dragonbard, redkatie, Purple wizard, zombiekill6r, Rouge Lead, 577661, wnxkitchenaid, cutkzig, Sporeo, System0X, sekeerne, Skye, LCL46fgh7, lferris1, Jt82497, scooper1, Happymuffen, Aleksandr, maschiltz, PeakeHaus, Doctor_Maximus, coolbeans55555, dmitrybaianov, Kiddodog, tdmcmillen, LacXav, StuartC, DGFigueiroa, EmeraldDragon, kmille, Paulo Roque, rdfgiraldez, Stephanie1978a, turing999, balsamina, rdaleg3, vdvd138, darkseariver, PIRSQUARED, Marquis Tinkles, stormyx13, rhys, fareedanyc, wallr1, RNADesigner, zacthemac753, kang36, cptguru, Schwarzee, stephancannon, leanicholewhite, Maya are you here, seriousgedas, Abilatanwen, rjonzales, vjayant, adibou_super_saiyan, silent dicer, sheyla, starreus, choo, aclockworkkc, pksvseng, lizdeath, Vorushea, CaptainColon, Billy Reuben, LaMariposa, Bekreth, captainpsychology1, iojp, PFChopZ, jackieofall, Anorectic99, firefliesjr, Professor Lamp, Mathiszenoob, Alec_7, FelisAnxietus, Gwindorzp, microbiogenius, Jack123X, lqtza, tc5801, jgabbott, lilyflan12, hi25114, Abdullah, simonk, funglish, joe911, Ozzy1313, TheDoctorInTheHouse, jcastello, ExternalCPU, zeroxoxo, zakblak, couchkitty, hizowie, Orkhan, arueda, randomone66, Eliisme, player613, nijaho, kyger1, mh.ku, Bpala, dmeyer, dentongal, Andoryuuta, rguioguio, osomblossom, MauroBedoya, Kienan2, Hayfieldlee, hibts, kijizz, lkjhnsn7, bholcombe1, RayNA, serinorah, smuthuganesh, Science Ferret, danryk, cwillms, BletchleyPark, roughb06, Jesse2016, Tigerlrg245, Magmatic, trapman, sergio_ruelas1, caroduf82, blueskycrf, Some Guy, gunjan.bhattarai, scharpentier, scash, cathaybrent, wayyne, brendanrmills, fangu42, wiillem, cheeka, gugish111, namchokdef, muenzer, Reklaw1973, aurora999, Monarch, hcvalentio, yamanq, sorgd, varjo1, StevenKell, emilat, JamesB, LordKunrath, sahndie, dein_kommandant, iamchriskelley, epicfalcon15, Kafi, SherDG, Klondike, radigan15, whatnot, BrosefStalin, anto9uspius, GMark, Alipony, citizenkane883, Corozon, Judric, mrfinntastic, Dalcus, georgethebeare, BlueBotanist, aviator56638, antimatter, umbrellaPrince, Benjamin_Li, deadifieded, Computerion, -alina-, sealyt, Ex, Kelyank, domdomdeo, Zboubine, vinniec7897, delaneyyyw, Histerion, viperious, eec221b, ggjeaneude, matthewtyler1, rna_enthusiast, Rivalium, jellska, jagodzinskaj, TB killer, B-Rabbit, potatoman12345, BichirM, UnkDevE, hwilkins, doodeoo, BKC, felix1998, antonchristoff, 18hhochstedler, pszym3, saidai-no, pansap99, Acuzik, kelvin.yu4, Crolius, Theknight7, RhysMac, Nightwarex, Morula, dominatingX, Vick, PVequalsNRT, televisaos, simonvozar, phantomrequiem, mrcookies, moleculadesigner, denisb, athakore, Viktor.mg, gpc5000, Go Lem, boomdude63, stephenb4, blaverentz, Raju, zhenning, haupt, StarScience, andrewcailliet, hualiama, scobyke, Ruda999, Z-Man, Unkn0wn376, kingboo, jedimasterhanuman, atomoton, hamtruon, Kremmen, Spooneres, Hypernova, DarrenLott, bolgir, tdarron, polly66017, Lalourche, yhomas, tjxp, ajvenaje_md, Kamalini456, dennisprokofiev, pierceh2, JiggidySwiggidyGiggidyRiggidyJess, Macjack, blakeballinger, fra888, Talmage, drake178, devilish, mjythgr, JohnCena, rye.coombs, AdaptiveCoffee, greendude, Pirus, jmayberry, DnL, Towel42, pinecat, zacharykent, timotk12, CAAndrew, Inosen, Fawn, pensive_sloth, ezbot, JohnDangle, artarius1, solar_traveller, dkbarnes2, emerilin, james.mackay, ethan_13, diamondscar, drakereborn, Handerson97, Ribonucleic Acid Junkie, MegaManSam, slymmer, xxxanderrr, kchambliss, Zainab Badejo, fafnir, gzhu22, kvalder1, soupnix, animal13, Dexar, aet36, jaycebradley, salmykanit, Delta168, jzaki1, hell2950, inchoatewaffle, Hank McCoy, EdK, Ayedyn, MCuis, NickyNewark, Strikefusion, natalieweissman, Cactus Jack, saxyness, Wcarta, Garod, LeCHIrurg, DeepRiver, xadad, Source de Lune, StajaStorm, durka1000, Moon Man, Kemono, argo007, jAEROd, markshvili, electro2678, JMZ, LordSoulis, tartule, Mr. Solo, shridhar, reginn, Nilbonur, mynameisbob, vogtmich, MrMean706, hognmeister, hkhan249, SumanV, pebbles, psifio87, tajclan, eiti13, doomie, yayforfood, LaraGazetta, Matt Indykiewicz, Broomster, TrulyAcerbic, komjum6, BurtonReed, a user of this program, propheteer, goerch, spiffykavu, Dywyn, DanielMaxwell, RaeVigil, Rzv314, yong syuan, joey70013, Wolf_01, Mikaeru, weaverl3, Moon, laurentbaba, Leila204, broofboob, Crat3s, zandramssystem, mtor2, babyFolder, migs, Duban, sasevy, tate, Jumanji, Zakki, EdouardBonnet, AaT, migulrocks, cryptc, fcell, clementin, syrthael, JMStiffler, alhariri, starkwinter, Loneliest, veer, tohtorisyker^, MatthewDrucker, rayhead, starski1490, Anteprefix, Nazamroth, sc, kevinyhong, anthonyperal36, jarrux, kolaps4, xb, ZeroKelvins, tardigradedg, Sparrow3183, curiositycreature, M-Stak, Safler, ottapav, ryisnelly, thanhngathinguyen, dheitmann, Morcha, Timm6539, andreyz4k, cave felem, SHv2, pez_eterna, MicroW, toneranger, wolfeac, azsr, chw68, thatguywithhippyhair, Salmj, nate300and4, wyom, telysea, johnsonalyssa325, AbyssalCeph, a52, Bio4Bio, LadyEnoati, Jo,,o Bernardo, schmuckerlucky5, b_ccruz34, RisingNucleotide, thserp, cd73, sixnorth, SirriuS, awangii, NikkiNeko2012, McHunsdonstein, slaterj2, artarius, mooper123, 02100, MartianTGI, brokenbadguy, jamjar, EwaD, alchen, upquark1, Josh198t4, comorbid19, milnus, Botvid123321, VoraxRaptor, ahearnc1, ahsan8244, Entropicana, Quantum, Aleboo1, miii_kiii, Spenni, IgnatzKackebart, SLSkiwi, flotterHecht, fayddle, Gage Boyd, johtso, Saitama, AlexMercer, Spartan0534, drewman, Doctara, cherry39, wingfly_1234, LFP6, CheatTheReaper, berserks, cavejohnson44, Bill Nye the Science Guy, koprocesor, Deafeater_PRO, Trip, hpets, 4plj4k, Gotoza, VirgieP3, user256, Gennady Stolyarov II, quest, kappa123, evanders, pearlofdarkness, ECAP, iwayzen, Infinity40, 0ah064d, Tauchsieder, Deadport, filipT22, alrianne, stelios3, AlxrrY, mdwoman, IVIadScientist, Frid.be, alynora, btbam01, greatdreamer, Zezel63, kartonrealista, luisraposo, spiritofbodiejulian, Ishimaru, ilovepi, kaNino, Janimbus, arma95, liligirl1234567890, sarkonic, BrittGay3, gross23, wolfhorse13, col_kurz, amaniduncker, fifoumax, abbifede89, bugspc, TylerMilam33, lamartinium, rsardone, Ulithariad, apocolypse45, ColeV, gigamech, SilentD1, abuyagop, dusnikj, bicykiel, caillou, Mason2222, cacamach, Thea, mscp, JulianZhu, jy3, MitchSqwuu, Tellis, thezymurgist, Elposze, emeraldflm, veerkhanna, Consilience, ViolettRein, thebigdoodoo, yacopoc, nkarray, Alejandra91, RosalineM, wolfxana, tamc, vatokykaika, nomad72, Bananadine, infeRNAl, enzo732, chuckleNchu, Madame GrËs, lorolo, Chanish, pagejade, Catcow, dragonfire1, Floris.s, noahsterdomas, bluenightcry, uhrag, metcys, nyegy, NVxWILDCATx12, ethong, frolickingspud, collegebookworm, ooloughlin3, catholicon, ascience4, Onymbit, BradenT77, annaofwonderland, KIM ji yeon, Richousrick, StuckinmyHead, alwin, mrjes, branzau, slimjimmy24, IZmanetje, IlPero, DylanH6, k14, Chloe_Kb, dronkihot, baileybalouch, Beauregard, Scientist formerly known as Rick, Anteers, starwars11, aakspuicomterixx, thedoctorgrace, Kesandro, Dikiyoba, trebor8201, it1023, codygeary, StAr, Bapt334, afoley947, Emeraldguy, Catch-22, Cdixson, kycklingar, gurumi, Elegans, Palatura, darwinsdog, jonnyflash, Firemyst, Daddy, Muzzhum, xSkyHigh12, SirFredrick231, Natabu, s202805, murphyhc, drewbert, Martian720, antsmarching, biogrl, White, GamerOnov, m4812, ammcelro, antreal, benacte, PJ, JanTheConqueror, NoDakTime, graph, kinsir78, llamadog007, nathanielbuck, Jpatino, kmccormick21, JEPR83, unlocked, deathbyfunz, jdude104, Prothon, rglowrey, i_am_syd_barrett, Seanser, BioChem1996, SeanMurphy, talwara, rtdub, blikdak, clayjmck, kovo_05, danlan, doradears, rsingiser, L.Klein, Aubade00, acipi9, hobbitluck, unusualbob, Nikenik, mohammadsdtmnd, mreyes, FlagFlayer, aubreyy96, rlharrison, Theocrates, lucaswinningham, salamalekos, sggursky, Freddog, gdnskye, eshun, Daderic, zoonut, EternaRNA2016612, ClemDavies, Artdog2005, Benzyl, karanlyons, maraam, heard-snow.m, Craahcourser, Leontis, Gummiworm, tgusta, valeriamin, ghosth24, WelpeAuto, epicFaces1130, vfrankenstein, matthias1373, dzsainz, voonkit, mitchowski, MistMask, mws, SebastianK, gwd1093, batochaussette314, xenon54xenon54, TheFoldinIowan, dwie, brapiro, mboozer, Sunder, jackmyth, dStreamline, mmariset, Spaceninja, CJH4, amaragold, Seikre, BrendyBoi, mevenvirgil, LonleyGhost, Sova001, cyanhivgp120, fildm, noahwhiteman24, enhandy, BLT, Ekiastheboss, trio, gem, amenephus, Gnida, kareybh, Anon1001, stevenglasford, ZenNOA, unseenantidote, 4terie, santiagoneym, Knight2050, wietze, Hydrvs, Tiro, a1b2, avery3r, Dischord, tuete444, mariamnd, ggrafham, marlasinger1470, Rheannon, speckle, titonus, DrCooper, nan, BroaderKey, ahocevar16, Niski, rrorapaugh, XXVI, starmile, 2thug, solrac, pcbPAK, Zavarin, Sirocco, emiliomendezpaz, 19772412, EHzg_Johann, golddragon21, Thep, Vergalon, eszipf, bekilewis, kworzy, norkie, siroderab, BetterCallSol, Zertu, TheFableFang, applesplice, DoppelBia, Saimon3, Cake or Death, mariska565, Lysergide, ghita, GamerGaming, Valdasgugu, Chadwig, steel112, Faithless, GeoPointer, poorigami, HarrisRG, MadRabbit, sherwoodtristyn, yarinetrobles, 2MG09, Kha’ri, Maristella, lapochita, medisad, briantingley, TerraForge, wadmanc, estebantucci, MCL, bialy3, Michaelaj1, kjtw64, tjscammell1, Poovent, Cpt_BA, nicholas938, Beligol, Noxbis, xxsodapopxx5, trystero, Thomas_T8, KCong13, Joshuahendrickson, xenorolla, ravenpuk, Tubbi, Evoluxman, Liber, Malafunkshun, Adorabel, aschachter, Bsmb1, JulianDelphinki, AlexSlimy, frostbite896, Abhinav Pujar, NicolasB_FR85, Z., dragonswordt, razorgrass, clementr, etIIenne, Huffsa, erenken20, Bjornfellhanded, portalbob340, coldwater, BigStinkyMoose, armerand, JustJoey, patbowne, mmmtstar, shaescarr, Buckeltulpe, fafv, bobthellama, tsaini, patandwerner, Madison P., mollycmcdonald, michaelf500, Ttimmeee, tankhunter, JamesRH, Bowtie, CarterCrowe, sheine, ya776, Rafa, Nigel12, mrperrine, leppinm, Mike_1978, fsp98, Mathias, julykus, eamcgill, wootmonster, Junko, noahyodur, Racounous, pbaidas, xenon53xenon53, Daiquiry, elm0119, Harrisramsi, gomaddy, Pyroon, ishipsomuch, bwright8813, Yanutopa, StinkyWizzleTooth, xTibor, Elward, FolarinOnifade, blasp, layognahor, DevWolf59, Treity, alcryst, vanlear, Symon Davis, hungryhermit, ThirstyFox, hds123, TheDarkLight, toniypedro, Rotifer, thejojobean, howaboutnoscott, HansLund, cataway, Konrad, webmax2010, KurtissG, Dannyboy, Tucoffin, KarBeyazd?r÷l,m, coolsombrero, thyson, khermerker, YoloYellow123, HallamMcIntire, BOZY123, Neptune, LectorEl, gabortoro, Derecus, hacki81, dheft14, machoota, BrunoTC, AnastGrig, jim2707, Lanrhartan, mkafzali, Quiztopher, SaxÈn, acentanni, Unlucky, gregor_patof, Jesper7, chapeau, JelliBean312, GAZman78, Willed, ATPro, aelahi95, latana, bo6, linio, SciTest, iamawesome, felipe.ellena, DavidKMurphy, jandew, Ralfee, kitty117, Kaavya98, jpatton5, soonspider, Bl4sc4r85264, LBlaster, Archaea2567, quiz, Theargonant, Gwidge, Kormie, osmarsm, Wojciech [PL], rachelriley, Ignis, Julio Jones, ich_net_du, kourama, BenPyton, josefine.folke, Anid, ChickenOverlord, leonamafram, celsobp, dtribu, Erik Samtmann, ZEENOEMESTRES, AdamantOracle, Max.1023, Onik, jasonprice, somethingcyptic, jules21091, joannaohoh, AwesoemLauren, Celeria, Foster, Krell, rocketdock02, Osuka, Angelflower0, Shika Books, kyle s, jentchai, poiuwert, mpajovin, puzzlewhiz, HudoGriz, Zebesian, Chaospro, andrei_wildman, Sherwood, jblaqu1, ebelaid, Joshashell, ktkat218, lisamad, Abby2012, colacadstink, odmir, GatoPimp,,o, pksvsengc, gbatama, Heramb Reddy, AlienC4, Cobby, Comrade LabTech, AI_White, gatherer, Abody, default917, mitjakocevar, d7u8d9e, Nergetic, MeTaNoV, Crotalus, Algotastic, Corranmac, dhinostroza, lightningelijah, Oreliel, kariaa, USCmwh, xcube, mlab81, Edsterman, Tunkasina, Rechenkraft.net, Szopen321, Tiphaine99, SrikuTheNerd, remerald, TheGinjaNinja, loomismeister, leo2, Messer369, Tiitu, Oukikoya, Tygerbunn, lherout, 1158499764, Aces 47, niashwin, harryk21, sciencegirl, Ijangk, Koboshi, woodie23d, fluffy3, Cmastin3, joeHope, hunter4928, mraider94, jmfish97, gatsby169, bonyonggu, LouisJacques, Nickbot606, starryjess, CreditOnion, tobi123, Simmo, CeraTheBird, jlentin0506, AlexR, baekhyun, mspsychorage, Midnyt, kafshari, Naoki, andreasstinks57, elpimpon, QueenDianna, Tyeor, Clindsey, postwage, ETERNALVLAD, sselion, martynez, BiancaLCastro, Amushta, vincentvegaa_, aria, Nol, dsands, mathcheque, ItsPiet, SLENDER RISING, LRpaul145, Oineh, adakadabra, Rudi Metzger-Wang, shaio, sayanbasu, Applecrap, mdodge, britsie_1, slytherholic, brainiacbabe99, highchemist, TR1406, motto53, Jac0_15, Mei li li, dGameBoy101b, PigletKid, jansonsrolands, deblagoth, CausticWren, m9898, Abatos, LimRim, Obscil, legohobbit2002, Dakirnz, ylu20030301, satrum, Omnicide, ejkellerman, gjneumann, Nephyst, uihcv, LauraMars, layerofgauze, pnhoffma, bchrysto, hangth, nickswan, wattsofbacon, jgreen77, mackinthehouse, ExtraKryspi, ndootson, AndrewBlunt, bootmii, brooke.breeding, spv54, Cerox173, euan.chalmers, atanimal, Pacificly, supertrol, Yuta10, J.ham21, Timus846, Beniona, EvilFish89, Heek, Hippopotamus, lleang, Zbyk, apexpredator952, Rojojo, DingoDongler, suitkees, reko, Kenriq, sfrail, wize, Spiny, Torin, LeavingGoose046, slegge4, cosettez, Yacuzi, PlaidPilot, itarannu, DarkSir, HawksinE, jlin0519, JQC1029384756, han.brzezinska, Brenda Geisse, doctorkasic, jzhao293, Kottosix, tolup, mishel, regi_the_veggie, nmengist, RemyD, B3224, DRNA?, Peter L Barnes, MJAldrete, Naqster, YoyoAarno, sdjili, DEdwards6, Tumis, salmohsen, robertodelago, blakjack, Marcurah, tank0221, Yeti, NervousHuman, Ragdala, steelbear, MrsMeeseeks, Sophieroo8, hchan, RichardMckanobb, virusvirus, MDokoupil, arthurm, clairebradley, soloi, APlayerOfGames, mwong537, mini1471, Miner9099, SG1966, elisatroester, cub4libr3, herenyon, emothershaw, m2oos, elstinger, krailon, achui7, RoninAsturias, syuan46, Executr, schalifoux, Piotr_Rywciu, Pandafighter, anan1003, sandman31415, s3tspy, altf3, biocycle, XTHANEX, oliviacho, Synthboy, kanayamaryam, bboylston, magistra, jsanhueza, WhovianGamer, Xander, jsantiago, ealohr, jomau, RNAribobitch, gregorymbrooks95, Zectron, snargleclef, bruhitstina, Jumjum, firegod954, selfishjelly, petrih, samster91294, richy3454, TerrifyingTaco13, kingkeem, musicmanley, Boslof, pidef, sfrye, whenham, anthonymur, jonathanA, Alberta6808, enorm, Remilia, Chemark, lime, momored, MarkusBergmann, mariocobo96, elpeee, Awesome Quest, MissGarbs, MrGecko, achant, darkhound, vickyrox2, givello, ScienceGod, giovig, Patty4444, MrFisher88, romanus, xaythor, jduber765, Neil1234, TheKing17, Glitter2, katalina2002, KingCodex, Brandonlam, blakesilver, gsing24, Wind2048, Miniburf, Digighost, virusdesctructor, dulya, Grallen2016, Spihc, tazmanianshedevil, TiggerLink, Zagreus, mctanker101, amberna, jafennelly, BuckledToe, mmxrocks, Crit, annaml, gravebone89, dRedstoner, Querty, TaGab, Vinara, lambdahindiii, lucas98, GilMota, lordnaanor, carmio, ozdamare, Malakh, gamma26, Racrbobcat, priyalpatel, maximegogo, damigapa, 3Balance, RotundCat, carabear333, strider23, lalashh284, Seror, John Woher, qwesdfa, modran, MathieuAA, harder.c, JX518, nshauffer, nanolasso25, siddhant gokhale, morphatic, fixed_entropy, siainc, unitedmadcunt, TiffanyStrider, respawn132, kiri93, Daemonsword, Atrix, Foliper, jpereira, PeAlc, TalonWing101, BelacF2001, mweitzel3, FelixIsaac, limenlumen, caranturiel, clollin, Bastian5, bingbong, Minigirl, Picses999, Ulys, M0X, ramulu, NTSX2O, Korosagt, Egavas, anachron, LockoBoomo, SamarthDesai, talasjudit, quihi, Dravd, kmares, rfm0905, Paternel, bt11262, heather.elliot, BunnyRabbit, crize, fromea, QVENTIN, PuffBall, TrueZodiac, Total, Cerox, blozeee, maxes21, jasperweyne, Fikoloko, fireasametaphor24, pie1337, Tikathalasa, Draco711, gauberti, Medk, jazzarazz, emily.banholzer, Yahuda, sataeeme, Denouchka, Lokiosus, 2020huntermerten, hmandujanoham, lassewe, mdurik01, drewbraham, aberry11235, imthescatguy, eternalbeast, chargyse, mcamp82, jfongemie, phobos, JimmyLy123, ShaheenSheeti, rsm080711, gtkrow, qwed117, archonxgames, LaCucarachaa, Adeife, esample, naylorm, Hex71, dragonlax, Stump Cutter, jdc, Instrumental, Myst_Nrg, cell(ebrity), sivatherium, GermanViking, DevonRolfe, stevasile, stevenw899, Bynmeister, Ashwill, Vyxz, Asho, modded master, allergz, AvalonKalika, Thomas Hake, kostyakozko, Jean-Bernard Tartalacitrouye, nlesmeis, ostorozh, eatRandy, Aberrant Drake, Mustek, Mr.Dale, dchen24, steelix14, Kuydef, greensun, BrandonMatthew, LHaines, SafinWasi, himateja, Pentrose, Lytala13, Supercafeine, JeremiahV, Thanemous, hallehai, jrau4, Jerdan2013, Lemetin, sachithm9, logan05050, MZA, azertyuiop, marmun, biogirl.lake, Filledagreat, geoffbrett, blin, Sigurokk, Kcfontaine, Futureman, garyhogan38, erland.sanborn, broger24, Coarse, Nukkunaana, kierstengodfrey, Crazy4Finger, Persephone, ItsYouGuysFault, DreamingCaliel, Eirin, Fernflower, Venderil, Minotoor, Luca_Giurato, 1SDAN, paulfox, Julia M. V. Warren, soulignighter, Headchange, Divra, Chintam, anthoni.noel, Caramac, BCool101, alewx9, astroblast7, ToninadaS, 212121, Deadbot1, Eterna Helper, qarlos, eladamri1, maddenh, alexwingate, Laura Justice, insinga5018, DirtyMilla, Marco St, thothdragonfly2, qtran22, jlspartz, nwu46, TheMoleKiller, rmeaney2, mharb, bgching, 83073, jxjosh937, wood89, dariki1, sonnyskies, hdong29, mfa, bsivajoh, CapralMandela, DettanKarmen, tdelehoy, arkad, coolninjarock, christopherdennett, crave4science, mathgeekchic, himmelmann, remingtontaylor, mathpathogen, asmit533, Jil4no, Elgryn, pvanwinkle14, l’ÈlËve bouche-pillon, jwhela7, oscarglo, Starflight, markmulligansr, Mippy, Baerdon, Extremist22, ottomaticman, stealthmachines, jpalme56, Patrick4444, ardavei, Nieljoven, nicer2435, roki.kos, Bernaden, Guillermon, element15, Kid_GameBoy91, zjosseline, smartspot2, bagelmanxx, Mairsil, yo_lass, masseli, iappelbe, Superd Sada, danielbelnap, laja4770, Drakorise, Azelef, hobo934, ScienceZoe, Anirudh231, radhikatyagi, osmunj2, sskala, ninabi, Tony.G, fluo912, evynbob13, eabrams3, nwg182, mdiggs, marsmission, memorykevin, SpartanJATA88, Dai-San, jujuBeans, Derick Hill, Judlas, cogitase, hnguy29, macclark52, Jenyah, xplocast1, Hufflepuff09, Stormcatcher, juhus, bubblecritter123, fuzzywuzzy2912, likegamess, jsong254, Leyti, E.A.R.F Health, Bethany2017, ssteph07, shotlolwin, chortman, Ladon, Akane_Akarui, kchung86, spowell22, Evian, Tchan, dsiem, greenshadow622, tharsade, ludojad, Kennysaurusrex, psandige, Archeopteryx, hakunamatata, Kronwerker, Feti, Remi13, bruno137, timlarrabee10, britfrog, rtkleong10, pbarbule, Uredian, awang263, ailuyomade, ylc193, Poisonfyre, izabellav, IceTalon48, Bankrott, TrenteR, himeji, AnkithReddy, tofox, AKown, slilienthal, abellin2, PDelly, gdrakes, sleekloon, LidCoco6, coolstar1611, KubaZgÛrski, laiserbeemz, shoeprich, JC2000, kenny2816, rlam73, xXrakista_nerd142Xx, aweitzel, NathanNguyen101, human004, littlestllama6, arantza1994, karniko, mattman571, tyster5000, xenon54, T30TW4WK1, jhill, davidivadavid, Gnik^llÊ, alayton4, sstaines, cicatrixes, hunterlock, David Dockrell, Ceazet, esiegel5, ahmedrassmy, Ferric.AU, ernewlan, Sharkkat, ewill19, mfourcand, Species5618, Arkantos, deltragon, Vinitski, Nintenkip64, Shaneeae, jcarlos6, lsjumbfan, dhdnstjr16, hwang584, PerryF, Margaret Dinon, dtep, gatoporonga, thesepaperplanes, jconrad, tonytony236, adimbleb, Ebenezer_Shark, gguido, bwondimu, gonea, Ultra_Lord, AchooBlessYou, eqmb, Fozilla, StaticalTech90, IMHELPING, Voila, grazzZapper, Recnalc, 17LuxtonFT, Mikeay, P-Criz, melindac14, draganfang, megansch, carpanellia, queq, zsf, lizzie1111, Grayn, aduncan1, joseph.cab, SathnaMoriarty, Felis_Anxietus, cozmen, clairehebert, Mike_w, alfrick, amckay5k, bakufun1, ingaffa, Weaslet, biomadness, coelhohunter, msalter4, nevillelevel, ruicabanita, L_Omar, bissa, Garnasha, SudoDave, rnahh, zwilson2, Polert, Smazz, JoveRNA, jmanlavore, aidenlkms, IAkumeI, space, Panoca, salcase, yanggirl1, fizzy9405, MisterAtom, ipo, Iyansa, Richard Hade, mrippen, sspeterson, N0inimeht, kho296, holyterror, pantushev.denis75, fffffgggg54, Domestic_chaos, roamerkill, YKTNSloth, Mennoknight55, gpspaceman, Starkyll3r, HiroProtagonist, crazyworm, kroma007, weak_chris, kruemelchen2, Adarsh, aresetar1, Anderps, TyrannosaurusRekt, Andyrooo32, sroffey2, hankpete, shshrey86, KHeyser27, Shraumanugvar, josephk545, FrederikVds, Freeboots, khamous2, Tudoman, Ness, Ethlaron, jedikidd, cuhlemann, xx, tristanmoon, zeranosdepozidox, Caeso_Lucilius, Cake_2101, aaron connors, aegnog, rakshaza, tmilano, plumcream, saimi, AuraSivan, DonavonEllis, Longdart, AverySebolt, jrikeman, natybob, faizkids, lokiu, woutvw, Smooty, emilyrush, WafflesPancakes, Pigoon, MrMeep420, rlblackbelt13, Uber Cheese, jkiestin, tvargo33, DrewWallace, Hollandalle97, mollekes, mliu334, AdriComb, habouhus, pythonql, ioob69, TNuWu, Donimo, nathan34950, Xeonicu, haggi, JFAN73, DeathByPrograms, Zint, odaliz44, MoleculeToad, kevin rose, telomar, biomeh, joen9833, wkcason, TheRedJuggler21, frootloops, tjohn26, bliep, Helix666, dowdyea, EX6578, FunSel, blanxfg, RoadSpike, pranasag, sarren, Borio, MDrucker, Jaunticxs, Mich4, FallenT, akhantha, C0wAssassin, XVCRT, TablesWillFlip, MAGNETA, leaveyou, Wafflesthecat, Maia4444, Shell, francesco215, jnorthe4, Sam Pettersson, Turtle1331, ecate, iswaffor, cchan827, j1, keeratsingh, Athanacia, Arthuriel, Minia, hohnd, The American Inventor, DIv35, daviddb, ferdsstyle, Jeeri, David_SK, Jollygreen907, OpenTBang, Don Lion, robertweigman, Tavikins, Max Tee, etexero, Whalet, hseitz24, Merivos, misterslim314, ericzuck21, riBOSSe, kcirone2, umpufnufguf, alpo, Geofly, Jfault, uddyn, tshall, shuan53, tmadge, rgxHost, vcarter0, Ccile, combatcello, timetuner, immuneinsanity, Toxicsneeze, Saotao, ross713, EnriqC, pblucas, karolina.pietrzak, hauterho, creator, PoppyScience, redstonator, ibatko, ZulaZoobRNA, bea_candy, dsizov, SimonWM, Korvin, unlucio, Aiden Pampo, Irthene, raipas, 6e7c43486c47723484e26b97f26dd7c6, lmabrey, OxygenFerret, mmcgil8, poochiethecat, Critter13, WiddlePug, nliv6178, ssengupta, seed_alive, strickerrei, 579365, hnilsson, flesh-pocket, jhagood_930, Erfevan, Wholesale, mot24, DERPSQUID, nsimon7, Eternamolecule, davidbai, dmclach, adamg, simonk052015, abriana03, furn05, bflaher, ETambor1, mekito, viettran1234567, Pokemongo, MiloMaeRory, .WolfWorld, a1rwalk3r, Noah Linning, hurinthalion, Rishabh925, kyle bu, jbowers188, Kunstbanane, Ispanico, bhgreenb, Nhali, fff, shays, agrow2, jlee2854, SLilienthal81, qnguyen, KatelynSomsamouth, 3than8or, Ardebar, CantStopHodoring, cas, ABeyer2, charles_a_rose, EldrichManilo, romanedevaux, Jashaw, Mcke, cwilli31, CreeperGaming089, ChillBIll, pcollins24, hellome1, katelyn.mose, bruno, 4science, fleshcrafterAngel, XeonTesla, esun7, tilbert, paige.roberts96, Algorythmis, syang, isaacdavid, FreeMars82, Picten, rezten, Trinity_, Krejberg, Flygunn, mbarsan, AxerHi, Zander, asilvag, MayMay, Gared, sliptup, enomised, Volodymyr, Danbi6, fghfghfgh, andre_cortez, SuperMerlin, yhper, SunkernSciences, Lhydelhart, andi_se, lderikx, foldingnoodle, cubedude47, danass, shobmann, snoddas, haynerg, Boara, BLUEraven29, adri0115, elad, sientist, Caramia, GinoTitan, Anseth, curranoi, dolteanu, bng123, ncase2, hvo12, Metalval, zylocks, karenzhao, pegasevil, laracomn, DAHW, zzd1992, Evolvedgamer, KimBurlington, BlackyAlex, ShannonChem1, dapbrown27, ArtFreak17, Alvengaan, Prokaryon, LauraMcMahon, Spiffy The Chicken, mmindich, ydayani, fiordes, kchoi8, gentelman_bear, maheshvi, heather.lee, Carole67, gabrielmm, dtk777, Bibriss, Patrick Cole, aTamSmasher, Curistel, RN, AlinaS2016, Migdalin, connorlink2005, bentmaniac, Spaceboy685, Cypherfox, rekorn, KrystineWang, LlamaLord, halocarter721, Thud, violetbrina, Postal, khobbs3, Ajarn Chris, Einstein, ellenofcrew, khaar, mangoninja, Deafwave, RespitorySystem, Dank, kalserules, cmazzi, crasher925, aadelstein, ty513814, ChristinaMetoikidou, thatfie, knj0610, ocelik, yzarwehbe, mbirds12, mintuk, zzjay, spovolny, Abena Adjei, OMJPC99, fireking16, evalos, Vancha, Demongate, jadeshi, lordcamz, Gerardo Jara, eevans29, anguye15, mariapproved, maihime, iyktyn, thealmand, Hiefel, ImpulsiveSpoon, mebious, slashghost, Lum4r, Jamie4563, Edginda, sbarret3, DOUBLE, Smachar, komo, QNVP, Purnima hariharan, noep, anvimar, GCLonginus, FlorianVO11, sllaporte, spmolloy99, awong234, CC333Junior, mliu366, GrandmaJulie, E.Blackadder, peter990011, marora8, MrX, mattia4496, AzathothTheGreat, purklecancer, d-wauters, elizabethallegretti, Xalndeer, ezhang26, belnoah, Nicetree, nvladici, XXBlankXX, BBH, tungakl, rpeet, jvmm18, PseudoLW, Xx_WildSmaug_xX, Joyce McCormick, hoxxymama, UnunoctiumAlpha, Geeter34291, SaninaVelee, Hemia, sirischuck, Omnish, ebergstr, LBFANAT, joantaveras, amberjp, Roads, TheBigFish13, MK38993, Tosatertopia.com, pnguyen23, profemoy, jobob, Cryogenic, toadyjon, Reli1218, Nikolai Is Awesome, egosum, novoid, AtmanHenrix, Dediles, k96as01, yhuan373, NathanGrimm005, momo12589, israel1234, flo, gratzasaurus, KBinauhan, Kerbalnaught, kamercani, ntgramstad, tarimo, fahme, M.Runions, Thebrosblack, flugga, nmhossain02, gnnop, johanna1904, kochieng, DouglasUppity, GooseBeef M.D., g5nty, Oogieboo, Jaggertay6, SoulWolf1227, Juliette, Sardinefinder, jackthebest, joshi.sumedh27, pmm142, ionut.axenie, Borrth, Rasend, tng92, katelena0, dcasey, nim579, stu_xnet, martindude03, sgilpin4, saksoft2, calebgeniesse, AKI_RAinicorn, lobotomo, TARDISflyer124, lmcfalls, GamerGEEK123, rocktvs, Britney Ton, Nathan_Kamm, JJREEVE, KeithoBurrito, hannahgrace1018, dthong4, Katokoda, BangoSkank, blazm, irnmonkey, Stranger Danger, zordonmlw7, Brough, Dr. Machinator, marbar, Tettecareca, starscape678, Archer2150, steinso, Bastian_5, sb12, craigc, anguye83, Miss Mint, vturnbu, svoorbe2, ^_^Anna, fionakennedy, Maximeus, engineerjosh, jray21, brookebenjamin, nsachde6, Mlg20coeur, tabu12000, kikiavicenna, cemkalyoncu, justincjohnson2, ObsidianCrow, conraeli000, w1ll1b0b, dannyderito, nycari, LiamCampone, mds8575, venemab, kwade4, accessnet, braidenphilpot, odrae, potatosalad, kidsister4, FarmerJoe, Nobody21, thopwood, GrannyHQ, green_panther, Zashiy, Jion, allenmalo, kalia_kona14, matrtin, ctobita, rusty2hip292, G.shelain, ondradoksy, naimahanderson, achen325, greenjac005, Pisobiech, kalleina, Dt615, borderwulf, sookie0825, ivybut, Storm_Trooper, banglesrok, pkriens, Razer, mchopcia, Mightily Oats, TylerFowler, gayounggu, xwang985, fabrioche, Madrawn, ShovelMeister, kint, jchingjo, rizabilgin, djdewott, astonian, pdeioann, Lawrichai, Stevenador1, Fractalus, PedroMGomez, hamster king, dsteph25, aherbas1, snazara3, azuredragon, thojbaek, hinomi, soha, Nathan Graves, ghty195, moonflower7, alex333sh, Cong_Nguyen, FifthDragon, Lab man, ehf467t, mlgmarkmlg, tsubaki, Opticulus, ToriLee, maungj, leiak, hlefors, alanlam124, switters777, YuukiAira, squarelles, she_elf24, rgoyal, Hruzicka86, silverlava, Jacobbart98, mvw, alonsoaguayo, nihaowangdong, L. Lin, VictoriaP, aussie114, JaQuez A., BloomingShadows, Jozmo, CANIHABEAPIZZAPLZ, Vis3rion, mhrussel, abrow326, Wajeeh Chaudhry, oreroe, agreaves3, TrueTail, gparmar, ashrk95, fmushfiq, Shriya, dkim553, marcucho14, wldmn13, m.mcelroy, PyroMancer1357, piscesdan, Xiexion, AndyEteRNA, acusano, dkim533, innate twinnate, NoelRoche, Rigatona, RSO2ID, levibalint, BF killer, superduperman, GypsyFlip, smallvinky, Random-Weird-Person, Otman, KokiriAle, jun1126, Kuhno1980, Thegamingsushi, sfstika, donny123, Overgrown Lizard, thomagra000, cerd, kodra, MrFisher, ewohlwend, xtox, SteelCastle, akhalil, baylw, Kirbypowered, Pineapple2Face, dimozg, Haku Temaki, Kang, pvanhook208, atributz, Silmaril, kizmit53, annikac5, blueivy1117, Xephum, bentlogix, Maximus550, mihnealook, EmilioSevilla, Caroline2334, ≈rgangsdreng, @Scientits, S1961866076, Cucumbergirl, Daholsapple, epl500, adouglas, knighter1, soybean1990, Jym, kbozek, sarelg, neonspacecat, JeBuzz, nicamarie, sshans, audio.01, mcapobia, jblaqu12, Mt. Vasuvius, ralichte, SlightlySly, Matthew Kuan, Crazylady, genops, aabdal2, SalCoombs, thefallenone2204, afranc9, skaterkfbs, jegarfor, aalassir1, Tigerlionwolf, diroku, echig, genetech11, alang49, AlifA, orangeninja27, zork2004, alex02144, salex, TrackAttack, Justine Staley, sdelm78, MWeber, bdk, Cathrin Lionhearth, Gondar, pumpkinpi, Almanorek, chefshawn78, ForzaCar5000, Nga iti, deadlypliers, magesparrowhawk, Askapa, whiteflare, mpesce, Rowmanow, mcgrawm, ouv, Farming23srx, Gilchrip, billiscool, ryanhsnell, wxy21, scottgrant92, jdemse1, zareef2003, Sturmrufer, zeloui, kittey, mayanightstar, moleculaire, pogi23, mghaderi, niccanney, snath8, elliezack, WFoley, Twisted, Oogama, aadeoye, freedomforfreedom, Haojerr, JanH, slazarte, tuxmain, rutzph, aisoize, rnbetts, Undrscore, edwardtan, Alex_Logan, khulanu, esavier, Fyrebright, la .lojbanana., Obstmonster, NotaSF, Chronon, fish97, skim975, isiloron, jethrowu27, alahmadi, Jaepheth, DanqMemz, CrissZ, gameplayer888, Iralix, kittykat1, Verbanderbog, krawson, MikeDD, smenon4, Plasmarocket, SeanMullett, stefan_nikola, farfromunique, poundsofnothing, RichardMark, Caeruleorum, tilen1, hessexpress, jmichelle, GMXMithil, e234, Miguelython, shackelfridge, DavidC21, paulneub, cpod22, AgNatishia, thuang92, veritas123, Oesophage, NerdCat, NikoKa, tim.h.kostolnaksy, Zomboy206, CVJ, doverlg, Smegenys, bekeep, aliersan, sgould, Floof, XD9lizzy, ScienceSuperhero57, nathannau, svantri, e.pitchford, lukefarns, mugetsu13, Deniz Mersinlioglu, dtsyplen, NDDbest, guybar133, lucygettinger, vizo, astarr8, eS5e, RolandMoriarty, BehNing99, Churmak, Joep 68, Final Warrior, mjychabaud22, msohail, Catecapp, jeanagustason, AkumaDono, banaan434, hetlock, lipki, MaitreNova, mbaggot, chippyweekes, Kotakiba, bsmith, tzohar, gabs999, TH3R3ALDANTE, Yliander, Nekkaria, Bethadose, navtron, nalkhish, mg1073, bnowak, BeRod93, joelthetroll, epond83, goodandy2, taylrd, sorr29, leor simcha, rikpat1, miuds22, mrok, IHelpYou, luckyshot68, WhimsicalDickensianOrphan, aliciacouto, Desolated_Dodo, yqin48, pmposey333, niniel, kalner123456, foxxy, k3lly_t0dd, Elcocov, girbobjack, brennaja97, 1ayeee.jr, haydutraduke, kolohe, cobonator, Colen Garoutte-Carson, salish99, Somerandomperson, willbeing, mpierc9, dnorwitz, bowlsmoke, xnitori, Martijn Klijn, luiggikf, bsba, Michaelhess17, Lord, julia_grross, missmmoonie, aoteabroa, Portaldominates, toscano22, JordanDave, Aden2805, BlerStar95, SirOcelot851, Doakungfu, SadCl0wn, n123, Chadowcat, osimakov, pleco24, Kenzo01, hackkpo, carliem, ChrJohWol, Jakv1, mr-kim, Xan422, john4563, hardy3, gratitude, RedBoober, 94fdrx7, xXjacobXx, hahmed, 01001101, Terra Master, Gadge, kolosova, muh42, Jakob.Gibson, Darius59, jrp317, adeprano, EFER, mantatail, kazafox, michaelkay, sanagnos, togg41, Psychoripper, chrisceo, alisonpurcell, TheFlashIsAGirl, MÈsura, Viscorp, AprilOlaires, fder, jayfunk, whitmar, Jezzaboo, Ubald, StrawberryWaffles, banuorala, jerry.mi, broadwer, kkaminsk, pdmock, justmeemily, artemka100, saversa, Binusuairos, 602, Spiritgirl, nforesti, curtipau000, pfudor, kluball, QuantumShaw3, aliu223, silerka, nawaalakhwand, aru54, Gabs_the_lil_protein, dan german, bmusselw, AndyJay, Jwb52z, raph.desage, badcorvus, tata1993, kover00, mlee688, aboubou, cleveron, Xydron, 113568343, Akkaneko, Heysun, khendarg, hielox, arelyquezada, mochi, eefremov, bgburke, WEenYThePOoh, bmosley00, raspberry, BytesAndCoffee, Crimson.Rose, Nbukhshtaber, addewyn, jrosati, Glenn T Jakobsen, Onk@lo, JD186, sjcook, Commander, Tanakakuna, adefelice, minabenc, 15boolah, SlayerIssac, Echecs37, TheGenieOfTruth, amelia kracinovich, tonemas, shong253, Ps1, jackgbg, iforgotitalready, Nikhil, chronicAnimator, jogundipe91, ejoe90, eblick, keithtothemax1, woodytw, jazoody, Spicypaper, mspeight1, master_son, OmaroA7X, ncastaldo2, Adam Grieves, ThePetarda123, Tannet27, FireplaceScience, gotauber, notJames1001, Mattkew, dager, MarioThomas, atran94, isabel11, pbrazzthemighty, bbridge4, forkihn15, smir26, Trotski, faditarek, sylviakinoshita, Nick_Linux, sharkguru, Theoneandonly1040, ghlee1219, KatrinBlade, Talifmo, Tyberio, rachel.phan, rickatello, isporrih, kurumiheartt12, agm783, Azor_Axestrike, danimal527, TurdusMerula, Halu, RamonH17, Khaysis, Stem, Grolsh le laid, rnaklask, nikzaq, Hugoale, stancmas000, ncastro, Ornjpeel, Ruben24, nono_you_dont, sakul360, wernergouws, mimeep, ceta, crayolalola, IVT, ashleighhopkins87, gooper20, Mojok, BulletNick, drklimow, lpartida490, danascullz, ecowan9, Young_Ramenz, ecd1192, ami, clemecar001, anggasendiputra, firedrake969, noahshawnpower, ellieg, Generaltheo, IsadoraGenevieve, madakelij, saustintaylor, julien-elodie, kljs1972, martieth001, Capnkrunch, ragingasian36, bmj1f15, Hourly Kind, SEBOMBTIAN, JKOtterson, Alejo87, Shiser, c.campbell22, Ilovepie, adamw, Mr.Smith, blah1878, doc jon, Hadeed, gent, DaMaster5002, ilost, HenryMartin, jyao1, kadir, Decken, GorpGorp, Eirok Burg, nyren, irresponsibleNinja, eternaman, greena81, cameronpalte, Valentiin, Ajgreen04, joseph a austin, Zeetepez, HAPPYNAPPY, LailaTheGamer, Mamadespik, pmock, maks.bond, WigglesMcMuffin, eoghainam, 21Martan, courtneygreen92, linwillis001, aubreymorrone, XxX_YABOISANZYFRASH_XxX, vazquez8, TKAK23, Susan Gaughan, Jovonndac, BrandonVan, Egodeus, Ysoscel, Yolonda, Biotox, ssakhi, whosvanessa, chunyi, eprescot, 7g3p, downinglibby, msmaclennan, flyout7, kougaro, arxidia, memgineer, Jsenecal, shamadac, Kingsteven, _147, adamdry, Nico Scordakis, alexjansink, element108hs, hukey2, alramirez1, silvargent, yarmin, kbwarlock, alexandreabeh, Camillou, pedanticaardvark, tiikerikani, aries713, Aj calle, klaverjans, Funster10123, jbuice, shb, vislous, ElliottB1, Taco, andreagarnicamunoz, natedawg49, mishfa, Faith Washington-Law, jkmcv, Devy, ayezur, sana528, davidsbond, cbrain, DorinaJ, HiPeopleMC, 0utbr3ak3r, Pianisssimo, Lenchic, carugo, a4368146, plmcr, MoonShadows, bazooka201, iriswave, tiredguy, kwh, 578780, julioflammenoire, theIOis, ListedMyths, ahysick, notyours, AstroPeep, bmillway, KimJongBill, Nick.Trovato, cybersun, Samantha_C222, moa168, Kyubito, GIBRO, Manarchist, kamisaki, ryanmcstylin, DionGames, skrent88, MChiusano, afsahafzal, Nessa0311, Osiriskiller, 22minnis22, voyager1, amathew, Kayla_Stevens_Schmidt, tumorraider, ccshnitz, mzhang5, trophe, Gracie.com, justabiostudent, rowaninja, BuildAnkor, queerferret, mcwasser, aagnes98, oman365, Juts, c_danovich, Evyslwyn, nick52a, JjUuLl, Jlim, VsAcesoVer, marcoleandro, TeresaFF, juzeddirna, ArtiSci, rybosine11, Dornito, casprus, felixsiu, B.vallat, dantepa01, edtecot, Megan333, zezice, Ribocop, Alianwari123, ABGriesbach, leoelbarto, linaha, Naomi Feldman, aloha.natalia, tberkness, laynaaf, FanderDander, fatimakameh14, lilfish, arlychay9712, Ionator, jordan12345, nasoboem, issathing, Augrey793, Unbeatomer, Kejchal, Crazyrice123, AbbyRagadio, pascalblue70, rapidfire, Goldrew_Amp, lucyjiao, Dawnsky-The-scientist, schmucker, kimwue, LagMasterSam, Apekatt, ScienceBroughtMeHere, HartySparty, cwanket, MontanaMuloku, Landshark, nofundamental, Cean, Angelica20, Alleycats45, jdmclure, 404random, Nike114, Duckman23, mi036, rider123, ragnork, Najimac, alina28V82, SLayGunner, BluntDaddy, Ibar, dianadockery24, scaintech, hiramwainwright10, BG89, shettler, Maxovitsj, belugabcool, Mattias Hermansson, rdales5, Joanora, Caduceus11, RiborigamiMaster, Msmrme, bdsoccer29, thedistrict, nbiley, DryToad, Ray0708, Calamity, emondal, Feardred, Leo_Gecko, breku91, M31, Kanta, Kyottee, nickyjo1999, desmond, lappaja, Harosn, Afterman, TheDefier, pioup, Zachattack525, lori_n, oem, zquoid, sarlamea, PG, doglover, juanjon, daniyak, yxia87, gecochran, ssantos4, Donutmassa, dave342, geek, kskalski, Vr?ina, HarshuEdu, Dmitryser, graceelizabeth, rot-proof, anderson207133, Error_707, dthielke, Shad-Knight, Sullivano, Achieron, Kayla S, zbjes, Neo201, TacoDwarf, Tyromaru, whitman5, MadScience, ingolemo, yutuyt, CloudCompany, blaize2377, jeesanchezdo, xphamster, RadiantDreams, WayOfDro, Northmen27, Blamishs, simon1094, teseting, mtw687, raman, Spaceport, nhanhhf, shahrzad, lbahmany, seanickel007, RomanKantor, paola, OchoFlames_808, mahdiassari, darkangelx, User42, polluxxx, SammyLerma, 341464, Nerubicycle, Agnes_R, luckyclover5, Gumili, 12lucyx, bizcasobes, HenryTheNerd, 3030, clockworkthinktank, linarie, aab70, lcbvianna, Batagang, ozzzzz, fedebio, samtuv, floodcam000, Lorathor, Wilkis98, allyrene808, Vade1234, Jonathan.Socoy, nateaross, ehollan3, mregnard, c_to_the_c_, regfish, biomadnezz, florashaman, Tiffanny, DubbyWubby123, marchisi, Becca Rose 2280, Jujuu33, Echuanga, Medicine_by_video_games, Feetzee, wchan123, aghosh12, skaplan1, Oesirpfy, rosie316, sof, heliotic, farkasosky1, Burcin, KrizzNA, VirtuousCat, Gonz042, zombooty, hanto, plat1n, L.Hoy, Aotearoa, friis120, Wojtek, Raymond G. Feilner, Jr., parichita, VDoc333, dougd03, skoff, ET Hazwire, Edison Amery-Zwartz, ThisIsFun, Aaron9101, Lis_atic, gt1234, pakstrax, Melony3, Katarzyna.Kwiecien, kreol, Leky, mcbride, Rhapsody, gsat, superethan, person_, lallllu, Sycamore, graenor, Noobix, hjohns29, leana_mindra, Duikplank, raitan, lililia, ggemme, kademing, bgw5064, et_et123, vincentkellyy, ChrisScientist, J_Starwind, KHSMA123, saraiispoppin, yasir, PRH, AbsoluteUnmei, nickr312, jnchappell21, machovec, Naputi_Today, Liloute, niv, stringtheory123, Rookiedu03, titousensei, DARE2, maxo, Smitty881, rowan42, litltex, Polling, jpark189, 0BlueBird, Plip, svillagomez, Eagle Bound, jjaffer4, FixTheFernBack1990, pab1990, cashew22, khouser4, byuan, johnbob, Georgia514, loyero, Bu?ka, emanuel213, Rance, cyberfox, Ekat23, blizzman1999, vladimirceman91, hedi, earthbound9, thefool25, Piggy, khanlamisa, Rebs, SethApex, Leoco, danielmcgee, JudyXu01, Belac, Loic2003, Boomshroom, bigredmadcity, EpicGigglez, Ketcham26, wotan8000, Snakehead38, Anjani, DAX5919, kackley5, tony149, Drunkenrhyno, Sarina, misa, Livogom, scienceisamazing, Yolo2012, JBogle, goobgo, cblum12, kpines, columbusinaction, criolongg, EeveePoke, alibobali, boltblast99, agrxkm, Greazmonkee, thematinator101, Helkost, zhto, nitsuje77, dinakha, corbineau, yxu442, imedeiros, may266, notelijahwoodsirl, rkart, mcardillo21, rsfmaquinaria, et246, luckypech, Sniper910, Lord_Aleptt, fydpfg, Wendig0, reyweld, sgochsmann, EagleEye22, christineortiz10, marhwini, Cupa, missivibleninja, noseinbook42, Funtagira, Addock, bunksteve, etherealducky, angosu, OrangeYoshi99, dicegamez, muser, ginaT, mcf, fionnan dunphy, eternaLwarrior9991, SheSoAlyson, orangefriedegg, islamgabi, defrajac, Skowyn, allisun, Caomai, Kgomo, ch25042, Shad, Whisker2005, hasard, Calliope, mustha, Leafsparkle, Cougy, draker585, Valdis, rupal totale, t0903591, lex555218, Macro_Lens, adroitPrognosticator, yanak, Lias Hosmer, cooba980, nelover8, ballin_with_daniella, Sebcarotte, shinamin123, NoÈ des Diversity Guys, jilly1075, lgellerson, Samworth, ssk, Lugs, mlowe, stepman5, tiedemies, franfrankim, Brennanscore, Shaolinchi, jiffy20c, jakemhyatt, MTaggard, sergiubreban, Link Shaoran, Ovorion, Fawkes, nusratnobi97, drgn8d, skeeler, akiraaaaa, Buttergott, Vegetableman, AUniqueUsername, The King Of Clash, weltonal, savanh, Tutur801, Anj100, j.barnard, Xellonox, nongenderedentity, dv_10044, jasontong, abehere, crystal chang, phynx, Martian_Boat, Nohic, Gamingscientist, eloipieux, Elite Player, geogeo21, mornplus, TeddyBeerSebas, cburkett, Zotshot, NelsonHMarinG, notdeadyetbob, awesome5008, StrawberryGS, origamer, davsalazar, antillon-beatriz, ant21, sophie.morois, elliottr, aliah.o_o, Lena E., Pedro Vieira, nodnarb775, adam.spiecker, DNA Master 007, svanduffelen, brooke9819, soadreptiles, kikidy, AlainFromEarth, armbreak, dtmclachlin, jp1687, atypical, o======8, mgupster, julianguyen30, Jojo2016, Haydo04, mcklsk, yfan84, Kumimasen, psychonaut, ventusvibrio, kosmasg, TheBIggestViralRNA, 09Q-SBN, jamiebosc, Proton, jeff0s, EteRNAllife, csziegler, Stefan C, ninja157, Johnathan, Jimjunlop, smoore96, wyh, AgentOfBolas, Habakka, ptolmey, GregLam, basseyabit, TroyW, Tribble_D, TheLeahpar, dimadan, mat20002000, Ditjedelta, Swinub, os10, Tunabun, shamrockp, Jubs, hreaper, Schmeckles, ihavenoideawhatimdoing, tiberius.duluman, AlThePlatypus, Xx.Wolf.xX, BIC123, adalel, Jovahkiin, kimberly.b, anton386, tnrlatt, Jesse Pinkman, baruch23, nil, phndl, andrew3511, ToasterBoxx, mchui, zarry, cayuni, Hammy350, Galath11, Bruluh, pvbookworm, gentsilver, dobster, Mgb0, Spug, CSteel, Hyenabait, Maxine, rhymeswithsnake, longtrang651, benpaduch, wonderben287, Asimov_The_Great, Marvel, nfried4, Wirvend, Immortalharp, andrei.ichim, ricemomma, ellianat88, t.asplund, Bilsh, gab21s, casare, kmfm2005, crazed4now, Jack Vera, seaslug84, Knoerdy, Curt_Folf, domah2, csfraley_rna, jacollier, daniel sun, abcdabcd, jwong521, bitman, Ine6, johnson.o_o, brandonta98, DarkAndromeda31, leian79, krismuasa, bjorncal, thesaladchalice, dakkaredsquirrel, autarch, matthew12taricani, klee667, yourguess, jsisidore, zellyana125, Bonojora, Fabuloso, Thunder_Bug, thead, enakas, ralphjin66, yaya671, ldorn1, Random Guy JCI, Sebaci, Lauren193, Kairos, gattler, Pumices, Grimwind, Zoynmkxq, otgw, sea7nc, millerm10, Neuo, elsyrin, lucelletanedo, idama, clone909, FinnZabaar, Frostterra, Hyva, bucketofsquid, crowan641, rbamhd, patszy, shavera, Max Goff, mariecorradi, breannamarks1167, sarahjihae, Pwoness, bonkks, ZyTr0n, strongbow, birdgenders, YeOldeShirt, 123928, jessfrancis, Gii2000, caitlyn.johnson, PishaMeans, enrA, kharyana, Mr. Snake, antosi, QIRC2LoW, jujogio, djproctor, noodlesnz, wstolyarov, Mehal, jonu_the_best, dennis16, sirrandom, sahmed18, hi529, bookluver132, lissisa2003, Bruh.its.rheigna, KylieDam1234, ptarecte, Eric IV, peepeetee, shawnxm03, mbohola, liaha, ztron99, JackDBass, yolao217, Post Script, polko14, thomnom, MikeHatfield, PrettyBored, kapkaart000, Cassie_C, esoryllom, Coppolino2002, osarker, TheWeebInTheBack, RNAdolphin, dtk999, chorus19, microbemania, Kreaturez1010, Stogsdill18, 19newcombb, breannan, jesscheng, etheenglishguy, SophieBooBob, Chelipotamus, nayfann, aurogar, kml, minjimlim, lixa9, Hannah Demers, HÈctorThor, lucaPasinato, Youlius, jcapp45, khiser, domvalc, john132, 312, Belarnon, jamie.bosc, Kaaktus, Sopheak, skrphilip, HyperLan, khynes, sbwtt, wow, goatboat, Belsinger, Mike_Eterna_Science, CriptoRio, Catt27011, pifpaf, Doadoi, Stanic, lgrainger, 784328, jpalmer44, ringachi, paytoncox23, BybyRONDIN, John_Benton, yokotopo, zvit, Vonrek, hackstarter117, hypatialovelace, sghosh1, RunRudyRun, brooklyn.kelly, dnanua, ncortiz, RonZimmern, PheleshiaPersaud, rakeshsahu, vtsongwriter09, skymagee, holdenhk, epbryant, cjmgames, mousegirl018, PÈpita, Onymbyte, profbdcohen, daanvanbeek, Navigator, lucien38, Trevorscow, WD3k, tojenanic, 1433, anakarina8, ViJay7019, yasminej18, jo.karou, HMK, lofty133, gurleen, mscollier, BaptisteTesson, CDR0420, Stalk3r, Matalos, hopalongsappy, xmoopie, Vini0101, randomblakman, bioismyfave, alisenchung, monkey133, basiqueballl, numpk, teresabg, jenzhang097, hanman69, miles1000, JuarezJac8, shorelines, 118lf, Farrout, Fealhach, mikkamikka, aconant, SilverKnight, ComfyBauer, Neythos, juditbestPl, Draksis, ilovepickles.com, ILIKEMEME, ampo92, seanzhao, caeleighxander, chelseajohnson90, rugology, jsosne, jorabanzai, tresus, ExtrementG, Ederporto, wraszowixu, malijah, bulletfire, pratt.williams, Jonel, luki99998, TheRedX, john paul, Ficik, GrassyInferno, jrogers4895, bsmall, Mizerable, Batpunk, masonsyd000, KirwanC, McRapperMcC, Wilfried Vanhees, Lhiebert, xxpaulob76, Aid11snake, CJohn76522, mufintime, BSLangley, awemanrank100, me2cool8, wingfly1234, shaimaa, Nomar, Durendal, dschwartz, jb1516, zobi, daleonar, iamgarethj, 2530289, Daisy_Of_Doom, someoldhobo101, ckirkpat, Alexavier179, Matthew.Nicholson, kilposhi, caiti-erin, Dr Pedro, pipio95, Sgt.Pepper, nizamra, Sherriff, zgordon2, wilson3529, brittstja, anonmouse, tydigame, mmetzger044, Edman, frazom, ketepi, Deirdre31, wasd19375, GigaDon, beachybum28, Leonitis123, donnadanyali, 2nd gear LUFFY, Wafflez007, Wintermute1992, nathaniel111, erina, emesse, zushakon, khallen, noahpinion, Geri, 1577, sohumd96, Stylish02, arempson, timlo, janel1810, Antoinedino, crunchytime, Juiblex, xiaoJu, Hatmic, Cr, Marvelously, sbadawi, omrot, IronDogo, cscott, Bugboy16, RishabVadvadgi, knonymous, LiL ickey, YrHopesAnDreams, jemrick1, Daedalus89, Tayloreolson, klovem, mvluong, puzzleman, EmilyWalters, chimptastic, 564391, gc717, Alonso, bivitarttus, rasmuskj, Trandel, RAVaughan, gregoire frere, cjelicah, idontkara, Anonyme, MABSR, Joanthyaga, jmaa01, ZacharyKZH, xenowarrior, julianguyen, boccitanus, scarysherry17, csarmiento, korille, killerlight, FaintingClown, YankeeDoodle, Darklordteddy, FRU, shadowsail, quietpetrichor, Anedlit, ThirdScenario, cheyenneconant, anmenqing, lwingo12, l2thompson, Brand2270, Dan danny, Dark, RenÈNewtonAlbert, xcombelle, stav, jneideffer, Jjamm54, Warpaint1263, silverust, tyler2002, werebear, Noshahaw, LeoBattlerOfSins_X84, vijyadrit, IrishRNAmaker1, edgar21, Seasonhold, Firedragon1408, RetroGameChan, lorin, Matt_Griffin, L0gixIII, DR.LCI, Kianna Mayfield, Namoon11, caverdema, alphadragon, ethanv, Veretion, JasonL, Linaste, lchamberlain3, graou76, Sophie16169, CoreClock, simplychrista, emmaline, gnclaus, winner, Desap, vm, olixx12, cybersettler, prestonholley, TL1, JazzyJ, salesslie, msells, edgarm, cspoonts, gingerInferno, alphameg, rosettejaneubalde, elB89, york84109, marychapped, eKTHA, griffinmic, duxhell, kcow, ALargeClam, PyroScientist9, enirei, Btgoal342, Ademir, enframing, Geobug, Onionne, Arjaeniodis, alana39, karlamichelle07, nba, AldonAshling, jaminrol, arminor5, LS1222, ashok, Siclone, Thej Bandi, NebulaM26, CEAHarfield, WajKnight, jujuliter, soundu, dmilsh, amktakha, garmili, Natachi, paigezellmer, gigglingCaduceus, shuenja, idara, suzim00n, Trapfether, dryasir77, Psiiborg, Papushoo, Alex12354, tim-timman, bwgregory, myfacehappens, somefish, shivin, Megan.Shellman, Ghost_792, TyTheSnek, vskvskvsk, Davie1031, Lui_Jime, Enderon, rhiju, gsim047, maximemoop, joshswimmer, BOO BOO KEYS, 3DCat, derederevo, MobileForce, properboredom, The Best, AnnaEThompson, Martiel, tigerreye, Bobaero, madistaub, RicardoLuis0, landastronaut, PR0SPER0, NuclearGopher, VanCleef, nnunoo, R-sham, sha28, Gargoyle987, merlins, a.millage.fox, karia, kjstrange, mbharrell1001, Tsumome, bray.sean, Marlyruelas, Kibeth, Helkin, Me2910, lafeedhiver, asterix25, ciechan93, volume, spetrov, royce, bullmar, shaaaaark, bcmonke, Ssschah, seanthoran, Jamescabangon, maxbu, Sgt_Soju, derajer, fangyu75, Moussa, lilyflan56, TopSLoth, Hannah6424, mstewart1001, rohithk1, Eli_zabeth, SilverSparrow, KubaW, jhpuzzler, ginnyy11, kaur, jgpfann, Thing201, TomorrowHero, YusufBP, si225, dshotz, MrSinusas, Guzz, jiwonkimkorea, DeadlyCreator60, Thiefington, GlebZmeykin, griffin21, bimmerville, baye.parrish, guigeek, GHMGOULART, hepster, FF!Sans/EF!Sans/UF!Sans, Bibibop84, aab02, thursdaytroy, stinkysonshine, AsuMagic, fippo913, Scrotie, 1nelsond, Diovion, Ghosttown21, Kevin Hao, bosspackers3, 118cc, Julia_Dickel, DarkxNight, asterisk88888888, turtlerock, Kirin, Scorpiodude64, drigerboy, meatballlady, KasiaSmyczek, nelly, verasue, yesmar, httpOmqCxpcake, hsquared, AlexKomar, TheOfficialJJG, 3rw4n, snapdaddy, Kaidjy, mercyfull_fate, ashwin.desai, Arcas, titip1995, juliannashine, MrEK, Anuket, zrosenth, brb2ty, shafezi, ndiazh, Riggs109, wombaticus, LordEnigma, jqiu1, aja4089, MCFIREBALL, nchowdh3, trondlax, leva7, NatalHydra53, judd.katie, Jaquelina, NicoCapsi, Gemlingur, nate2222, Nicolas Mulfetti, ethan.locke, rusalic, Collapsar77, karay, zartren, shadeymatt, CoralCobra, txukulun, munamohammed, KillashandraRee, the_lall, gypsymack, TheSouldrinker, tedy, superduper, ShahDarius, firstmatezombie, AnsonTC2006, sience9, eloyharris, Zukoawesomeness, sta1amin, nkatagir, 23TangRichard, lisastojanovski, vinbro26, SeaJayRNA, Joy_Fan_Roop01, cedric, a.yates4, david.biles, Sebie, gemini8, bartaner, carolynkim, Antoine ch, ertesh1, avaxu, Jared_Orion, kondratevakate, bmcwinn, Makala Nye, dprywitch, Daarwin, brutus12, arthuryeager, orklah, jone, Zenytram, thelall, JustGarry, sdabic, achouten, Frosfire, carrot, RNAMachine, jettripx, Helmut Coolmeier, vucko, amalis8402, BlackNight1697, Mooticus_Maximus, Dragland17, Rambe, thenning14, bio101, madcow123, ryand, Jfiep, giovanna_gracielle, gallareton, SarityClarity, shaym, Pako, Uniporpoise, book, nancyw, ghosteur, ARN-LLK, OffbeatCuriosity, TrueSpiritGirl, Jehu, SCarey, Aserita, mds_00, todd6485577, merinarasauce, olyviajefferson, nparker87, pablospalletta, chloe_hess, skrimon, FnnyRssl, Fierlantijn, jup1, frkmaster, Jordok, YariValtean, YellowGamer2004, 11killer17, Michdkr, drjevsky, pinkishdoplhin, daniell411, dshariq, 8wmasreas, CaptainZap, coderah, xarpus, ewalk1, TheInsaneGod, MENTALIST, DeniseB, kitkatyum, Heliceras, earld, domforfreedom, arecka, tubaflub, e721420, rynomachine, droricu, Kolob123, Geminem, Adaukt, Sachera, rtroxell, trollhunt42, Rinske, kbourque21, majolka, Meistereder, nerdy creeper, loveRNA, ezio, angassmith, CatFREAK, FLAmbuRNE1, apang1, Uderzo, pamela.norton, jkm412, Melissa.Miller, Khuran, Ecilam, Kimberly Westervelt, ivanjr89, QuiquePA, Buggbyte, coltonq, Hemell, Aradros, Linda.L, Sfeshey, arunduraig, nensi, thebigchief, Abbygh, ZacBarakzoy, sgerson, HybridWarrior777, b00kw0rm, Light140, radiationist, milla, Amber_95, v4r5×8, sandriix, britneyh, Prince_Tat, pastelbleuy, Jwaemsets, HoraCZ, theaudie, superc1212, Lazernoodle, PanikLIji, rere19, janetmaymiller, AndresM, yehos, Bshadow4, huitgh, emiilykim, smcalpine, an0ther, XLVIII, namo, lizhosanna, ablueninja26, Wgirl32, maijacole, diembe, JakeTheGreat, Been, DoctorWho14, Fishkiller79, shinamin0218, Koroku, UrAnIuM, CapsaÔcine, jmalcol1, Cody.Winkler, Reseal22, dhrg, eiryberry, mklowry, OctopusProteins, karlb, A.D.N, sciencelover, GreatBigPhony, 301sebas, ibib689, BZZZZZ-7, awkwardben, danhan, jessie-huang18, filman230, capu57, fixmancol, hmp10, ziplock77, Mechputer, oriramikad, fedor3000, mathers0606, brbrenneman, chemistry, Nosyeye, Billal, yoyoyoyo, Sockbat, test432, rayan, mkewing, tobi175, sheatynan, Asmath, luthe, Irtaza, 2bjackson, Abc123Ultimate, PuffballWarfare, DrBagelBites, hawkyle, Cat Legge, JustScience, blongey, dredstylet, coreywhitten, shutchi3, 10to21, Carlin, jamesgraham141, frolickingpotato, hibobyoyo, airb, tryst1n, Themannn, kollise, ddeeeffff, jdbollin, Amunre, abcd123, Epsilon, Leafheart, jephnar, Lorgelran, HannahHend, molding noodle, Ana Carina, pl, gredick, archie88, 18kmcgrath, tehstripe, amit233, Preofessor Goose, kurt5413, Triangulum, Im just a kid who’s 4, JavierPozzi, traviswavr, Frank_, Adaixa, Jose Da La Garza, dnarel, jennifer.graves, El_tequito93, steven123505, sarecguo, annisa, slc577, MalenaLy03, UrhoP, avalet, Anand7, Valko, kalex, Carlosgrr, datgtaboss, nbernema, JoshuaBui23, SunMaeShine, Piaf, AMD9009, mic.sca, Illyreia, swolfe, WalleFails, Zacharilliac, ReluctantCynic, ghandolee, kortatu, alclsqjtm, reneethecyborg, NatTheNat, Skyesis, nfernetti, Kalarily, 1617674, BrianHo, mrhottoe, Meepers, edenb22, bc32065, rogral, KupaTrupa, pnd3m1k, pereiraj92, tjv, Pinacho, 118ge, 0something0, Othernaut, IceJumper, LizzityRose, houbar, beacandy, AlexanderSach, TsavoritePrince, lorraine.smith, adwoskin, 340atlantis, frankydp, ThisIsMyUsername, theanoxirouchaki, Noru, masdar1, shake_a_leg, Stimpneider, agomaa, seny nguy?n, tycase85, IsaacRNATeam, giulianova, golux, david0mckee, babyengstrom23, LoveLife, KimPossible, soreyabeltran, mskibniewska, lollitafermion, zombieful, Nanachronisme, tobiaszkois, amandaashtonbooth, lilioniana, Heathenist, davisaj5, andrewmcamp, Taylormhix, Grimalkin, wolfbela, Atiur, jore, sjs2609, Dr Moriarty, Pr.Xinel, rivana, bigmanskey, JulieJ, Gabicat, Mkay21, Doltboy, julianguyen2211, jk70, CHaosDragranzer, Kaizer, acookt, LubnaKhan, derfred990, mmille30, SirRainy, ZenH, conradlau19, Snr, daviddag, litewave27, ezlikespie, larry.allen, jaroslav, hattiemeigh, red.arthur2c2, mia.ferrell, patistarz, ryliejk, catarinasiopa, mali, SimulatedScholar, Tritonis, Luk3ling, meowman889, ChrisSkillo, benjben59, EfeGulener, Amedon, eli.apple, szulfiqar, rawanA, Riskyshot, mcampbell24, Kaylamv92, Andrewwu, jakabe, itynes, xmen43, Burba22, dekusk, saepharnh20, felipevergara, Artoria2e5, Roflkev, jschober, Leschu, annainoc, thersippos, shefalilathwal, zaklinao, liamth, lostRNA, Amidinbancroft, Quasar1015, savannah tucker, gtrachel, IceRunner, trystin, Wesley.Estep, Dr_Tobias, Brici121, volrk, FatherWeebles, KimiaN, nothanks, adarshap, IncineroarXD3, raugia, kpham, skeletonqueen, rekone, Wags28, mathiah, kathymore, GoatUnicorn, r_alistr, callmescott, Juliana, timbro, alayton430, Adamf1002, Gqflex, Baccaccio, Saens, Togator, a_kiger237, Keyse, ffwdq, Acer-Rhus, malagrond, Demi2722, jouyei, chimeforest, joeycut, bhredsox11, twstdelf, summ1else, Bulraki, nlautner, Hecaton, kkbsamurai, newrecycle, Coolman60542, DrBacon, NeuroPyrox, klsw, maderjm, Gideon.Isaac.Ong, serpentea, deathlikerabbit, Enden31, mikechn, cookiing, Addie.Simpson, BlueDogs, ER2108, matthapkidokarate, Jen36, mattmatts, toshipwreck, corqjffp130, j_prchal, dmshapiro, CheckYourPants, Pete198298, masterchiefek, yudha, Amlfsk10, Alikat8891, Hugh_Mungus85, mufflera10, samanthers, eterna_player, cozzers, egbrna, jay.stengele, Titouan, jnorden, antoutou217, gpgraziosi, suziesnoozy, arugner, hmkuntz, Damiansteele, dstorey2, Khamul2k16, Agoldgamer, nachi, hassaney, azbcd123, Maxi, devilord, brycerichardson1, DeanoxyKeckose, AniilaACMA, Emily75, AnnaWaczynska, s.davidson, henrycat, rex the jack, JaneTheBrain, riurs0331, Blackpopcorn, mattlscx, malekith13666, Tes_Tickle95, MisfitMagpie, evanwroberts, Potentia, chazz, labworker, bluesmoothie, phil_andering, NeodymiumMagnet, 118ma, sidramalik, NullCat, WhoRNA, catalyticwaste, pipis, pikcolle, dennator25, doudoulepoussin, manongirly10, eduvin, Da_Awesome, Ostiole, mario56865, mhh23, Shorty94, msioda, horacepang, MineMaster, yudfghe5y3sg, zazz96, yevtal, jordanfield111, lecaco, Tyger257, ANN, SpeCtra, Pikapie5, Mahboob, SoBreezy, popdopdap, carinatze, LAW1205, kj415j45, Edgar10711, lorddragon, Reilly1, studdles, prplmustad, krodericks1, bluemeeple, Gannis, mikeywally, tolvera, Ludwik, aleizter, tristandu03, sergioyamasaki, Chris_WPG, EvanSMCK, 9810173238, Dreadmaster, bord12354, MisterMikeReed, mathewdee, derek1906, boomer52, hondalove49, jaquelinegondim, paleo2014, geemili, 20267, rnjensen45, Toofy, zhanglulu, C-ZERO, neonnitelight, trystink, BioRocks, M_F, 1RNA@atime, Stebe0412, Pollyanna584, gabadors, Firewall, Madhp123, airplane45w3, Outis9, faielgila, vampirepony, Nathan1, eBAHN, lexivdh5, Snailutdza, blazergrazer, pizzaemoji, Feylin, stardark999, kat_was93, RedSkidy, Space dogs, DarthMenace42, dofusk, matt7188, axehem1, Jakehill, messed17, Nandix, KarenFA, taylorknauth, Frazone127, lkaylie_, NuttyBrannon, jl314, Marky1994, BobJones467, BiancaOCastro, yamatou, aweatherly, SaveTheWorld, rdrpenguin, CadarGabriel, valzalan, Numex106, RienCus, Scythe42, RedSkinn, tiberiom, Wakko, kyumin137, ducker531, duffdevlin, acrogers, Arrow360, toryyamsd, ChiaraBells2, Kayden99, ebolacure1, shimamakani, mateiradu88, sean4046, CCTYx, IvanZhang, chingyinwan123, peanuts45, lkmo2858, aude, aeppacher, nidwhal, NicoPanda17, jimmyrems, 376390, Tracisullivan, JonahAbrams, oliviaugarov, Hubbard.Jacob, tiger856, alowry555, Caiello, dungpham12002, Akozu, d3xbot, sanfoala, ARFR, randyvasquez0, jschmiege55, balavikramkt, Daycody1, iluvlife12, herrmaddog, jerryfizzsoda, Ivette, mecca417, maddiemoore2001, Grimmjaww, basepairworker1, christineortiz, Joel1, alroc7427, ukmjhngbfvds, Motag8tor, marysims, Sara-LittleFRU, kowman, ButtWeiners, squish5, jcadwell, dillon.moore, YangToinette, Kaljiz, mindfull, narcomancer, 940410575, HHHarvey97, Okawashi, Detovan, goku, aresathena, Bobonius, Tuskarr, issailius, bard10, Actually, blooderised25592, levimcc, Adrijaned, Martiniella, rochelletr, iuzziel, eselmon, Mr.Phreak, enshem, bremlin, Neutrino3141, Gingy, fsysy, Cyber500, Adebisi, braidweic, Gritz72, pinkdolphin12, OleNeuman, chase_rippedbody, eboiro, olerock, OllieB77, sathwika, sherryhe, jshanley, Kenneth1024, taylor_welniak, robertocf, Felix93300, LMJMeman, rasyidufa, backster, Lazy6787, webslinger, nlivings, criptus, CoreGamez, mhornbuckle13, sapiens, emmahorgan, rebsbert, the45thdegree, beaumontbass, amelie.tsh, bhuhn, YouAreEmpty, nyulascsongor, asteroidboy, tuadib, trams, maxwildcat, cebeci88, Alex Lee, beaver142, dylanchyde, DanielCS, skatra, kojawak, TechGeek16, Snaz, laurasligar, absopoodle, roscoeae, Antoniop, Pixalstorm, Stretch45, sandwy, Jlamps, aboersma, ewilli16, jtaylor0089, UN4, chido, jrm98, MCmcswagtastic69, lukgil, Thevoidguard, alisare, Julius.Wulf, sabrina_bard94, Darthlich, jdikec, DragonSlayer99, saree, 6and4, Lord Naanor, hboshra, Ibrahgj, OlaBackman, hjkki, lumb, wandofgamelon, kghd, kingofspain1234, svperez4, ovidiufelixb, errorlog, abkernow, bezago, amaStarr, awaite3, MrFiveDays, emilyh11, pandaplee, justice17, thepope1233, mlaskow, ~closethedoor~, jdwest, hyperblade, jacuelinenguyen, ForestMastr, awokral, Murdy, irishakes926, Mikeypants, mielcars7, a.wald13, Liszt2310, deathwalker1, raichichi, Clarissacordova, mgranski, HatimRF, ecausey, Yan Von Triskel, Unersame, david112358, leverly, TheCoreyGreen, Mitza_003, kinga.urbanek, nmrkaist, darkmoon, jaga, Mamo87654321, robbie, cc1122, Marius37, jherrrick, timothy r., elsandosgrande, luis_93, rose7326, romsofi, Queen_MyMy, thewrz, BatJJ, zacxo, dand, sefaroth3, el18839, DocGratis, laurelca, Nyabi, anthony4tner, Cataclysmicseraph, rachelbug24, lemc, detective6, A scientist JR, corbak53, gagarcia20, abbye.gernt, ifni, toast62515, pluto, renuchepuru, Fistandantilus, SirDylan, mwlee85, Daleron, mrnolanmorris, C_Aarup, olive12000, huntereddington, LuvesScience, MadScienceDude, haase103, homr, Slenderman60, carnaverone, sfellman, Salad, Deepayan_Sanyal, gauree.srini, azerty, iijesse, kiavic, Pengilla13, Alx, gabriel.thomas, Dilldog, Bhavesh123, Noel, joanneklaas, Latrok, shelby.ward, edoved, Solipsis, galen444, dumana07116, billettkp, jeemusik, zork2006, Zacheryery, Jakarrie, wildfried, mathwhiz5, Ashfulness, Hristo, Zelyn, markthaines, jerseycitybio, noelnova2, caleb.coine, perezed10, srtokes, Dumpling_Trash, cutiebearjames, Ischenk1, cactus-lord666, gmontanary1, theinric, GermoSG, theguywhoeatspie, bonniejiang, akwaugo, foldboy, NeuroLiza, Guioma, cyna, brandonchou1, ssss4, Laminariy, Brook, roman.kishchenko, AmandaCookies, pazzze, SweetReverie, DyingGiraffe, jgp, emoell, PerMortensen, elcapitan, Grandfather, VanessaCruise3, dmayer1, Austinquarles, TrueCuriosity, Burnaboy, lindabennett, Josh.Upchurch, MadSc13ntist, dareng, truman, haager20, emi025, WindRider004, gummi059, beck_ike_, Azen, anurashrestha, arthur2500, e666666, Raitmeri, salmonsalamander, elijah, traforch, peepeevs, Person5000, rgutierr, Zillamon, fotamas, coup1393, aundal, StasGtas, nightcore, Manesero, 9ar7e4, Ermagettin, apolloheo, NoahKlee, superkiller, alexonaci, hTRpardy2015, kamalparvez, iamgoat, ahonan, michelinseal, VALLPA0420, rahul.anand.1, Nemesis18, lampfishlish, Ill_Neglect, Frankentree, eva12, JustinPheonix, isaac_verdow, jumpshift, Tonner, Lalettayin, alicemiller, ZainKeegan2003, syn hd, ProfM, RVSdoctor, Pirocat1, Artarm, Blackish314, Spielberg, Biologiste Sceptique, wither, ethanoquenda, Nathaniel., The Floppy, Polika, pGialis, nysossatrina, kindekat, mhealy, MLQ3367, Matt118, Dr22, Aroswenn, yoink15, ShannonNN, catarecat, mattan365, Sh188y, el_guapo, Jeb, Hugh Mungus, Atani, adeline, JacksonBIlbrey, VectorB, immaculatekraken, biologystudent, SiddharthKrishnan, Wikos1000, Ryzal18, glamur8, brichey07, dmaar14, Bladeskiosity, alexko, Wiclara, epf13, Gonzostono, adunafresca, Sattos, JoshPhillps, castromirandacr, maziar-67, red1commander, Joeyiants42, Jesse Bourret-Gheysen, youni, jujul0006, Julia Le, mf.whitby, cgmoren, superib22, DancerTheo, AstroBwuk, bojanje, DarkSight, ngamer, mohnishgs, phoenix2050, natalia, rtwb, FA5878, DarkMKH, Skynet0, Devecseri, jenniferalvarado399, yoyo3654, Rorschach, adriola, IzzyIzzy27, mgish2, ritutrivedi, AlinaD, nacho90, jordanjernigan, h.kruchoski, gmiller73, MoMaDu, RanbowDerp, tho585, terrariafan, bwolicki, jimbob32, ayylmaoo, Mokaregister, Spudtagus, niseminoshiro, shawnguy, haseeb1399, DaveTheMilkman, Volonden200, YankeesO1, manateegrace, wizardmuffins, meaganhill, porldolphin, Charles Olson, Nightscloud2, tkc, Snowstorm100, jackb33, loulizbutt, somiew, bergen_paige, rebeccaloha, deathwalker, jdgibbons1, rajasg, claw107, ram-ok, Taylor, connard, Ikrobin, sir.lior, anishyboy, Reaven, eric278, Bastien.069, dannyj, MistahJ, Cherribo, rg519, varsha_menon_01, datboi2805, MadCowRNA, Ciren, nikidavani, ncpdanalyst, d382, CrematedCube, LOICVAL, joyousspider848, rgne, daniel the futurist, P1a22345, Than2000, scienceking, Wilglide, Anukriti Dey, Cajumasi, BarkingFox, puzzlefreak, Aikita, Schnapscience, paandragon, ekbergeron, BT, Attila Arslaner, WolfyTheFighter, Peppersniffer, BFlash567, CezarRullz, biddikrik, Nino, Qriz, Parmenio, mpw68, bsw42, wrose13, Jason_Kim, kwee23, sleepdeficit, Aluxx, thetume, Rustin, renatomoor, adenk, poojakumar, GRune75, sierra26, sleepiechika, Calsapal, ANDYCHO, phicell, Joak59, cubrusso, sofialombardini, n8tron307, MxylynnMxrzo04, Queensalis, LonesomeBlue, Megaoptuimus, shuktija.sanath, shubham_patil, rohithp, BryanG, purnhagen, tanyatejani, wilgustavo, bulkhead2003, birds_and_birdies, Jaydrop27, Fluffymurderkillerstabber, myraelon, Kumatamo, eragon514, slozzu, rpavelka14, dinoserious, Dna557, adrewred1022, chocho, Zolthar, jpenczek, franz, Prysm, Harry Merrifield, Camhi, DJM_ch, thiagonchuba, Cubbydo101, jsicksate, vigilante, adrienne_richards, cbach2, yehielc, dagiles, fishchick7, keysmaster, jessicaangarita, WiseWords, sifu_g, Rodrixus, dklewi, accouto, Sylfare, Abby20122, lgroothoff, 50, walter1996, Sungolem, mzwee, EternallyGrateful, wacko star, jasongrey, Kingsley98, -Flexter_Dexter-, Duggan92, HarrisonWells, Quido, ltidwell14, Naughtoncolin, kirasiowle, jarrett.talbott, CheesyChaplin, angela.he.1, hoffmac3, 22itay, nickywojtania, reindeer, laduchme, CarlOl, pureid, SilentET, SassyCassie, elisewinterr, AJFILMS, yeshwanth, Narrexx, sissilessard, laughingjack13, MoonSlippers, 587003, ayana.shibata, HDCEOS, Kyosean, mashynv, gdeioannes, mallary, lebowski, ks, nmsarm, itzpapalotl, mdraper, skagel, robertalden, Jackal, duchesnesuarez, DaisyFat, slugiano, ScienceDJ, flom, tygerlin, Jordanacrlee, Brawtik, randomath, IAMTHEJUAN, lexivdh, aepollack, Toooooooooot, khendil, dyosifova, Luxian, rebeccadeangelis, deryny, Pixlow, sudachi, lenamolda5, JustaGamer, wohlleby, ttran19, Alpha10000, Lukedellthecreator, gualys, g00lywool, drkyuremphd, NotoreoN, margienide, Mizzrim, Delta_T, jackwall, jake_breaker, sahkto, DutchMama, biggiesmalls, Invisiblord, Scott40, Kotchar, zmox, deezedAf, MahmoudAgha, donotpress1, wadebrinton, karener, GSicknessTV, Patman2305, Pondguy, Monster Truck, Careinho, odile890, joeslick, Zanejg, abilamb, bigbiff, sylvainPicard, kkenmots02, PikCeLL, Meghanjohnson88077, crculpep, Darleenax, nightwolf7788, Xylon123, jcourain, swapan, sarahpo, dixsonco, Robotminc, hlmeyers, goe01, MIxtar, Bezu, clayshuffel21, musiclovers, j.hafer, fluffykitty1022, Kara.Johnson.1, Bavai, Maefic, Juppuh, marygonk, naomihogan, Dorvilus_B, samroyle, Lomqe, Genius1999, Sraemer2, davidgo24, tayaress, gracepaik14, pojoman, Syluan, mclandt129, Toniche, hamster, eternaviks, modayil, dicknips, cjordy98, tollim, sunjay1123, dielsalder, Reki, Ultrami, Kendall Horn, sprout, gyrados, Alexander1022, artholax, m.sayyam, kalexander, zhet, ashleychanpong, edetpog, sasmtergurl, Cybernetic_Overlord, Hoopadrien, thejet, filwithjoy1, Jhuul, DragonairOmega, AkusherA, Supercoolman, samaracarterforever, AlabamaTF5000, pierreyb, futurist, remysanlaville, pmpham, LiverIsCooliGuess, jnikkel, hhild14, leeeezer, modziom, jezza1764, dddevindev, dosgig, VioletPines, gmasrian, Kim144, 1lyke1africa, TheLivelyGuy, Moleculejigsaw, cloudeeo, TheMannyMam, kittensrule58, mclem, auredstone, diamond_dust, Ezekiel Suh, mrityunjoy, ScotsHack, Eight, gilaadnir, AGioielliere, noemjam, Jaydon, achapet, JeffreyWu22, JeffDeaf, alexishill, kendall, rhyatt, redwood_11, sarahzar, elrbrown, peanutbutter, cheekyben11, piesandwings, derrickthewhite, TheBarkingGoat, batesh, dfour, lordemryys, SuperSine, IsaacKnight_69, ProjectWither, GamingWithLaura, 1venteiz, bobsledai, zad114, kal5791, mjortberg521, amitsahoo, KTQWestie, sueunc, TSW, gianathyaga, travis888, amyopara99, fogminster, Nynyo Sand, SolarNight, hana_hirano, Normal36, InfinityRNA, darthkyeong, altostratus, harrytheninja, RAMETARNA, sokhovat, IANIS333, brewer.noah, samannihal, mael2014, turbofunkroosevelt, Y.PLAY, SigFig86, morgan.higbie, saintwalker, Nathstar1, Smartest Boy Alive, noblecarbon, hemophilia, iopi, exactly85, jowint, asher, pratik, iamthebiologist, nataszachef, Tmcm, Dusty1444, la petite coccinelle, howierd, Mrclownman, Narastor, kdembowski, spopovich, Starcatcher, BobTheBlob152, spiroszaf, h1n1h2o, taha2002, jeune faucon, kaleighspitz, 15minutes, ian.way4, Deuto, Jessicaransom, csgerlach, CDmc9811, fanhuanji, kotenok2000, retrat, Lady, laurenblaga, Annalie, d^vsnubbe, Robotbrain01, sebols, emilysferle, bruno1618, AqueousPanda, Drewh19, dahyun.jeong1, michelleaye1, Jonsey21, Djurz, lay, DwarvenSteel, Trivia_Lover, mattghia, REMINITIONS, Jennabear98, Goats4Life_, Tha_Croat, D.Levos, metalmcfall, felics23, Gabby, petrosxp, deku_link, glace8, whenyouhaveadogbutyoureacatperson, cwong0319, Nikolay1, Edwin1, Noyboy, CaseyC20, rachelerinwilson, Ahmed Ramzy, XXDNabled, brudad, samtliga, guffy1984, TimBell, ENRIQUE, leaozinho23, Longmire, DLobanov, duchowneyg, toto 85, LAX4DAFINEST, HRutkowski, skoob, Doolap, mountinlodge, elicita89, adembo2000, 1scholzc, njia, aw, Hobbes1989, Itz Masterii, madmax24, brofessorSC, emakdaddy, mareDeath206, dchavez14, Happypeaceofpi, sfarris2, Squidoss, FunSize101, KrashHazard, Layla, sudipta0413, Bzerotin, MikelParsons9, albinohunter, marialominos, JJRodny, Jacob Hoy, pilF, JFLO, tombot33, jstanley_7, boky, GumiGumi, scb0044, aarntz, lord_haxon, Naldrun, Llath, jhhu, kristabarnard, crackheadbobby, dougdibou, Agrile, mister_fancy, Corlin Fardal, snickerdoodle, riedigost, kyrian, royalbtc4, evanchooly, Eos, Tatiana, clovercow, A6ronD, OsKy, ashokkrishnamurthi, RDLP, HD_thisguy, paulina126, Saniya Gayake, kinkajou, Deb.Tong, annettetsong, Sasirekha, lgfulton, Yasoo, peaches29, Ivan Tiniakov, Honeypool, ScientistMan96, SUPAH SANIK, Eli H, MeatShield, Asturvidor, naf2002, joey_h, tomasek, winterwit, semoir, Tigran44, vegas2003DG, maxfreeman, purringlion, r.miller23, Jokar1331, atomicsneeze, Ihmed, CelynBrum, zarzo2, rsealcoon, Anima, terraform, Pangolin_Ninja, gpj921, hawinst, silvermist, Bearat, Romancito, ilizzie94, pacoduran, DrainBramage, DanielHulbert, phillip.renton, narki00, pkp98, kmwon, tman555888, sma97, Aforgomon, jst04004, Jackson_winger, asshwin, camaxandra, iamnopyro, Xambonie, MaxedC, 6farer, monkeymolotovs, snidernatalie10, Inglip, DLoh, epunk_Andoni, sciencingsara, awesome_tau, Koerli, tiffany.kang.1, Rickyslayer9, IanAbsentia, MichaelS96, derousib, amavan, Zaelis, autumnstar, dawsonb77, nop, MagnusBrickson, Matthewfisher126, UTookMyUsername, Max2003, JettPanopio, thetrol, teahus, scd418, ameliemelo34, ro.pras77, BDNC, draster34, sunrisedewdrops, nadomako, stephane34400, franclau, bliblu, Tyromancer, joleen teo, Nicholas1024, Pumpingfe, konradm, cbargell13, benjamyn.jefcoat, tweedy, Roderigo, km94, Methyltheobromin, Major_asia, Etertna., gonza460, jorosco, hapkidista, MrMcCracken, morganafsm, micah546, ALOT of Caffeine, molly.minnick, _brie, kv53, karolina.pietrzak2, andrew.duvall, prettyempic, Sylens, aoveony41, ananymous, iSteeve, nata, imoonbeauty, dafrog, WeirdAtLast, Sir X, baconportal22, Solomon19, JaSun, xna_north, fabulacraft, theiris, Shardy5421, sgrocker17, wydaniel, HAIL x 2 x Pitt, ca627, champgnesuprnva, DylanGames76, Dafitko, Cell(ebrity) Apprentice, ammydaroach, carolinerrojas, Aiezz, alyssaabram1, EPICCHEESE, Sweeteevee, CyberDemon, Fer G H, sullivankeegan, mbi, PrincessTaty, eoinmcq, Rin826, Kingjones, enavaljevac, Covenant12, BaptisteG, Sebelgique, camilaburne, Super Scientist, casmineg, Drrozz, HAPPY CRAPPY, viktron, hkim1223, Nthy, widder, Xenon345, Jayfeger, bravehp, Nachojim, Mesp, etern4, somechemstudent, ulfik, thefatman, luisramos, nightisday, pashtonw, Bookwriter, Hordvar, kylewalker, NYX15, maschakk, OptimusPrimerib, sokimen, Alana, xav178, LewisB, ROAR, Kramonth, Sung, cmccoll10’, kamoke, jakesosne, tagardner2, soccerkey, ToloveFLCL, steveycline, Irminsul, a.luma, meli.drd, superkuh, dank_memes21, Manroop.B, Elly-mao, may, pupseal, Wekzel, estradadrian, callmeag, Pysrilexot, lawmer, someonecool, 2016@eterna, Karyorrhexis, micha, cynthesizer, RNA4CancerCure, 303974, jjensen, Inevitablylogical, vcortez, gabriel15, CyborgPenguin, geekman9097, uma.kelavkar, aochengco, rhoffee1, Mysteryman03, Destmon, MacawSister, vanphilc, avia122, Deaddarcy, SleepingInsomniac, Jose94, joshuachou16, RNArer, ptlappe, mkebede5, kfeatherston1, kgodwin3, wteal3, Lsni3775, nast, Lou-Lou, nekomimi, jenfairchild95, epicoy, Aber45, ztokuno, RenA, tanveer, berna22, prociewbor, Cmscrew, PeterPan, LAB man, dfilias, delmo, wither122, i8ucl, renots1982, MOISTENEDNOODLES420, Cantigaster, spamboth, shollrb, seif, gabrielhawkpot, AleksandarM86, newyou, Aillinn, epinto, joepet, LudusPater, millb, DeadkidWalkng, veshka, Mr.Jokr, EverLark, ajpartelow, zois, Electra27, BaconKnight, hale6, ringo45, dianafedjakova, cacorderojr, AllanL, GenuineTexan, monipao, kedarchie, babitaneupane, mkap, Sanora_Ates, jman15393, Jade Keras, En_Dotter, RNAstudent, thelivelyguy7, ggcc, yasky789, scaff dem, eure1088, SpicyMangoes, Roderick H Yin, kasakito, alexno18, judy17, VintagePotato, 66jb, kateadams, LilliaB, skytheasianer, giis, Isabella Rose, TheBeege, YanWoo, gkramer2, CaptainPankek, jennif3r693r, pojko, karenlin123, Lgrace3200, gabirelis, Nashova, miuosh, harry.w1, mporter14, dnguyen14, CatmanFighter, nuclous, asterisqi, diogocurralo, kennyiscool, Amargi, Aj579, kitten, anderson1116, slovenka2000, crooksandtheives, IziFox, Kyurem81, rogrido, Ase10, Airlegoland, david20090523, LRaske503, gaetan, 186879, Joah_135, Vahme, sks, yassine, kennalynn04, Doremiflustist, eternaGamer, bwabwa222, Fallontine, jv0535, violentine, agentpixi, the_sum, Jettaleigh, magikalex, NinjaScorpion1, PedroRibeiro, justanotherboy, hydroxyl, NiCramer, yagod, GCTB, Magna, Sushi Cat, janeisakaye, Sapin104, asilentwolf, rconniving, Tseng Alvin, Vad99, hashim, Qpid, mace, jww243, riveren, fearsomerooster, nbwtf3, tejaspchat, Theonintendo, nicknemer, achapoval, lpusic, lahein1970, Paulgreenway, pmbrooks1, Raikore, ojmao, addt, lewistodd1881, isaac.z, qqq, morganhundley, citizuser, selosari, willswish22, TheAlex31, htaherkhani, emm.grace, nennie, EGSD, grays107, Nick Sousa, ShadowCube, StepchildoftheSun, joseph_seamons, Dantara, Wolfram74, qCricket, hcurrey14, crazyadz, nicolasfdg, Ernestodlc, N.poox, Blinkerz, colinjmcn, reidrodenko, neefer, Elfy, will.colbert, nickbudz, wyshugart, theresa16, mkeller7354, smarthuman, rmaloy, hashemsalem, jackdog45, svanderheym, Jaixmemly, Renton_Innes, apsk7, MorganClark, patrykwielowski, callywally, arangosergio, sequoyac, G.W., snidersniper166, Agk2014, Phoenix26, jed122083, Semored, maxispazio_1023, AJT333, baileyhirota, elias.erwan, Takos, Emelia_xo, nancykz, Ayoin19, DelToboso, erythro, carlguack, fjs507, elsherbini, Astuch, Zageron, YelyahNaloj, amberdanielle, feli0922, j.foster, popsicle2019, atzi8360, rpmolitor, guido99, mcdermer, shooresh, HoodTinkle, ansonberns, 23chrjam, ankitrana85, texankaren, achilles1205, floyd, octopus1414, moip123, Tatane142, novemj, memesanddreams, jvoli, annemcnairn, test535, Xodom, kirankmr, myceliums, AgustinBrusco, dnwilson1, kcarmen, packa, karim__2004, BobtheeBlob07, DoctorWho16, Xavier Wilson Lit, momoclark, RNAtestaccountGGG, cardsox425, Kcrawford, marisvalentini, cashcon57, sigmundrs, sencha71, Ftkalec, Tonyshia, Seha, TheItalian, shifterks, grillo, MASTERDERP, Donbo, AurorNayt, DragonL0rd132, ima_cure_cancer, jlee7, IronEve, jacob.libbey, Febe, Mathfrost, Claudia Vivoni, kristynakulbova, bdaniel1218, waterbear, sthornton3, Lari, TaTsE, saffytaffy, OliverQY, shudson10, test515, justksketchie, Jomuel, klapm, Jackson24681067, Fox_Arthur, skodborg, TD23ASUS, tudunk, Goon124, laniejung, idelmuro, mannylovesmusic, patrickseed, Nic622, mouseofmadness, potterland, jacobk24, j97henry, alina.fabyanchuk, johnskim009, 78904565, Elarim, elcaptain, Silesor, Gp3115, tayloary000, Astrael, maceyway, MarcusIsLaughing, jordanbonilla, Azor, gravi0la, DaBurgah, marteden, adkuiper, ricarn12, mortrek, matburton, aadevries, Kappa_!, El_Gordo, Rcook14, ntegrtytildeth, MicaelaHurd, linneacookie, Sevan, greendoor, 1grafc, cdsproles, WFLO, malachai, Percievel, Fishy Human, grogan, atla, jennifer.popescu, maxic, warbear, darkmoonrising, Dr.Tentacule, adilliot, joibouphaphanh, braxton779, kcin2001, dracke1769, Litixsam, Hydrollium, Bobutter, richardvahrman, davidmj101, SkyWolfy, aturc, loverde89, trutruc, Vonkensington, g4bb, photonjack, biosos, gavingavinchan, pemabro, EmmaCy, jennythefennecfox98, darkcoolboo, kenneli, anbailey, mindofmystery, elnirath, ishhhh, marcohern, Varoth, Jenna2017, jorge23, weskwong2, Allymoose, LilBluestem, serhat, BobGamer, vimes1984, dacp, jelliott14, Jailine, PMEANS, iqir, Lejibet, schaunard, CMOsimon, oraniki, starbort, iminto, WynterMoon, rpetter, benckx, casimirth, JohnMcClay, sgarrick, bouriquet, cabezablanca, ratster99, pielikey, Connor R., bubba ganush, Neuron13, rng159, jhyoo, Codexys, dianehardy, hotcreek, Alexis19962, Mr Fusarium, Melanchthon, BHooper2016, ek7opa, Bsnliu, oml13, Nicolas.wang.1, emoglia, [Haku], cobel, bibi, Aixtzr, catpoos, Tooth, h2ik, mrnat, roxanneheart, robolobo, raptorrex666, AllyV, JAILINEEST, aschlueter, rapefu, Triste Realidad, 118af, TomDiderot, klodrik1, fusiongrl1115, HCBIO404LHoffman, MagnetoK234/rna, narayangill, Lumpa9871, chris2b18, D0rsug, acloyd14, cgandon, chemnerd, jkehoe1, aresetar, alsup, etseco, kforce214, julywaters, alecrdj26, rachelt27, Park, Mehdi4719, CC314, Zaneta, magelan913, ilerm, justacea1, Kittyhen, znep, Kuk, 1RNA1, cosmoletchik, NBG, BobTheGreat, JoeMancini, Neal the savage, Rayzapper, Fennex, jlhanes, Mattymatts1, ZRussell1, KthulhuFtaghn, DrStrange, openhairpinleft, 1245678910101010, test1306, fritangas, Lillies240, Bigginheimer, gregrhunter, BENTEN, GerardoJara, mfrenz, anitarobinson44, bobina44, byolog, amparo, mclem1, Shivani K, nogoalz8, warlamb, MasudaSharifi, alexxiaoxiangyu2005, cussoandre, Kiraraud, Hassizzlee, lolamarie, jaweghor, KarishmaReddy, SleepyRat, kirederf19, thegrottman, THEKNOWNMODDER, daWinshu, marknever, Felicio, cocameli, Evgeniia, MANelson, JBthesciguy, esmurria.a, ayowhatsgood, bOip, Ms.Mustard, Vinceble, RedInk, Lilie, 1gr8penguin, rdew, emilynguyen, isaac.sides, wdseth, Awesomefrost, Romank17, faisal254, lokivari, manishsamson, Buddy5534, mitchell232, 17victoceb, at0m, Margotofeles, bluhma, Davidcwalls, hizon8, dhartman, VicHimself, joeywoneill, dufdufduf123, bgyinr, Rhzoid, lepoulet, Salveur, lni123, alexannan, NestLife, sci_girl, Albert Darwin, LouisBarrett, SuperEscalier, Jordanjohnson, Haantzy, Le crÈateur, slack, agbunag, megazarbiman91, LeORtH, dmrmay, Sinichi, clarkrm1, shester98, sana.quraishi, Romantic, alimaverde, weateaboyatsea, SC Cyrus, Preston.smith75, AndrewYu, kathys, SanthiyaM, brittanyrose920, Novamonk, ahzdii, GeekyOtaku, dallendanger, west222222, buboh, clr, Zegnar, Adua, DRPHYSX, madgehugo, Dado, anna.mikolajczyk, BearonvonDeathskull, reado, SaharahSarah, EYEYEYEYEYEY, Joas, steven85, aie, JJAndrews, marius2417, DzikiZiemniak, LigthningStorm, precociousPathos, eren41, dardalte, zako, loop.golez, jry2001, Gerardo12, Los Macheteros, Kahless, Adiictoo, WhIteVIper83478, amikelov, Bubbles667, tonnio, 2017rewphilip, meganhess3, tiou622, chromebook11, J*, Nicholas Mephis, mar109, olivianeyer450, navw26, The Lettered Olive, Chroev, dasgenre, smithj, Endymion, Jhue, reneochoa99, EZAardvark, AugustS, pvincentruz, Professor_Cortex, Veronica Soler, Openfire, RichardMorris, jero1717, oskarvonephesos, hentrekin, Dipakarki, jennielynshapiro, polishben10, Tenebris, chloebeard, yorky, parisamahmud, Guilthunder, aashna.g18, passeurdemonde, sudobutt, Connor0323, jcseward3, MySecondNameIsMolecule, Drashti, 01014750, xXLaPetiteMignonetteXx, samamirfar, qteapie7, jaden2000, GrandMist, jdigeron1, e101, akz191, crystasugamura, angel.yi.1, ameena, Jbuff, Kyushi, lillianatlee, dragonjake, Zelyn Lee, JonathanSchroeder, bwil, Bryco, sterne, SigmaBetaApollo, huthut, MajoraFedora, ana135, luc1s, manitu2000, Carmorik, LesterTheDolf, TheCatsMeow, czikajarl, MissRead, KeiranN95, mouchix, Snowywaters00, Adrianna, lewie, norvegrland, Dawngirl9, Hawk680, nvrogers, Metroplex, pirete5, Meghaford, drjobert, KrzysztofNowak, KileyB120, demonicsteel, Paytonhilling, JJandDjango, Vidar, shademan11, joss33, BearCat, phillip_phil, kenda106, madrover, primus09243, Aminoquiz, nebula_point, ragnaric, abustos85, revarcline, Isakswe, MolybdenicCrystalform, msommer, Dugg, cjack22, mmarcus14, gcwingate, wong338, The_Geneius, lilja, SeattleCRW, abievanss, Jorelly, YusufNM, morgy, Hoanguyen4202, goolywool, episcopunk, setsail, Bnanishu, Mav.16, amber7malik, anarok, DocBrains, tiawhitlow, Edwadosan, DressageGunner, BoredBro, monster, katarzynawasiak, makises, supermarioglitchy56, SamboBro, gmalfatti, MoltenRock, trookey5, Reece Jones, Merifri, gaja, j.cole, alligrace1234, Maustschool, bbc001, budga292, DiegoM, The0therBEN, marcosbraganca, jkalwa2, GoldenM, tombarbsand, gatherer11, superloutredelespace, eyhjiulei, The Z, Thezi, interalios, klc8272, tissana, GabbyHoffler, mleighton, hellohumans, Delvis343, CharlieW, ThEnlghtnd1, MsJensen, shacamin, Josiah94, cwdavis122, ghujka, frede115, shoaib131, dmd, Darth Kyeong, Avecchioni1, the eternal creator, Teykana, Alexander.Tashkeev, antony.corwin, Secboy, netik, juju50, RIdgeC, TheDarkMark, lorettaf, filnias, awatson5, Mahmoud, xohorses, royrschach, fatima2016, AliAlireza, sinanbaltali, zoemote, feffab, mcginn1234, EIMR, nancy.shaw.1, anoano22, TigerRaichu, Dome, richardmalo, uokwuolisa, DiamondJub, Langolyer, Hortbek, mizuchi, freshkide, TheRealDeal123454321, mincerafte, Satruqui, RondezFox, Dienonychus, test221, Cyba_Forge, AnotherKen, willjones1989, bribri13, Buzz1215, Bharadwaj.M, titidead, Adry_1, kkalwa, alph, MFG_hankie, sonono, milkymanda, mendo200, papageunu, DinoNerd03, bored2death, andy12, han.zeien8, fbraccini, softball_2315, Fyrax6, Nexus227, wrhc, jhuber2, pavan248, Ian9, Straw Hat Luffy, toich, jadrien1226, TheSuperDash, HellShade, kkelly4, rashiel6, emiliano87, doorocket, tspencer3, Cameran, Buster96, XtremeGming, corvus_obsidianus, test2903, stock21259, JumpinJac, ocus, WaterMelonRules, dorinda42, jillianxjohnson, HelloTheWorld, alnoperr, gloot, JColdren, Laurasgwd, reapercat, Kobrus, Littlemisswolfgang, Chxck, janeyd, lsnliu, Mrees8, vichawkins1231, brausch, No1ZeldaFanMan, TWLCFoster, womble dung, grandest, Itom, zombieoverlord, 5166954165, TardigradeNerd, maxharris22, eikcaj, Odyseus2, Xinlan, jmcmullen, Patola, kendradavid, J Woodburn, dasoro, pirate123, Pkitty3, nwiegand, Rico*****, Pinkman, ryker, jahuck, HayLeeDavis, ArnoldS74, Zaelix, leipzigRNA, chrlrys, Tupulointi, Justin338, hiouf, Tyro, afhollmann, Kurson, kainoa14, phanu, Exitron Apte, ugolee, Manix03, cuil, Googlichou, npshapiro, marioistheworstteacherever, Sultan, Mememaster, mallow, entropicmonkey, julian0223, whatnonsense, Braidweick, esued86, bo, shaebutton, Spreden, Xenopsylla, IsDoctor, Povan, Nougat, cpfatigue, judekershaw, faliah, wren6991, Codeman5199, shortcutter, nuwar, lukasthesmart, Hxcking, Frelghra, Purpureo, BLTaber, TheMagzuz, makonomy, LiddyNiki, vytran1629, onyxninja, JoshuaBioinfo, 7896542a, Tomato, AmelieMelo, ellenbrady, Dr_fovea, nmohan, jenkinszdj, diegosuarez, barney.tearspell, osch16, SunSimiao, hare, Innerlines, GreatWhiteBuffalo, Anaxagore, koziel, Jbaixful, ragnabo, CarlosF, deprezke, evharte, radecule, DanielFord, willhong101, mrmola, Stryke, makayla.haggerty, emckeown1, pejowei, Daniel_fasey, nailana, minerva66, Ana_Gabriela M, dboy1015, Haojerr123, juan cena, jeppettoh, KAmrine99, irenepellegrinoct, PreKrebs, norseboar, indigopari, justinhuu, Sanchitou, starrmyrtle, deep-venus, uberkiwi, RNAtwister, stuppid420, Nephelai, ejcarino, Laureatka, Bio-Robot, hijoscontreras, signlover05, Ilphrin, Materistic, monaxue, Ana-Marin, cptredbeard, fjade, kedwards13, kbrendel, Kurkio, pagec, grace.lazaro, GMillie, conorjfhowe, VictoriaCappon, ghery, ehs4n, gardenpea, mgeistler, jigishjpatel, syfygirl, Acrylude, JordiSM, fsbx, IamaGOd, shinymetallic, lastbluerose, Gladway, pluton31, brannonlynn, albertox, jaykapadia72, Infiniti1, mer135, RedSpah, Beat Em Bucs, Ern, GetzMonay, DeathPulse, yukimitsue, j4ckn3sia, chris_topher38, mswhipple, cjsprogis, CraftedGaming, rick1689, csherff, rlin, MAZEZ, muscovymischiefchick, KB171, ecline, happyFaceScientist, Alec.Santos, BlueFawkes, coolmariahamster, mgp0127, Aston101, ShadowMimzy, snaranga, restigou22, Dazarroc, gribskov, aelahi24434, jedikidd08, AdarDasha, Poitier Stringer, Atomos, kperss, justinberryCPEBio, fedeg, kromaine1, burak25, maitreotis, AndreyMozz, ElvenNecromancer, kylekyndal, Artifexian, dureshehwar, mediacodex, dlenart, TitanStorm, totor, Classyklutz, cstb1, Aspeek, EdibleMetal, logonxix, cdpeyton, Bioskeptic, boxchanp, Magikakarp, pieoflords, mikecody2, Kwaters1996, dclone2, Dash, jaguar98, acs1972, alfa2, werehog47, sancm14, Wizzdk1, mcie, Maevell, AustinJamesTheThird, POSIDEN2, VastianZZZ, Dirctus, kimnos, xxDontPanicxx, LenaMoldavan, jamesdemchenko, sadieoflaherty, voldemorder, spencersimko, Urbain, OwlMage, aert, davidjm, tutu2666, Andromeda31, sevoro, playboy78, dpfan42, hhentschel, poober123, WATRDAGR, DVSJonathan, dawezzz, SwimTeigetje, darkh, anthony1, jyaworski, nicole.josie, nmeyers, MadCowDisease213(Jeffrey), sergiiomadd, timbio, strayjtu, Cas13, jokafor1, Jackie123321, basg, defectiveDemiurge, cindyzhang99, tanabornstein, Furlucis, deacey, atrain99, NaiF143, Chenrezik, cmitch819, testtest512, Catyboo101, Ilrae, mayrobe, NoraNeko, andrewking203, KriKriBioChem, adecks, skulls_4ever, leloucho, tatalex, test8114, JackTheSlipper, Rdilag, degracemm, rbierman, EpicGentleman, BenBoyle, jc31, b_howell126, cienc, ngrous28, LittleSully, Hesham Webas, mrtrules32, smashy, mhemphill618, javort, nonm, testbarney1200, flipsyde1, Metalbeast310, Sanjana16, jmelamed, Brycebybee, Jaysonium, pathrna, davidemowry, NLDeMino, leoweigand432, ham_shoes, acon, liloute mauricette, hjbrouwer, IronDoc, matou26, Huldor, dipsy0720, sksbell, FatJustFat, boby, lecalaza1, lcs577, sinan, Dennis Ward, Medic2017, napen123, amira adel, Shiroan, BrittanyMK1, Smirfyman2002, gatortor, Arenwyn, queenlilith, CurlyGrapes, jfbrennen, quarkmarino, Bean_Fish, anthonyseaman, Krischan, alandiaco, billydaniels, vilmanke, adrian.pavel, REAL_POIRIER, Obaid_123, gauriketkar, BarbaraJohns, samanthapete9, Hyperium12, HayleyAxelrod, katelinn, teyhla, LEGEND, efuji, Lucianaville, damien.choron, macas105, aelgammal, gundus, rdthp, iameternal1, mistermurfy, BrittGalante127, Ariana1012, Guy Shaggy, derivan, Asj96, amswid20, emmaokapi2001, NTH0005, undeadcowboy, firesparrow, yellow_blacket, DarkMaster, Windayme, grace_face, krissmennell, lizzyleonas, TsukikoCurrier, azhang21, b13, SAbeannie, deadtired61, shail, lptaylor29, desired_username_here, Videogamer70, jcreevan13, test423, davidn73, udeudeude, janghyeonggyu, Blinzzo, Akmmaher, simrun.ursani, dong951103, sadpony, MRADAMS, SkiaFox, oiseauperchÈ, ShyMagpie, mccle132, virtuosummer, bloxbox, jhabtema, rhartogs_eterna, lwongsensei, migythebomb, R0obiin, kat_w, choffman69, nana, zoeyyoungg, sarge, HaotianQ, Zero Progress, Gainsboroow, PaulBrett, manyu, DrGrunty, mrocland, agent3554, tunrau, Devinm311, boopster18, romeubpaula, Protean-above-all, anchuu2243, Xavienth, asebiani, HammerChild, teodor78, mdietler, arcanepikachu, Karmito, oweny, msampson7,verizon.net, Haartemis, Robert2017, SupermarioMLG, sabersm, Thanatoons, Bilal, pramod_recode, vbenavides, Pa1gemiller, BryanGmmmmm, TheBleep, arshad, ingressbetauser, sovinirs, srdavos, lamsi, etsulliv, Profa CZE, slutzy69, weinsm, 1stevenp, andhrimnir, Nuno Marques, swagadactyl, EHSMiller, TopHattedCat, Amira, DavidNatanael, ferg, Lotesse, firesword, zamfire, loukouma, kdease, PoorAnimal, Eirah, kajellereth, watertank, Belga1, hanmaldo, Zonner2001, Samoyed525, Fantaisie, dickeyjohn12, ikraen, a.ahmad, gogata258, fermion, sunnyswanson, abowen24, Sandor Finn, amneto, AKSCIENCE, Whitefox777, antartic, Ch4osknight, pandora, edwrdteach, Hector, xylophonw, jemarroquin, redcat1, _Maximum_, Gzarl, mr_imaginator, wallerc15, NitinG, minehorse, dreampie, silverknightzz, leegaarda, micia2111, Doran.Woltersdorf, Geredd, avanderley, NanoAluminium14, copeland, TroyIsMetal, Red Gab, marktrofimiuk, SB52, Table5004, TeenageTimeships, Jojobarbar, CarBoysFan, samoapolo, Yoine, PedroCunha, bk1010, villavo1020, eviljohnnybravo, 186095, tubarocks, yf.o, Nikita Brukhanov, young duke cho, SamuelTSO, Zink, Izenibad, Wreck5tep, Ian Day, DL, fulikalter, azlen, anynooone, MineLinkFR, mariam25, michaelmior, izepeda, Hydrogen2Oxide, lucas.p, VanOutsider, mayorale_, nkirkpatrick, henry24, Sovian, poplarbul, Dreamer5959, kharkins16, Atherne, lordotw, marcus42, nathanK, Fire875, averde, dragonwin, dcandl1, smccombs, erudium, Skillere, kaelync, mezzol, lluks, mattkobi789, emau, dgrem11, Fg95, Pengwyn, Hippi Nasehorn, bill nye, drwibble, Antiquus, Gagcha, dfriedland, jfowler13, nlstudies2205, Magic, kurujail, TheGoose, EhTee, Gabrielle, Jathany, kuhlkilla, Flodarius, amarkus, butterz, ByggareBob, TwistedTheCrux, rainacorn11, supodin, Hyperion203, WafflePotato, hammolan000, SRain, mutator101, nmcnulty107, K-Train, Sliderlo, jackie.kephart19941, mahdi-g, Rchitty, beto9202, maiya29, nichti02, GameFyre, laurenm544, elportugesh, mabs, aiden3dd, lios87, mpena1, JuliaMit, mwilliamsfu, sarahansullivan, shu, breanne.ault, cydittmann, chia_cpc, Onesmy, Kulpas, drkenji, tomoedachii, krondog, diergotarman, fishfingure, skandel, geizio, Ayex, Ares_4_Life, mariomaster57, djladancer57, vhrico, nailah_din, AnimalLover022406, Dr.Pizza, HPApples, tcfreema, Pe-T-eR, taehmmartin, shay5222, Isilme, Savino Falco, oioipo87, khatniss, atmorga, Frankp2491, LordFlashmeow, kkohli, meka renault, loxie28, gilaki, Obliothing, Zagreos, nonma, mdf17, alx-001, coily, jimbojimbojimbojimbo, goldentwo1234, Jukagiso, odeiiis, spacegandhi, comatics, profbiot, eternal sreator, exposer28, Noraip, Tsa6, Nihilanth, thatguy010, VenusRain, Snake2417, ShadeTreeScientist, Jideox, DoctoX, AadiGaming, gjg3210, colinvignoul, ahua77, Nickatnitel, emmailinca, ellis183, Cal_Capone, Playaz, Scorpion508, olha_vypovska, ACRichmond, srmatiol, GhostGuy, kachupyn, seablue1234, digidude, marymmm, beubeux 74, Platinum_Hunt, Happycowsmoo, valto, addylaurel, chubbz808, Alex132, skippysk8s, allynharris, jingle, SkyFinder, s97, einek, Macbest, dubois, snelleman, Its_Me_Dude, sstrickland69, efilippa, discombobulate, 20170203, xerrix, UnderTrashLady, Catlinv, Chameleon12, dudehdrjs, WagsTon30, ChippyofAmerica, arielm, Xencaye, I Like Pie, MattKeehl, ByronH, blackice, queenjane, mywither, scottabutler, Tildi1, zoniryan, Miguelftw, alohmann, TRees1999, julesverne, NikosBlu, waly206, phizuol, Khemistry Kat, Antony246, chin.nems, Muzozavr, jasminetofu, morg, allaboutthatbasepair, CRIDRAGONS, hsmith16, katakolm, Becks, MormonJesus, ralphandeli, VanCleed, 41cy0ne, niky1, paulserch05, rgeheb, tolland, ntbm, 422201, Maryam Ahmed, elemenopeeo, soadreptiles1, ShunTheElk, NeuQ, LuisDNA, smflanch, nodir, .., skky999, Julio974, hebjo, shakahapa, nstephe2, sonikiliky01, kleptine, JAWS1275, duygu, moli, Stecj, Faitmaker, SuesakuBlood, jrosy43, Josef D. McClammey, adnsalvator2090, annika.chan, LandonZ, trueanomaly, Pario6, daviddesalvo64, Nogard Noir, Senga, dan18662, Heinri, hypermechanic, amritha.r18, goldentreefrog18, hanxyolo, Briag, Jacob A, SirDusty, kcarter2016, AccGaming17, lee737612, TROLLERDOGE786, jeffledwin, cpezz, Sarahi21215, nikesilvermoon, earnold1, lunarbug, SultanFX, photon666, OrsonAround, zh69, mani, josemar, hyphenbash, jepe, rapidash, Emma_Lindbergh, ganipa, rey3002, oly08kckr, Gustavo, ivypaige, bananas are yellow, Octodon, narezul, jocelynrobles, martijnmunnik, AdraniBelegil, roaneskin, althusser, SquirtlePWN, pOne, usiadia, Memi, CRIAVENGERS, ailenvittor, kennedyball, Christian Tsivouras, adamatallah, MuteForce, Fra Frusciante, tsteiner, crazzyboy1219, ivan390, MageofMind84, kiksmahn, TheZMan2960, Espequair, Douglas, Sunshine0, rmon, JNJones1, M@lice, alexisrafael, yuvalturni, ptismall, pazelle, faithroz, miller6790, TheKnack, carefreepenguin, kthakur, Dr.Shadow, mariellanewell, skyeboy191, chsevy, gamador, maek666, UsurPedro, TobyOrNotToby, monkeyinemigration, parski83, ohassan, sherff, dardenofeden, pinkpanda05, daniellejoelle, sfmeade, Crissy, jericotyler, mnrkrs1996, lamaral, Caleb99.0, bobtehgreat, RainbowDx, test1080, TaoTurtle, laurentc, ginomio, quantum154, bern26, wbt, DATURTLES, ZenDog150, Damiano, zhanghua, TBTD10, BiseDeMetal, aoibhinquinn, phillipsar, Mayacny, Alma, NewMoose, derpitron 1000, Hairesthai, abbycroxs, UnDeadcoyote, jisaacson14, fyc2108, genesis11, paddy_o_furnitr, coolkid69, CyanNightmare, CXRom, lolitafermion, nilloc171, asorren1, Kojak5280, turloughcowman, LegendInShadow, Eigam, Demonio_mx, Eikyu, Portgas.D.Ace., tredegarshea, MysticSilver400, tokintalisman, Antiepic, rick shaftington, daxmell wutt, dakota_c223, hunter.goff, letsdothisthing, katwas93, test1380, Antoninon, paola_jose, facial, reginashkreta, Rafe, AFritz, eddiefranklin, ashrivenai, SGSalem, Huraqan, anura.shrestha1, K_Berg, drkill, ghoston, EverLark13, crilab, EOD, gf-test1, Po, rna_farmer, Quanican_Bleu, paradocks, neverlavender, isaacm, aHumanOfPoirtland, berserg1, usarthur, Jlewis, arcamax, sturgeona, supespacerotter, seren di phytie, Echoech, Kiracutestuff, AndyBloop, ride2946, Dpirie76, the scientist, kittens99, crisis111, ycarmi15, MedicalScientist, yiays, nukers473, Killabeez, GretchenG, jasper1234, iloveme, ghost23239, ris23, mccarthyelizabeth05, MATHBOY12, rapunzel152, frocto, OleDegrum, resmander, Mouradif, nich, Golbezmeteo, Telvarin, marcusblock, Tsolstice, Bragolatch, anica, jrthom1, flopau, itzpanda, SMDCharizard, Luis2016, Icarus_123, etinaude, sun158, test12525, brandt.weary, Aku, lxlarkin, RedMixTheWolf, ricizubi, Sanctification, yokoko, Arlnoff, eladiodr, Usmed, otebios, dch10, moiz16khan, BentCookie, Elijahcarter110, Skalagon, Sisoma, Zhenya232, drstout, makin, kouzmenkov, Dax Mutt, gacarson, EO97, RnaLilly, behanelizabeth22, Sousou, WaWaLuLu, Mechpanther, marckristians, tubez_c, OasisMaximus, jakkara, amyleerobinson, 7ships, sathy13, acurtin, shanedsouza1, TheLonelyEcho, randomguy1234, silvan201, bluffburrito436, bgohan, mjk2221, sanya.rehman.1, miniverse, AvaCassidy, dantoto, AthenaFoxx, spinodalk, AhmedAyman, 0smallwm, MrKagouris, MuFuChu, esyang, kroquet, AshtaraSIlunar, Wounded, Etvader, Gerb, Skriff, DRDNA, Eagle7996, AntNGira, Sans, BlueBlaze, Tolka, emilejeanmart, melias9, demonsorrows, LateefWalker12, armanb21, janandraczko, mattdo66, HGMACKAY, pokemoneterna, Milk1234, ACatNamedFelix, roaringlion2002, Lindsayvolley98, CgW2go, romelsan, steliotesd, fkristina, marwan, timboboss, MoonPaw, Gustavo.M1, Torghal, Ainsley, SolarDiamond21, mehmetesk96, Kaldrick, zsuzsanna.gere, Saml64, monkitten, jorgeisabadsoccerplayer, eclift555, willdavis, XenoZCooler, dpedds, Dommos, dfpe, schwagerjjr, vekysya77, tom d khat, moralm2, kpedro, sonicmiller117, Wojtek_Domin, muse51bt, jhnkruger7, Uziel33, rexny, jcmusterman, bobbyb264, jshallcross, Kindama, Pocky4141, fejao, TSkorupa, genetix, Hampa, johannesjuenemann, jonturner, cemangini, ElephantHeart, alyssa.anderson, tanguyofgood, lamcho00, brisingr2001, Sof2929, hgkamath, Baatr, wrch, Sig, andriykopanytsia, larry25427, 13turtles, nick1243, annie33, EpithanyRae, unknownME, diegulu, FritzsHero, myrrdin, DavidHan, saraurier, Jimmy P, matogoma.meite, peebs, pinkunicorns06, devonwg, rajkaur, gabriellecrivello, mopman23, fetiola, alextheboyzander, IBricchi, f-stein, socky1234111, kubica94, necet, Sthefano, mcass520, ash greninja, ingwa, ewang125, davidmitchell12, tomduffy30, Jayman, Penga, 598905032, DomesticusRex, Torrence999, Aceofkings9, ethangmt, GrimnirFaltz, Jellifizh, thepianist59, Whaty, AshKetchum, arexw1, tarik.azouz, ruetherford, captain, ErwinDurzo, bobbey, princesscarly28, KareemDoesMc, zoz2124, vicioushalfling, delkaim, fatimak, dnewton, prismatiQ, yekbunbihar, SubaruSteve, WilliamPesto, hudzi, TrAsHeR, allenwalker, kptkurczak, atrus7, Iz888, Assaulted_Gnome, dudelpudel, torikh, spencer1keane, dmtdmt, bancoesa, himan.tabani, DrewA, eol212, Zence, Lucane, THOREAU, Doller1, scarr9, BlackScienceMan, Homebrew, jonathanwade, MonsterMunch, Needlepoint, tspencer, mkaufman1219, OG-LOW, codeblack77, nicolebessette19, Brussell11, mercious, AnnieQ, test6356356, ma529, blu_blac, alex1454, samieru, OhAEg11, ulrich0810, Emericx, angelD, FlargleFlangle, mwave, ainsley.ballard, BenjaminN, Paarthurnax, axxiomm, Leoco0007, Feline Dragon, Ricardomd7, Chris63479, joelc11111, Maestro31, clemclem06, Tucker, BigDaddy69, Karategirl80, hsgdfgahfafhahfadhf, Vladik, reefishes, robertlyan, dhruviscool, Lifchitz, jagodzinskajustyna, ProudMuslim, AmandaMoudy, wwright, demenkodavid, matt429, jgraff, jwilli3510, kuroi13, Timboskinz, sharksting, aaronconnors, rednaskela81, Jeff35, Jyotirmay patel, signguytodd, stef of earth, johnsonc, Nick Fleury, stevey, vutrangyan, Davebeevee, jujuzeking, Skillzdavid, tday93, ailene025, CreazyMintaur, ??????? ????????, lukiv, WebsterWolf, ccormack, HitSongs, saatvikkher, cxndxcx, ReillyDeffendall, joel951, Doc Roberts, Kwokodile, takagisun, vanessaballeza, satishbty, lydianb, MuseinMotion, ValeriaRod, antonkulaga, gaijin1997, jluther32, Fabian, summerluv418, ddesalvo18, amomarie, mikka_is_tall, Kenyana, RainyBei, coreyday89, Knugen, nathanh, the fatman, rlgcitizensci, amaccormick, sulfolobus, leyl, octaviandavid, cjacquemart20, garrison, ScarShadow, rebecca.dake, giorgio, spymatt88, allasophy, harkiran, Jennifer Pearl, hm986819, Spartakaktus, violetnebula420, lanny20033, j.science, Moody981, jasperjohnh, Marinelfox0521, emelyyacunaa, Hashem, coz, haugrhol000, MartinSamantha, whiskymisky, silkydoug, PsinkaJones, a.mcelroy, Ivansx1, mockingbird, isaac.howell, ps24jeff, maxmax, Stinkles12, mjsbgold, kzazar, hamid, kevin.heinecke, Arcamorge, Grabrabriel, avoliogab, padidehdanaee, Lukia_raregrove, oldbigal, leehotz, ricknm247, AllieGirl, sambea, kyidyl, Eryn, ode58, memoaguilarjr, yoffset, SamCat_14, Zacsar, andersonvom, tiGer72, bacon117, JuanCena63004, Schulio, chris62, KrystalFrances, jarednat, TrueMK, MacGomeyBear, thumperfuzzy, suikun245, Hagrahell, cesarpagan, IconMaster, AngelBear, emily_sch1, xiantk, ratgr, Dieguito, matanana, otmurray, Tortoise, anibot, Andrezitoh, bluecombats, thekijin, SmoCoco, aweidldl17, mabragor, zohan1922, Neel2000, free2choose80, PalChumFriend, sydneyelizxbeth, RecklessCaution, Pablodb, FraggleZilla, r0bert2, DavisKenney, PotatoPalooza, joecap16, brunamoraisv, dilcannon, nate11, ngiaffoglione, Perrine567, molitor, cock-juggling thunder cunt, CoopGalv, leshroom, JakeGeffon, mnemogui, Nehpyt, TP3413, rokccnfv, matteovinci, psychologist, Shadowace112, reginaenox, jeepwran, WitWolfin, fatjedi19, alexandrujuncu, Svenyon, kdc898, Fantaisia, seb1, eflan07, cianisasmartguy, troyang3, Vadkoski, test22222, Carita, BSOM180, LFP6_Test, dhbgirls, hahn7301, bubbalex, tnurlanov, srspore, bascule, VanessaCruise, mahmoud504, fauxnetiks, Griffin01, Revydestros, kprotoss, luismuniz2016, patricia, upinsmokemon, neilc, Lilac, dpfels, npnelson, Superarm2005, Oktay, 4everRNA, pmitchell, vineetkosaraju, SteveWaters, Roysboy72, jennywren351, SonnyCoolMan, bradleesand, Jeaneaj, ErikViking, Shepard37, sesamestrong, Svenzie817, lazerepilasyon, peetereeter, danielMOFO, Filminfull, Daniela, DanaBeve, 77Tennifry, test720, bartzilla, babitan23, walkgraph, cocconic, kcoleman13, dinna0890, benlavi111, KZON, 5target, whereisbrian, quelia509, klctech, lighting2468, fixolasxd, penguinguy, sammex, addu02, indigopizza, samuel.russell31, mariiaamm, kurtnphoenix, AdditiveBook218, Colmeweb, MeganHansen, MochaLove, mave617, fherchov, karlbass36, cuukie_munster, ome, egale, david12345, dfvbrosh, docdan, Paaf, naseanh, crazyEyes, kilker101, Lidgered, blategameR, sohar, ugorur, PieThrower, tko, Hackathorn, Saiph, little light, Cleversy, ananas123456, mariella, 10102320, sahask, Dragonfire, Bonz, Nautilus567, mporter1, kingbenjamin, jenjen, riagad, cbond6590, MrnaRrnaTrna, Palms, SIVAGNANA, aliacoalwell, TrippleJ45, Majeed, jfoster, booby, dick2014, piotrek2023, seeen, spayne, Arena, xue, Jessi Jeffries, christo1, Skp43tc, rebeccaj, cyrilled, jocynicole, raginbrit, higiwili, KORZINGA, johnnie2038, ewilliampeterson, ac0852, Connor123, gatesfoundation-test, Alif_A, Domestic Chaos, Dash62g, r6preston, _SMG54_, thatginopeche, admicile32, DeviousCrow, rfc0005, machinelves, test7129, tekno4, Zhen_T, jwiswell, mikemass19, ZeroTheUltimate, moenco, PullJosh, itsallaboutmike, sostahl, hung10032002, Zimtelfe, Backpacker, PJG1996, lanzhu, mscowen, andrewdouch, andrevivi, test360, ohanwei, ashleyhamrick, achupacabra, test222, Grayseeroly, Noks19, susageP, ChatBot, dawidsimpsond, misbolsos, gjk27, mrf, Chath, alephn, TGerard, rudolphjosh, tlooms, framegrace, MarioLeao, test220, Rebeccareason1, xshagg, wuyehwen, Draksis45, sanelrocio, banan120, Aiko_Eynos, bagnoll, alyssaking, gatesfoundation, bartozaur, Batou, ultimate4800, Akehebo, louban06, gottliebpet, rune3132, jazming11, John3-16, brofessorLyons, TheLovewaffe, HiddenSquid, SolarLiner, rchakraborty, Zesty Kumquat, Tristacular, kelseyfennewald, JosephSchwejk, soumi12345, adilrillon, Raszobs, pervyshev, heyshahboz, drwho42, Moonhaven, lira, Kevinpmac, diabeo, bnhump, Appollo, bob1234, rubescmar, ElonMusk, 16SpringCLauraVlacovsky, matou008, coljenkins, MASH, PearThreGreat, seanm, cvs.manas, alarosa, cadeboy28, kmcmenimen, Plasma2, borkpaladin, potato22, liliputien, Bwegs, TankyKiller, dhgghkjjhg, sauerl2793, Galaxspheria, Jeroen1602, laydylayne, Panda420, Dory fish123, chels1995, Brejo, LarryCuomo1126, Ageuke, JambonneauDorÈ, buttermilk23, Jarte, cssndrx, 10509i2103, DouglasBusa, Top10, amand13, krbillings, classdung, 8lie, Dwarow, Daddictif, YokiDiabeul, tomtom90, RVS doctor, miller1198, PReynor, book81able, BSimo, TWeston, Hawkey, Hoodielum, danielyim919, karentankist, bulha, poj4639, alysabuckler, derkarlotto, sara11111111111, iyess, klangen, selena.osman, Crookwood, cromaxis, bandlife, Davanint123, PhantomGryphon, Samuelada, AcruxYildun, Blacke3, pokilo, Rajan, live_and_laugh101, Eldar, mmhajla, DimitryM, Hettle, pandak, rflores23, nodnerb93, Osqi, maimas726, charlieg, Mike Reed, jcharles, applepie, Darkleader22, rarulu, StevenMelts, jero1987, tholcomb1, bullfrog, bdavis98, pinball wizard, adc04, Hoike808, loupi, Miketung, tastyteacup, henzo, aurumjd, NGzC, Matyas, kamdongibson, Seyedali, Kaz, ls3cam, mei ling, milky flynn, ashrestha4, tigerperr, Pixword, tdohert2, smitty3544, Park Soo-young, Eric Hamber, robi50, corrina1, ReapersEdge, stbo, Psycho, ribonucleic61, K9gaming, cconciatori, Brejen, SecretAgentNemo, bluebird1398, davidmcfreeman, Bubs, TrueSky, rykean33, braintwyster, zanzabar, Paivaan, Erikito, jschnabel735, goacego, Schamallo, coolkid12f, massimo, kylesun, Thursday, Suvidu, kp12345, Frucelle, sacausey, davidpat, chr2, testrookrd, kenzie_menzie, Strum, WesB, suzeterna, Quwertyn, tfinelli, ZAZOU207, ColMunchy, vluffy, dodo71, bubbamcbubs, procterronald, Amelie3, spikeydudey, mustardMan, trey8050, theangryunicorn, srdjan, joetalo, Abhilash, polark, rubarde, TheDawnDragon, noneedforaname, pandasecond, teejthenerd5, Maxis2000, tosina, 320123, sluke123, flixipix, Sango, mz7p40, cbao, edusoto, jorlando989, cheo, Ericlund19, dragonrenard, lydialydia, monkeygirl, stormAlchemist, dim90, Loogia, GabbyCeja, praveenjr, anbien, JamesKirk, walkedgraph122, NinjaNarwhal, eli.weaver04, automon, Meinew, solorofog, takpa, quasar28, RicardoR69odrigues, Cmcdo44, fcpway, ProgramALaAlex, Jannine, DankMcMeme, avzanzag, Koritsa, mbgilles, AlexanderdL, FrozenTux, gfb2003, Didier35, Johnson, than, reissfc, Leakybrains, TastyAngel, sophalex13, Donky, jelly123, Dimbas, mike33, luuuca, MeowsterHunter, dismalbogs, axedl, Ironeyeiris, andromeda14, spidermonkey, elicemck, UnsafeHurdle, ale, ???? ??????, little.dragon, bemljj, hanabanana, PColtman, mykaelapeterson17, gowachin, Amal A., Ms.Williams, bringing-midnight, tamas, UltimateNerd00, noooo, TraÔzer, Jorgelgq, olajka, froggy20, bekkel, Ginger1885, Stannum, SeaJayBravo982, BobÌ?ek, mandudeguy111, didi&aiai, dahfizz, DragonRanger, mikko114, eonur, wp19, bjohnson97211, walker01, adi2, arooj, Lupatran, williamp55, Fedian, grabas, lotus7productions, korshe, MasterPlasmide, renandf, wearefamily, Reyuen, barjos, ColeBianchi, clarkegar, cheercoder, magnuskri, L132465, richardrobinson, robymcbobster, intersysop, Butter3646, Chlorocidic, Jimroide, Shruti Dhara Sharma, alexis.lyles, guaninedestroyer, HopelessOptimist, cheepwine, jcallej0, Eva777, coolk, Loki-74, lucasmacielbiotech, francodanussi, davidprz, HAY SCIENCE, nachoninja, duskglow, xoxo_teddy, mebrou96, Shaheen, hiroman, LagnSteel, patrickmen11, theravin, RCHoss, Kenny01294, magyar, dell188, jonny4547, Exalaber, Sam_Ravenclaw, AhmedHexa, mistymountainminer, looool, Boss360, Lvenok, LittleFRU, king063, Trans-cendance, marygill, Milkshake, bryanlim, Raleigh.Billiot, AdrienneChan, harbir123, calculo, chuckdang, Nightingale_Royale, mayrablanco97, hjgf, SophiaTate, pentester101, Lord_Crakoz, holymedecine, Heidemac01, elementnumber46, humveev, OrigamiMarie, Schiffinor, arg1215, fg, Camoes14, askro, xdperiod, Barik1931, TroyStopera, DOM31, Pinguim, tafos, Brent Schofield, alfmonc, Andrew4656, wipawadee, tweety0407, henryachan, carlosydos, IORNKING, HDBarbee, Justintoxicated, nickbooth, jp1224, HelloTheUniverse, glass130, joestudent, dbrown, Jdog, evilbiologist, Diogene, LucasAlmeida, Jai, mouseketia, jennifer24415, TruVenom, anurashrestha12, fapercin, JayFromNJ, acizkul58, couturin, ollieoe15, Jrbj2002, TitanPlayz, 12jp74, HScience, Fuzzybear, nfuente, qi4e2me1234q, blrosene, mtujamin, jbeletic, lraine, mrzwing, Byonzub, darchambault, gaminglazoorYT, Katiiie98, hickster, zundraim, EnigmaMXM, Infinite_Apollos, Didda, Folder1365, gabe.stokes, Stevan, Xavier11223, Firepuff0226, Gyrobotic, charizard325, woethe, maquique, coffevampire, Akrata111, Achraf112, stormclone80, JuliusHeinrich, cawolf, danielsundfeld, aaeeae, aaryanb, Gromtroll, jakovazor, thegamingfish, Aircondition, irimao, renocia, boi_social, silurio, DimaBiba, Lozy, Mr. Mollusk, Nofudecos, loumarylucy, ivg98, lupus_astartes, Louloudu44, Troll3, Kam, purzelbrot, Space Dolphin, paytn19, FriediFehlau, ZToRx, pokpok2710, the-pen, A churro8, Faisal, kobou, TheDoodleBanger, Dylanslegos, sivertz, katimars, orionmango, Tuberculosadeed, divisionbyzero, lmonti, SaraBot, PJ Kotze, Trystian, ViennaUCT, theeternabot, dtsaltas, Julianparengo

## Methods

### Code Availability

Code for training EternaBrain and solving puzzles with Eterna-SAP is freely available for non-commercial use at *https://github.com/EteRNAgame/EternaBrain*.

### Data Encoding

Through the Eterna puzzle-solving interface (Figure 1A-C), players can mutate an RNA molecule’s base sequence by selecting an RNA nucleotide base (A, U, G, or C) and the location on the puzzle where they would like to make the change. Players can see the ‘target’ structure – the secondary structure they are trying to achieve – and the ‘nature-mode’ (or ‘natural’) structure – the secondary structure predicted to be the minimum free energy conformation for the current sequence. When the nature-mode and target structures match, the player has solved the puzzle. In both target and nature-mode states, players can see the predicted free energy of the molecules (in kilocalories per mole). These values are routinely used by players to guide moves that make their RNA structure more stable. These energies and structures are calculated using Vienna 1.8.5 (6), which provides the default energy model in Eterna; additional tests with newer energy functions are possible in EternaBrain but are not reported here.

All information given to Eterna players was encoded before being passed into the CNN (see Table 1). Such information included the nucleotide sequence, nature-mode and target structure and pairmaps, nature-mode and target energy, and locked bases, as follows. The nucleotide bases were encoded using a standard ‘one-hot’ representation over four input layers; A, U, G, C were mapped to [1,0,0,0], [0,1,0,0], [0,0,1,0], and [0,0,0,1], respectively. Simultaneous mutations of bases were treated as separate changes, and any copying or resetting of bases sequences were encoded as [1,1,1,1] in the otherwise one-hot input for A, U, G, or C. For encoding the ‘nature-mode’ structure (minimum free energy structure predicted by Vienna 1.8.5) and target structure, dot-bracket notation was converted to one-hot representation, with unpaired bases set to zero and left- and right-paired bases set to one in three separate input layers. Another representation of the nature-mode and target structures used is structure pairmaps. Each entry in a pairmap, corresponding to a particular location in the sequence, stores the index of the base with which it forms a base pair. For example, if the base at location 1 is paired to location 10, then index 0 in the pairmap would contain the number 9, and index 9 in the pairmap would contain the number 0 (using a list starting at index 0). For training, the player’s chosen base and the location of base change were provided as labels and a standard soft-max (29) loss function computed agreement of the neural network predictions and the actual player moves. An example of the final data encoding is shown in Table 1.

### Model Construction and Evaluation

Two CNNs were built: one for predicting the RNA base, and one for predicting the location of the base change. After running several experiments on different CNN architectures, the following architecture was used for each CNN: 10 convolutional layers (with convolution size 9 in the dimension that indexes across RNA sequence position, stride 2×2, and pooling; the number of neurons in each layer were 1, 2, 4, 8, 16, 32, 64, 128, 256, 512, 1024) followed by 4 fully-connected layers (1024, 1024, 2048, 4096), a dropout rate of 0.1, a sigmoid (30) activation function, minibatch size of 100 solutions, and Adam (31) optimizer to minimize the error of the neural network. A modest hyperparameter search was carried out before settling on this architecture, as described in Supplemental Table S1. Construction and training of CNNs were carried out using Google’s TensorFlow (32) machine learning framework.

The models discussed in this work were trained on Nvidia Titan X GPUs available on Stanford’s Sherlock cluster. For puzzle playouts, the CNN initially attempted to solve the puzzle by iteratively choosing moves and updating the game state. The number of such moves was chosen to be the length of the puzzle multiplied by three. The move choice was stochastic, with probabilities of base and location based on the respective CNN output renormalized to 1.0. If this stage was not able to solve the puzzle, the SAP was applied, implementing the player strategies in Figure 2. The SAP changed specific nucleotides in the puzzle according to the player strategy and compared if the nature-mode structure of the RNA more closely matched the target structure than if the player strategy had not been implemented. If the nature-mode structure more closely matched the target structure, then the resulting nucleotide sequence was kept. (For speed, closeness of two secondary structures was defined based on the length of the largest matching subsequence of the two secondary structures written in dot-bracket notation, using the SequenceMatcher class in Python.) This process was repeated for all of the player strategies. The current implementation of SAP uses a few straightforward player strategies, favoring simplicity over computational complexity (see Figure 2).

During development of the model, we evaluated the CNN using cross-validation, training on 30,000 moves and testing on the remaining moves; test moves were pulled randomly from the data set unless noted otherwise (29). Playout tests were carried out on the 100 secondary structures of the Eterna100 benchmark (10). To ensure fair comparison to prior work (10), we did not include puzzle constraints on, e.g., minimum or maximum number of AU pairs, which arise in some of the Eterna100 puzzles when they are played by humans online.

### Descriptions of each SAP strategy

Single-action playout (SAP) consists of the application of six strategies:

1. The first step of the SAP is to correct incorrect base pairings. The algorithm traverses the entire length of the sequence, identifying any mismatches in base pairings. Any incorrect pairings will be mutated to A-U or G-C, depending on one of the bases in the mismatch.
2. Second, the algorithm will identify any base pairs flanking a hairpin loop or an internal loop and change that base pair to G-C, since G-C pairs at the end of stacks lower the energy of that subregion.
3. Third, within any internal loop, the algorithm will change the first base on either side of the loop (see Fig. 3b). This prevents mismatches with the flanking pairs of the stack.
4. Fourth, in any internal loop with two bases on either side, a U and a G are placed on both sides. This is a strategy (U-G-U-G ‘super-boost’) discovered by Eterna players which drastically lowers the free energy of the RNA molecule and takes advantage of a special parameter for tandem G•U’s in current RNA nearest-neighbor folding models.
5. Fifth, the SAP places a guanine at the beginning of any hairpin loop to lower the overall energy of the hairpin, again reflecting a special rule or trend in current RNA nearest-neighbor folding models.
6. Lastly, if the puzzle is still not folding into the desired shape, the SAP will randomly flip base pairs in target helices that are not predicted to fold properly. Eterna players often do this flipping, as an incorrect orientation could potentially be the reason why that specific area is not folding correctly.

See Figure 2 for illustrations of these six strategies.

### Puzzle-solving efficiency

The successful puzzle solution by EternaBrain-SAP would not be useful if it takes more time than experienced players to solve puzzles. Supplemental Figure S4 shows player and EternaBrain-SAP times for puzzles of varying lengths. On one hand, EternaBrain-SAP generally takes more moves (approximately 2-fold) than top Eterna players to solve most puzzles, across all puzzle lengths. However, this disadvantage is ameliorated by the speed with which EternaBrain-SAP selects its moves: on a single 2.5 GHz dual-core Intel i5 CPU, the automated method takes less total time than top players across puzzles of varying lengths. We took random samples of 50 puzzles that both EternaBrain-SAP and players solved in order to see if there was a statistically significant difference in completion time. A two-sample t-test gave a *p*-value of 0.020, indicating that EternaBrain-SAP is significantly faster than players in solving Eterna puzzles despite no specific optimization for speed at this stage.

### Author Contributions Statement

R.V.K, R.D, and B.K formulated the problem. B.K. and F.P. carried out data acquisition and data curation. R.V.K. carried out the modeling and analyses, wrote the manuscript, and generated figures. K.R.C ran validation experiments. Eterna Participants contributed problems, moves, and strategies, and review of the manuscript through online Google docs. R.D, B.K, K.R.C, and F.P edited the manuscript.

### Additional Information

The authors declare no competing interests.

