## Supplemental Figures for "EternaBrain: Automated RNA design through move sets and strategies from an Internet-scale RNA videogame"

**^‡^** Group Author: See Acknowledgments section in main text.

This Supporting Information Document contains two figures and two tables.

### Supporting Figures

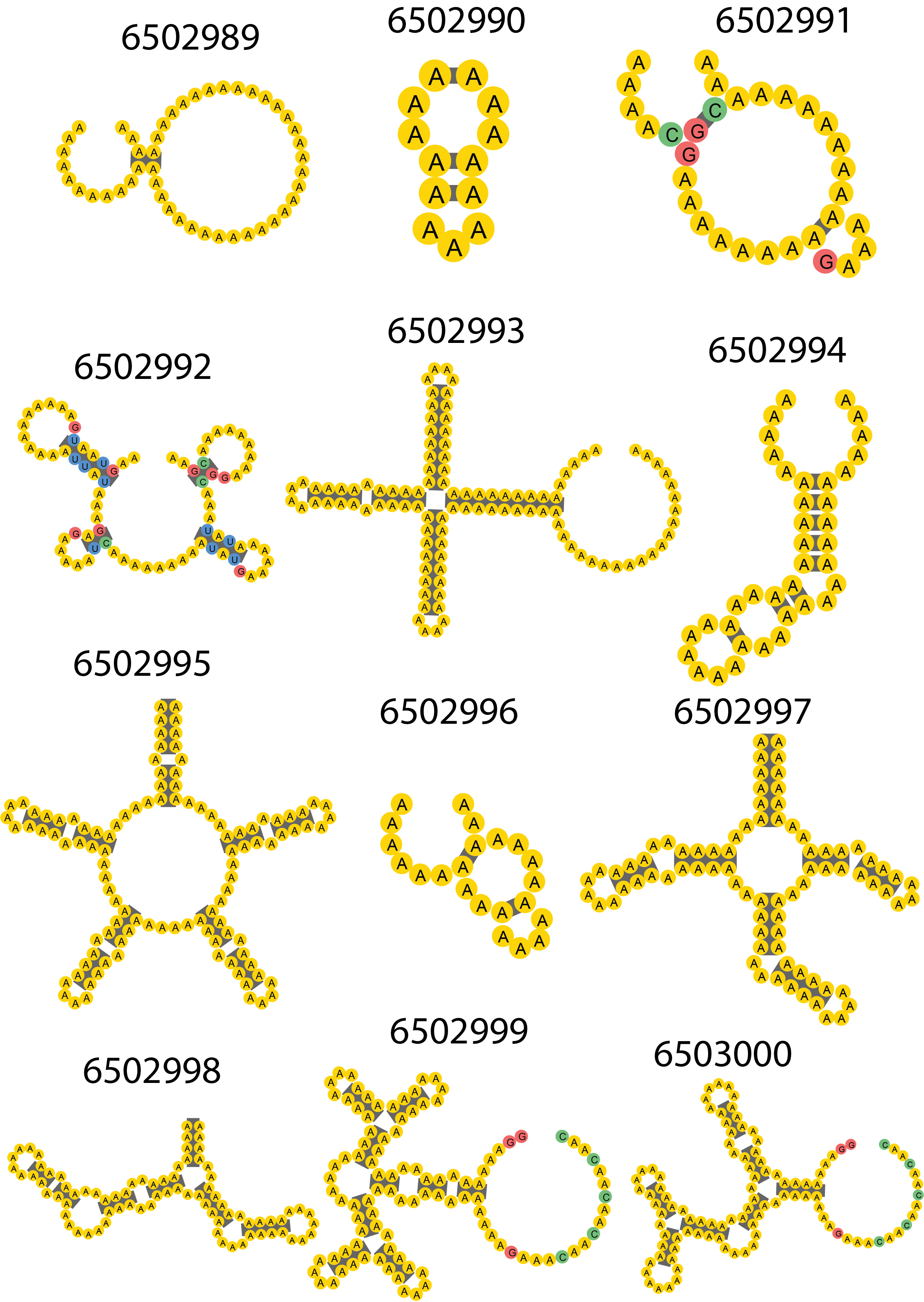

**Supporting Figure S1. The twelve puzzles used in the *eternamoves-large* dataset.** Most bases are adenine to represent the initial state of the puzzle before any mutations are made. Some bases that are not A represent “locked” bases which cannot be mutated. The 5’ end of each puzzle is at the top left, with the puzzle drawn counter-clockwise from that point.

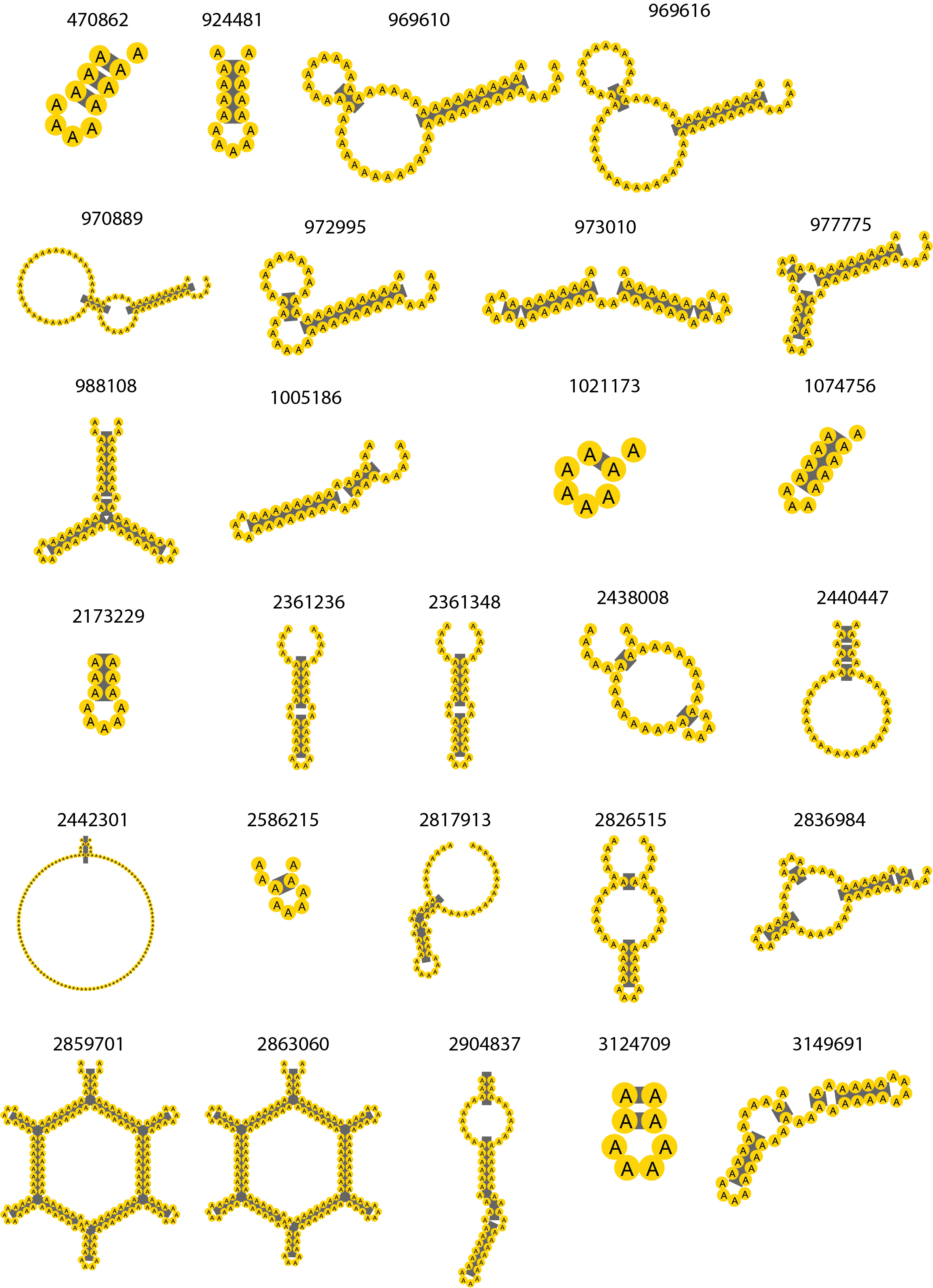

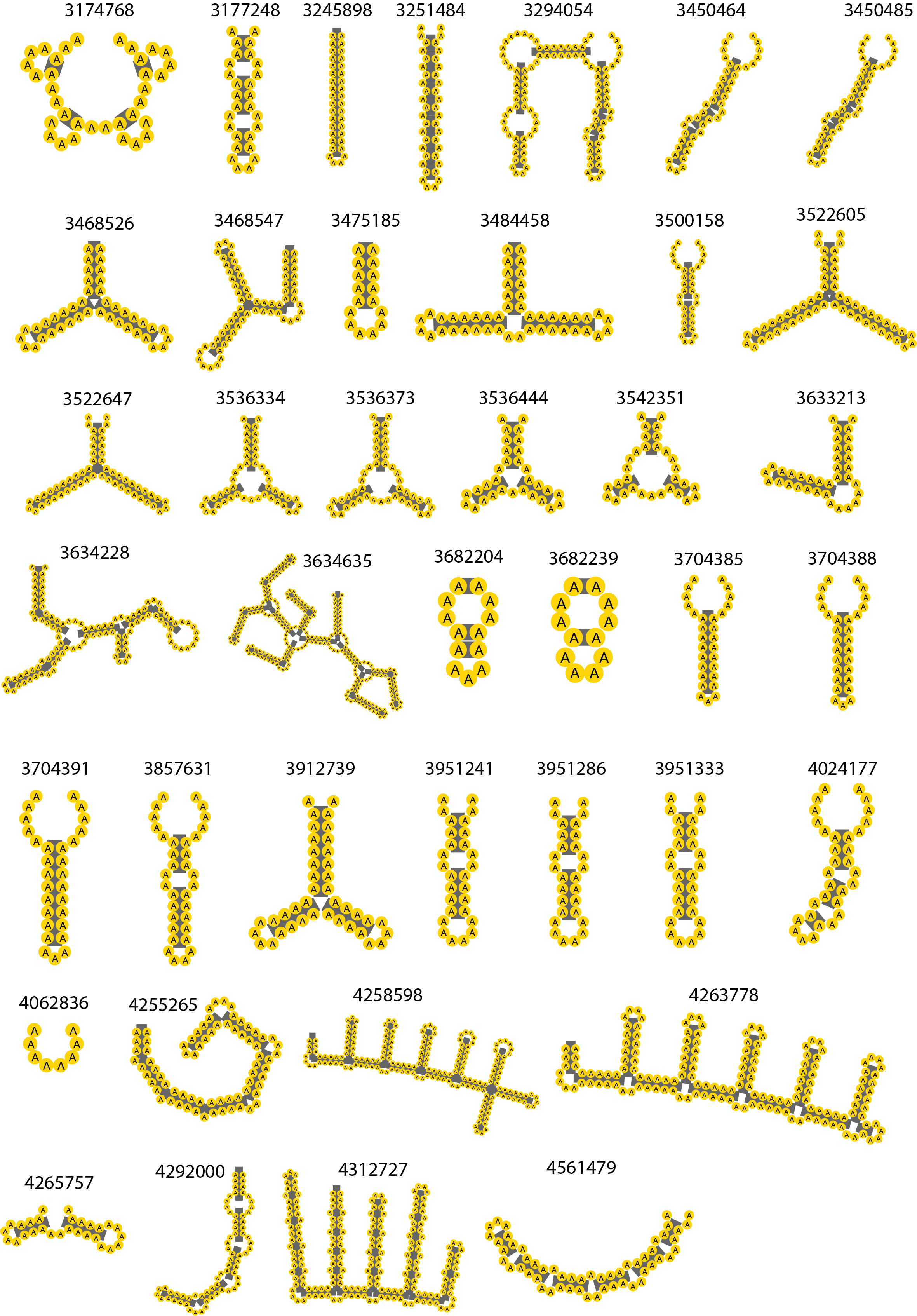

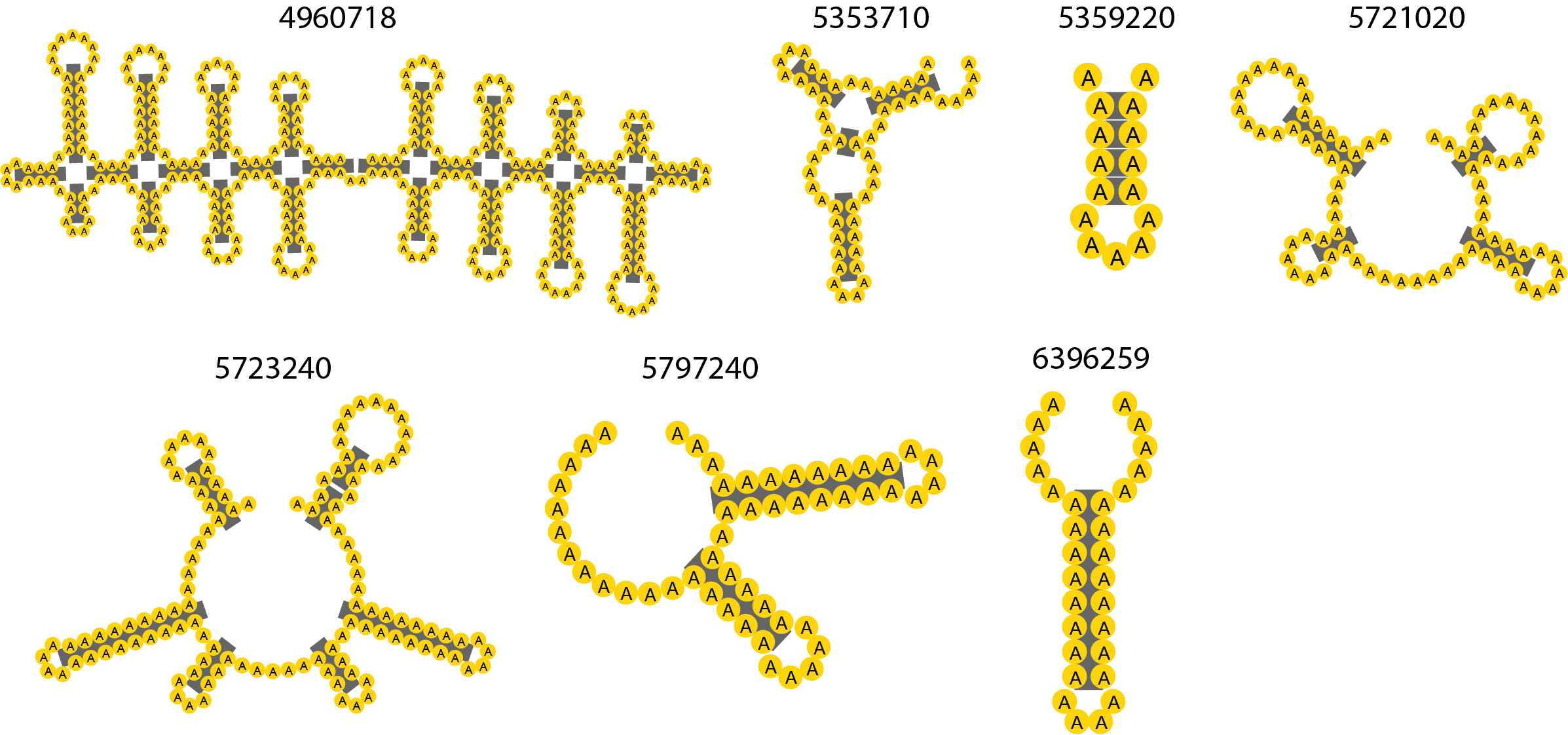

**Supporting Figure S2. (this page and previous two pages) The 78 puzzles used in the *eternamoves-select* dataset.** Some puzzle structures are repeated because the locked bases (bases which cannot be mutated) are different for the two puzzles. The 5’ end of each puzzle is at the top left, with the puzzle drawn counter-clockwise from that point.

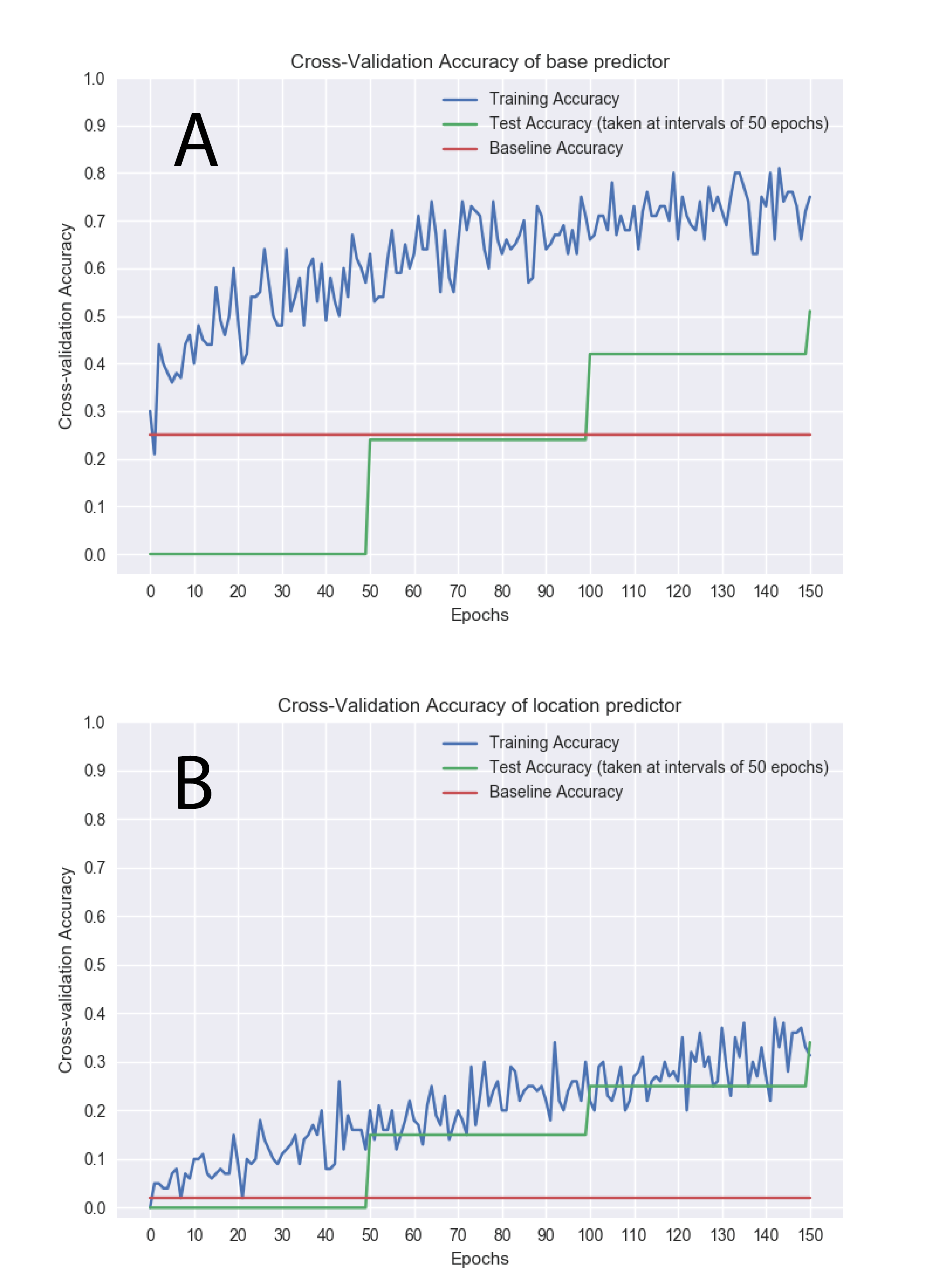

**Supporting Figure S3. Training the EternaBrain convolutional neural network on *eternamoves-select*.** (A) Cross-validation accuracy of location predictor per epoch of training. (B) Cross-validation accuracy of base predictor per epoch of training. An epoch of training is one complete pass of backpropagation, retraining, and receiving feedback of performance over all 30,447 solutions (305 mini-batches of 100 solutions each).

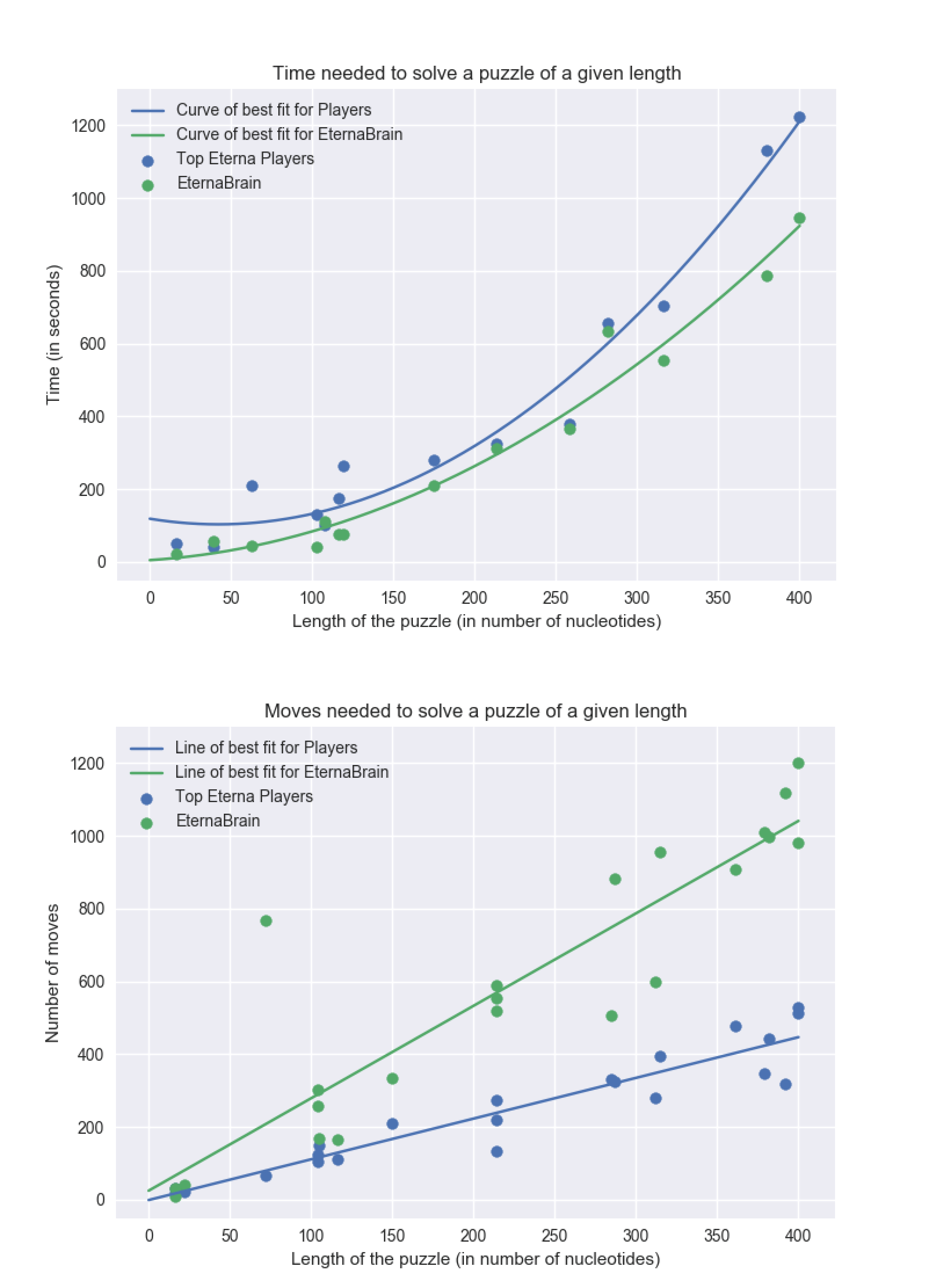

**Supporting Figure S4. Time and number of moves of EternaBrain-SAP compared to Eterna human players.** (A) Median time and (B) number of moves needed to solve Eterna100 puzzles. Puzzles that were only solvable by Eterna human players are not shown.

### Supporting Tables

**Supporting Table S1.** **Hyperparameter search for neural net architecture.**

| Model Hyperparameters^a^ | Base Train Accuracy^b^ | Base Test Accuracy^b^ | Location Train Accuracy^b^ | Location Test Accuracy^b^ |
| --- | --- | --- | --- | --- |
| 0.1 dropout, Adam, sigmoid, 10 conv layers, 4 fc layers, CNN | 0.50 | 0.34 | 0.10 | 0.021 |
| 0.0 dropout, Adam, sigmoid, 10 conv layers, 4 fc layers, CNN | 0.61 | 0.26 | 0.15 | 0.018 |
| 0.0 dropout, Adam, relu, 10 conv layers, 4 fc layers, CNN | 0.33 | 0.30 | 0.05 | 0.03 |
| 0.1 dropout, Adam, Sigmoid, 10 fc layers, DNN | 0.30 | 0.24 | 0.03 | 0.01 |

^a^ We trained 4 neural networks on the *eternamoves-large* dataset. We trained two different types of neural networks: a standard feed-forward deep neural network and a convolutional neural network. We varied a number of different hyperparameters, including dropout, number of neurons per layer, number of layers, optimizer function (Adam, SGD, etc.), activation function (relu, hyperbolic tangent, sigmoid).

^b^Accuracy for best model trained on *eternamoves-large*.

**Supporting Table S2.** **Number of solutions and moves for top players used in *eternamoves-select*.**

| **Rank**^a^ | **Usernames** | **IDs** | **Num_solutions** | **Num_moves** |
| --- | --- | --- | --- | --- |
| **0** | mat747 | 267 | 6666 | 926857 |
| **1** | akhyatt | 1623 | 183 | 11989 |
| **2** | machinelves | 2577 | 2 | 21 |
| **3** | lroppy | 2804 | 1949 | 161644 |
| **5** | tommyd | 4375 | 2723 | 204442 |
| **6** | Eli Fisker | 8627 | 8032 | 2141064 |
| **7** | starryjess | 11775 | 97 | 1707 |
| **8** | armin | 19442 | 855 | 33768 |
| **9** | Brourd | 24263 | 1018 | 244522 |
| **10** | Meechl | 26574 | 313 | 9732 |
| **13** | c-quence | 29762 | 589 | 25905 |
| **14** | stevetclark | 32487 | 241 | 7494 |
| **15** | jandersonlee | 32627 | 2277 | 106044 |
| **16** | choo | 34334 | 232 | 6255 |
| **17** | drake178 | 34596 | 158 | 4194 |
| **19** | hoglahoo | 36921 | 1632 | 107248 |
| **20** | fluffy3 | 39309 | 56 | 1717 |
| **23** | 77Tennifry | 42101 | 2 | 26 |
| **25** | JR | 42833 | 4706 | 618972 |
| **27** | Jsci | 43776 | 2144 | 284749 |
| **29** | wawan151 | 44191 | 255 | 39711 |
| **30** | macclark52 | 44631 | 22 | 1164 |
| **31** | hotcreek | 46281 | 6 | 118 |
| **32** | jnicol | 48166 | 241 | 11290 |
| **33** | Jieux | 48170 | 4216 | 698491 |
| **34** | Tesla'sDisciple | 50065 | 1588 | 86498 |
| **35** | Malcolm | 52361 | 5478 | 740470 |
| **37** | RedSpah | 55082 | 3 | 83 |
| **39** | janetmason | 56579 | 435 | 14565 |
| **40** | garydfisher | 57654 | 797 | 41876 |
| **41** | Omei | 57675 | 6136 | 426299 |
| **42** | wateronthemoon | 57743 | 882 | 38259 |
| **43** | cataway | 57874 | 63 | 1943 |
| **44** | Hyphema | 58224 | 57 | 9943 |
| **45** | rnjensen45 | 60391 | 10 | 260 |
| **47** | theravin | 61244 | 2 | 7 |
| **49** | AndrewKae | 64544 | 1135 | 161290 |
| **52** | Gres | 77611 | 498 | 52423 |
| **53** | ulfang | 78801 | 866 | 51782 |
| **55** | Zanna | 87216 | 1309 | 49302 |
| **57** | salish99 | 132701 | 67 | 657 |
| **58** | biggestlegoheroicafanever | 133043 | 415 | 31166 |
| **59** | wookietank | 136909 | 576 | 17795 |
| **60** | VirgieP3 | 143547 | 106 | 2889 |
| **62** | Mayanne | 152013 | 379 | 9998 |
| **63** | benrh | 179978 | 6510 | 565822 |
| **65** | skyblue | 207838 | 2061 | 188398 |
| **66** | Marzena11 | 209752 | 240 | 9534 |
| **67** | Eized | 216078 | 321 | 19185 |
| **68** | dl2007 | 223393 | 1800 | 1773005 |
| **69** | joy45 | 225023 | 1062 | 49752 |
| **70** | DeNa | 231977 | 6991 | 949354 |
| **71** | cynwulf28 | 233206 | 5798 | 545017 |

^a^ Participant rank is based on total number of puzzles solved on Eterna at the time of the study, and not just puzzles in *eternamoves-select*; not all participants solved puzzles chosen for *eternamoves-select*.
