## Supplemental Table S1 for "EternaBrain: Automated RNA design through move sets and strategies from an Internet-scale RNA videogame"

**Supplemental Table S2.** **Hyperparameter search for neural net architecture.** We trained 4 neural networks on the *eternamoves-large* dataset. We trained two different types of neural networks: a standard feed-forward deep neural network and a convolutional neural network. We varied a number of different hyperparameters, including dropout, number of neurons per layer, number of layers, optimizer function (Adam, SGD, etc.), activation function (relu, hyperbolic tangent, sigmoid).

Accuracy for best model trained on *eternamoves-large*:

| Model Hyperparameters | Base Train Accuracy | Base Test Accuracy | Location Train Accuracy | Location Test Accuracy |
| --- | --- | --- | --- | --- |
| 0.1 dropout, Adam, sigmoid, 10 conv layers, 4 fc layers, CNN | 0.50 | 0.34 | 0.10 | 0.021 |
| 0.0 dropout, Adam, sigmoid, 10 conv layers, 4 fc layers, CNN | 0.61 | 0.26 | 0.15 | 0.018 |
| 0.0 dropout, Adam, relu, 10 conv layers, 4 fc layers, CNN | 0.33 | 0.30 | 0.05 | 0.03 |
| 0.1 dropout, Adam, Sigmoid, 10 fc layers, DNN | 0.30 | 0.24 | 0.03 | 0.01 |
